# Non-invasive genomic sampling uncovers novel connectivities and origins of confiscated gorillas

**DOI:** 10.64898/2026.02.03.701083

**Authors:** Irune Ruiz-Gartzia, Harvinder Pawar, Marina Alvarez-Estape, Joseph D. Orkin, Pol Alentorn-Moron, Sandra Ruibal-Puertas, Harrison J. Ostridge, Claudia Fontsere, Sebastian Cuadros-Espinoza, Kirsten Gilardi, Julius Nziza, Richard Muvunyi, Ulrich Maloueki, Damien Caillaud, Neetha Iyer, Urban Ngobobo-As-Ibungu, Escobar Binyinyi, Etienne François Akomo-Okoue, Augustin K. Basabose, Kamungu Sebulimbwa, Alfred Ngomanda, Juichi Yamagiwa, Miho Inoue-Murayama, Yuji Takenoshita, Shiho Fujita, Barbora Pafco, Terence Fuh, Nikki Tagg, Donald Mbohli, Emmanuel Ayuk Ayimisin, Emma Bailey, Mattia Bessone, Tobias Deschner, Paula Dieguez, Emmanuel Dilambaka, Andrew Dunn, Anne-Celine Granjon, Josephine Head, Veerle Hermans, Inaoyom Imong, Kathryn J. Jeffery, Kevin Lee, Giovanna Maretti, David Morgan, Emily Neil, Christopher Orbell, Martha M. Robbins, Crickette Sanz, Nadege Wange Njomen, Jacob Willie, Barthélémy Ngoubangoye, Larson Boundenga, Franck Prugnolle, Olivia Cheronet, Ron Pinhasi, Ivo Glynne Gut, Marta Gut, Karine A. Viaud-Martinez, Virginie Rougeron, Bethan Morgan, Christophe Boesch, Hjalmar S. Kühl, Mimi Arandjelovic, Eiji Inoue, Katerina Guschanski, Klara Petrzelkova, Martine Peeters, Magdalena Bermejo, Germán Illera Basas, Torsten Bohm, Tierra Smiley Evans, Javier Prado-Martinez, Martin Kuhlwilm, Esther Lizano, Tomas Marques-Bonet

**Affiliations:** IBE, Institute of Evolutionary Biology (UPF-CSIC), Department of Medicine and Life Sciences, Universitat Pompeu Fabra, Barcelona, Spain; Department of Biology, University of Copenhagen, Copenhagen, Denmark; Département d’anthropologie, Université de Montréal, Montréal, Québec, Canada; Département de sciences biologiques, Université de Montréal, Montréal, Québec, Canada; UCL Genetics Institute, Department of Genetics, Evolution and Environment, University College London, London, UK; Center for Evolutionary Hologenomics, The Globe Institute, University of Copenhagen, Øster Farimagsgade 5A, 1352 Copenhagen, Denmark; Gorilla Doctors, Karen C. Drayer Wildlife Health Center, One Health Institute, University of California Davis, School of Veterinary Medicine, Davis, CA, USA; Rwanda Development Board, Kigali, Rwanda; Faculty of Sciences and Technologies, Life Sciences, Kinshasa University, Kinshasa, Republic Democratic of the Congo; African Parks, Odzala-Kokoua National Park, Brazzaville, Republic of Congo; Department of Anthropology, University of California, Davis, CA, USA; The Dian Fossey Gorilla Fund International, Kinshasa, DRC; Institut de Recherche en Ecologie Tropicale (IRET/CENAREST). B.P. 13354 Libreville, Gabon; Centre de Recherche en Sciences Naturelles de Lwiro. DS Bukavu, RD Congo; Research Institute for Humanity and Nature; Wildlife Research Center of Kyoto University, Kyoto 606-8203, Japan; Department of Zoology, Faculty of Science. Okayama University of Science; Center for General Education, Kagoshima University, Kagoshima, Japan; Institute of Vertebrate Biology, Czech Academy of Sciences, Kvetna 8, 603 00 Brno, Czech Republic; WWF DRC country programme, 4630, Avenue de la science, Gombe, Kinshasa, DRC; World Wildlife Fund, Avenue des Martyrs, Bangui, Central African Republic; KMDA, Centre for Research and Conservation, Royal Zoological Society of Antwerp, Koningin Astridplein 20-26, B-2018 Antwerp, Belgium; Association pour la Protection des Grands Singes (APGS), Nkol-Eton Yaounde - Cameroon; Max Planck Institute for Evolutionary Anthropology, Leizpig, Germany; University of Konstanz, Centre for the Advanced Study of Collective Behaviour, Universitätsstraße 10, 78464, Konstanz, Germany; Department of Animal Societies, Max Planck Institute of Animal Behavior, Bücklestrasse 5a, 78467, Konstanz, Germany; Institute of Cognitive Science, University of Osnabrück, Artilleriestrasse 34, 49076 Osnabrück, Germany; Senckenberg Museum für Naturkunde Görlitz, Am Museum 1, 02826 Görlitz, Germany; Wildlife Conservation Society (WCS), 2300 Southern Boulevard. Bronx, New York 10460, USA; German Centre for Integrative Biodiversity Research (iDiv), Leipzig, Germany; Leipzig University, Ritterstraße 26, 04109 Leipzig, Germany; Arcus Foundation, Cambridge, UK; School of Natural Sciences, University of Stirling, UK; AMAP, University of Montpellier, Montpellier, France; School of Human Evolution and Social Change, Arizona State University, 900 Cady Mall, Tempe, AZ 85287 Arizona State University, PO Box 872402, Tempe, AZ 85287-2402 USA; Institute of Human Origins. PO Box 878404. Tempe, AZ 85287-8404; Ngogo Chimpanzee Project, Makerere University Biological Field Station, PO Box 409, Fort Portal, Uganda; Lincoln Park Zoo, Lester E Fisher Center for the Study and Conservation of Apes, 2001 N Clark St, Chicago, IL 60614, USA; Panthera, 8 W 40TH ST, New York, NY 10018, USA; Department of Primate Behavior and Evolution, Max Planck Institute for Evolutionary Anthropology, Leizpig, Germany; Washington University in Saint Louis, Department of Anthropology, One Brookings Drive, St. Louis, MO 63130, USA; Congo Program, Wildlife Conservation Society, 151 Avenue Charles de Gaulle, Brazzaville, Republic of Congo; WWF Cameroon Country Office, BP6776; Yaoundé, Cameroon; Terrestrial Ecology Unit (TEREC), Department of Biology, Ghent University (UGent), K.L. Ledeganckstraat 35, 9000 Ghent, Belgium; Centre de Primatologie, Centre Interdisciplinaire de Recherches Médicales de Franceville (CIRMF), Franceville, BP769, Gabon; Unité de Recherche en Ecologie de la Santé (URES), Centre Interdisciplinaire de Recherches Médicales de Franceville (CIRMF), Franceville, BP769, Gabon; International Research Laboratory, REHABS, CNRS-Université Lyon 1-NMU, George Campus, George, South Africa; Sustainability Research Unit, Nelson Mandela University, George Campus, George, South Africa; Department of Evolutionary Anthropology, University of Vienna, Vienna, Austria; Human Evolution and Archaeological Sciences (HEAS), University of Vienna, Wien, Austria; Centro Nacional de Análisis Genómico (CNAG), Baldiri Reixac 4, 08028 Barcelona, Spain; Universitat de Barcelona (UB), Barcelona, Spain; Illumina Laboratory Services, Illumina. Inc, San Diego, CA USA; San Diego Zoo Wildlife Alliance, San Diego, California, USA; Wild Chimpanzee Foundation; International Institute Zittau, Technische Universität Dresden, Markt 23, 02763 Zittau, Germany; Department of Biology, Faculty of Science, Toho University, Chiba, 274-8510, Japan; Institute of Ecology and Evolution, School of Biological Sciences, University of Edinburgh, Edinburgh, UK; Animal Ecology, Department of Ecology and Genetics, Uppsala, University, Uppsala, Sweden; Institute of Parasitology, Biology Centre, Czech Academy of Sciences, Branisovska 31, 370 05 Ceske Budejovice; TransVIHMI, Université de Montpellier, INSERM, IRD, Montpellier, France; Department of Ecology and Environmental Sciences, University of Barcelona, 08028 Barcelona, Spain; Sabina Plattner African Charities (SPAC) NPC, Aria North Wharf, 42 Hans Strijdom Avenue, Foreshore, 8001, Cape Town, South Africa; Department of Integrative Biology & Infectious Diseases and Vaccinology Division, School of Public Health, University of California Berkeley; Institut Català de Paleontologia Miquel Crusafont (ICP-CERCA), Universitat Autònoma de Barcelona, Edifici ICTA-ICP, Cerdanyola del Vallès, Barcelona, Spain; Unidad de Paleobiología, ICP-CERCA, Unidad Asociada al CSIC por el IBE UPF-CSIC, Barcelona, Spain; Catalan Institution of Research and Advanced Studies (ICREA), Barcelona, Spain

**Keywords:** wild gorillas, non-invasive, population dynamics, connectivity

## Abstract

**Background:** Gorillas are a group of African great apes with two species and four subspecies that are currently critically endangered or endangered. Previous studies that analysed the genetics of wild gorillas from non-invasive samples, such as faeces or hair, analysed short mitochondrial or nuclear markers, which may not reflect the wider nuclear genome. Recent technical advances in target capture hybridisation, enrich the endogenous DNA content of non-invasive samples, allowing contiguous genomic regions to be sequenced.

**Results:** Here, we generated georeferenced genetic data from faecal and hair samples of 280 wild gorillas, sampled from three of the four gorilla subspecies, across large parts of their present-day distributions.

With this expanded representation of gorilla genetic diversity in the wild, we detected three population clusters in western lowland gorillas, with the Sangha River and its affluents acting as significant barriers to gene flow.

We reconstructed patterns of past population connectivity between western lowland gorillas in the north-eastern distribution range and Cross River gorillas, which may have been facilitated by a migration corridor also used by the Central and Nigeria-Cameroon chimpanzee subspecies.

Finally, we predicted the geographic origins of wild-born gorillas, achieving a mean prediction error of 65 km, with a population-level resolution for mountain gorillas and some populations of western lowland gorillas.

**Conclusion:** Our work characterises fine-scale population structure in western lowland gorillas, which will be informative for future conservation strategies. This proof of concept in predicting geographic locations of wild gorillas, will be useful for future applications to geolocalise trafficked or rescued gorillas.

## Introduction

Two species of gorilla are recognised, each of which has two subspecies (Grubb et al. 2003). Eastern gorillas consist of eastern lowland gorillas (*Gorilla beringei graueri)* and mountain gorillas (*Gorilla beringei beringei*). Western gorillas consist of Cross River gorillas (*Gorilla gorilla diehli*) and western lowland gorillas (*Gorilla gorilla gorilla*). Anthropogenic factors, such as habitat loss, habitat degradation, land-use change and poaching have impacted gorillas. Infectious disease transmission is also a significant risk to wild gorilla populations, particularly with regards to Ebola (Fontsere et al. 2021a; Bermejo et al. 2006; Caillaud et al. 2006). Cross River gorillas, eastern lowland gorillas and western lowland gorillas are all currently critically endangered in the IUCN Red List of Threatened Species (Maisels et al. 2018; Bergl et al. 2016; Plumptre et al. 2016; Estrada et al. 2017; Imong et al. 2013). Mountain gorillas were previously listed as critically endangered, but have been downgraded to endangered following successful conservation efforts (Hickey et al. 2018; Robbins et al. 2011).

Genetic data on gorillas has uncovered aspects of their genetic diversity, population structure and demographic history (Alvarez-Estape et al. 2023; Pawar et al. 2023; McManus et al. 2015; Fünfstück and Vigilant 2015; Fünfstück et al. 2014; Xue et al. 2015; Prado-Martinez et al. 2013a; van der Valk et al. 2024). Among the subspecies, western lowland gorillas, exhibit the highest levels of genetic diversity and highest estimated long-term effective population size (Strindberg et al. 2018; Maisels et al. 2018; Prado-Martinez et al. 2013a; Scally et al. 2012; Scally et al. 2013; Thalmann et al. 2007). Western lowland gorillas also occupy the largest distribution range of the subspecies (Maisels et al. 2018), and previous work has suggested that geographical features such as major rivers may demarcate their genetic structure (Fünfstück et al. 2014; Anthony et al. 2007).

However, the genomic studies were mostly based on limited numbers of high-quality whole genomes generated from blood or tissue samples of gorillas (Prado-Martinez et al. 2013a; Xue et al. 2015; Pawar et al. 2023; Kuderna et al. 2023). Ethical and logistical concerns limit invasive sampling of blood and tissue of endangered wild populations, such as gorillas. Where non-invasively collected samples, such as faeces or hair, have been sampled from wild gorillas, analysis was typically restricted to the mitochondrial control region or nuclear microsatellites (Fünfstück and Vigilant 2015; Fünfstück et al. 2014; Anthony et al. 2007), which may not reflect the wider nuclear genome. Here we aim to fill this gap by taking advantage of technical advances in target capture hybridisation that enrich the endogenous DNA content from faeces, allowing contiguous genomic regions to be sequenced (Fontsere et al. 2021b; Lester et al. 2021; White et al. 2019; Hernandez-Rodriguez et al. 2018; Perry et al. 2010). With the aim to characterise population structure and connectivity of wild gorillas, we non-invasively sampled faecal and hair samples from 280 wild gorillas, across the distribution ranges of eastern lowland gorillas, mountain gorillas and western lowland gorillas. This included sampling sites of western lowland gorillas for which genomic data had not been previously generated. This was possible due to extensive collaboration with researchers in the field. We applied the target-capture hybridisation protocol developed for use in chimpanzee faecal samples (Fontsere et al. 2022) for the first time to wild gorillas, to capture chromosome 21 and exomic regions. With this increased genomic representation of wild gorilla diversity, we aimed to characterise fine-scale population structure in western lowland gorillas, and to corroborate whether this is impacted by major rivers. We also generated whole genomes for 12 rescued western lowland gorillas sampled with Flinders Technology Associates (FTA) cards®, and 11 eastern and western gorilla museum specimens. By integrating genetic data from the georeferenced non-invasively collected samples, FTA cards, museum specimens and previously published gorilla genomes, we aimed to assess gorilla diversity across space and across sample types.

## Results

### Genomic data from non-invasively collected gorilla samples

In this study, we have non-invasively sampled a total of 280 gorillas from faeces (N=265) and hairs (N=15), and recorded their geographical coordinates (Table 3 Supplementary). Given the nature of non-invasively collected samples, we performed target capture hybridisation of chromosome 21 and the exome, following the methods developed on wild chimpanzee faecal samples (Fontsere et al. 2022; Fontsere et al. 2021b; White et al. 2019). In addition, we sequenced whole genomes from 12 FTA blood-preserved samples from rescued western lowland gorillas and 11 gorilla specimens from museum collections (Table 3 Supplementary). We performed an in-depth analysis of capture performance for both chromosome 21 and exome. Capture performance was assessed based on library complexity, sensitivity, specificity and enrichment factor metrics (See Supplementary section 1). Enrichment factor, library complexity and capture specificity were similar between chr21 and exome captures (Supplementary Figure 12, Supplementary Figure 15 and Supplementary Figure 18). We note the exome regions of chromosome X and chromosomes 2-9, 17-22 were successfully captured; whereas in chromosomes 1, 10-16 few reads were present as a consequence of incomplete capture due to technical reasons (see Supplementary section 2).

As a result of our sequencing effort, we generated sequencing data to a mean coverage in the target space for chromosome 21 of 34.6X for faeces, 6.65X for hairs, and for the exome 16.7X for faeces and 4.43X for hairs. Whole genome data was generated at 1.78X for the FTA cards and 2X for museum samples (Supplementary Figure 22).

We performed thorough quality control, excluding samples with <0.5X coverage in the target space and more than 1% of human contamination (Kuhlwilm et al. 2021) (Supplementary Figure 23). After integrating our newly sequenced samples with previously published wild gorilla data, we filtered for relatedness to exclude self and first-degree relatives, following the approach of Fontsere et al. (2022) (Supplementary relatedness figures 27-31). The combined dataset included whole genomes subset to chromosome 21 from Prado-Martinez et al. (2013a), Prado-Martinez et al. (2013b), Xue et al. (2015), Pawar et al. (2023), Alvarez-Estape et al. (2023), Kuderna et al. (2023) and chromosome 21 of 3 gorilla samples from Fontsere et al. (2022).

After quality control and filtering, we analysed 240 newly sequenced unique gorillas from 23 sampling sites across the entire gorilla distribution range (Figure 1A). Including the previously published data, our final dataset comprises a total of 294 unique gorillas from 33 sampling sites (Table 1 Supplementary).

**Figure 1.**
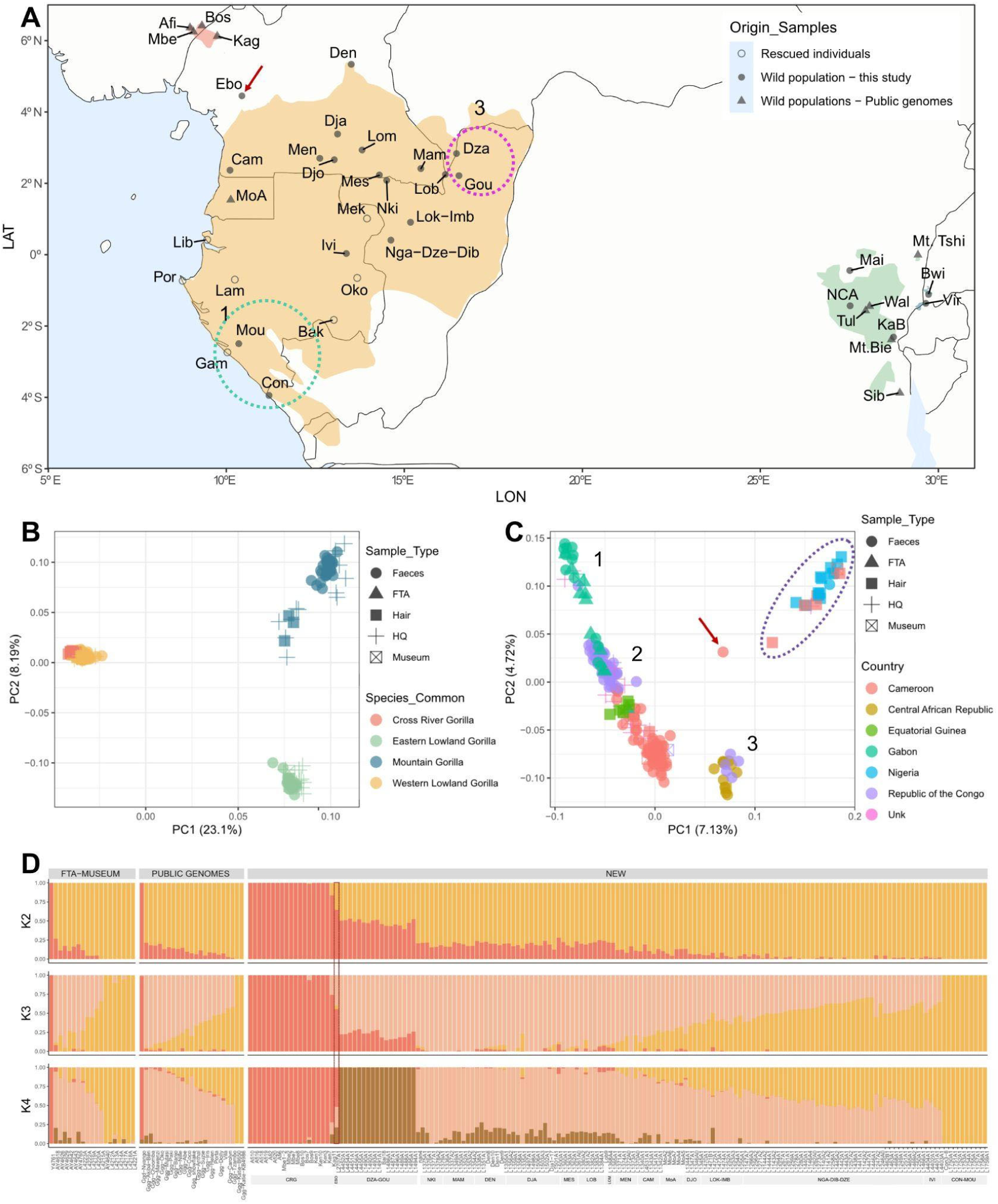
A) Map of the distribution ranges of the four gorilla subspecies. Plotted are the sampling locations for the newly sequenced non-invasively collected gorilla samples (filled circles), the recorded locations of the FTA card samples from rescued gorillas (empty circles) and the sampling locations for previously published Cross River gorillas (Alvarez-Estape et al. 2023) and eastern lowland gorillas (Xue et al. 2015; Pawar et al. 2023) (filled triangles), for which we extracted chromosome 21 and the exome. Distribution ranges are coloured as: red Cross River gorillas, yellow western lowland gorillas, green eastern lowland gorillas and blue mountain gorillas. Each sampling site is labelled with an identifier, the full names are given in Tables 1 and 3 Supplementary. The labelled circles 1 (in green) and 3 (in pink) correspond to population clusters 1 and 3 of PCA in 1C. B) PCA of Cross River (red), eastern lowland (green), western lowland (orange) and mountain (blue) gorillas for the final dataset, of chromosome 21 including newly sequenced and previously published samples. C) Principal Component Analysis of western gorillas (western lowland gorillas and Cross River gorillas). Samples are coloured by their country of origin. Different sample types are plotted with different shapes (faecal samples are circles, FTA cards are triangles, hair samples are squares, high-quality blood samples are crosses, and museum samples are squares with a cross inside). The purple dotted circle indicates the Cross River gorilla subspecies cluster. Numbers 1 to 3 indicate the three main WLG population clusters detected. D) Admixture analysis for K=2-4 of western gorilla samples. Each column represents a sample. We group samples by sample type, into 1) the newly sequenced FTA cards and museum samples, 2) public genomes (previously published samples subset to chromosome, and 3) the newly sequenced non-invasively collected samples. In panels A and C we highlight the Ebo sampling site with a red arrow, and in panel D with a red rectangle.

We find that in a principal component analysis (PCA) samples cluster according to the four gorilla subspecies (Figure 1B), regardless of sample type (faeces, hair, museum, FTA card or high-coverage blood sample). We observe that PC1 differentiates the gorilla species (eastern and western), and PC2 the eastern gorillas (eastern lowland gorilla and mountain gorilla) (Figure 1B), which is consistent with previous studies (Prado-Martinez et al. 2013a, Xue et al. 2015, Pawar et al. 2023). This divergence between eastern and western gorillas is also seen in admixture analysis, where the best inferred K, K=2 separated the two species (Supplementary Figure 47) (see Supplementary section 3).

### Genetic diversity of gorillas differs between autosomes and chromosome X

With the coding regions of wild gorillas, we characterised differences in genetic diversity in the autosomes and chromosome X. Gorillas have male heterogamety (XY) and so theory predicts the effective population size (Ne) of chromosome X relative to an autosome should be ¾. Genetic diversity can be used as a proxy for Ne and here we used heterozygosity to compare the exome of chromosome X to chromosome 4 (the autosome with a comparable sized exome target space, chromosome X = 1,257,695bp, chromosome 4 = 1,316,883bp) in wild gorilla females. We observe reduced diversity in chromosome X relative to the autosome (Figure 2), which agrees with theoretical predictions (Schaffner 2004). We also see that the regression slopes of the relationship between heterozygosity in the coding regions of chromosome 4 and chromosome X differ between the subspecies (Figure 2), with only eastern lowland and western lowland gorillas being similar.

**Figure 2.**
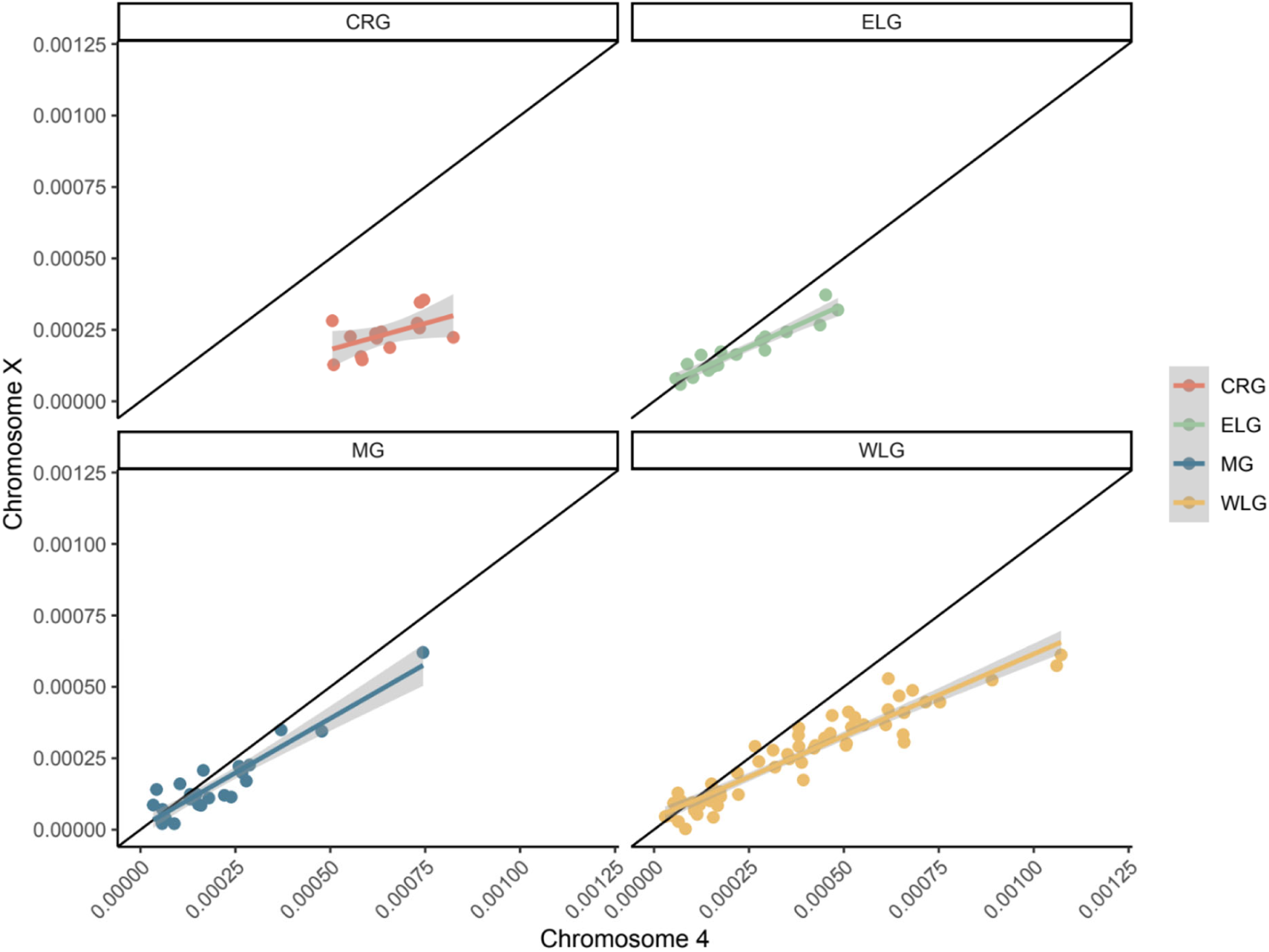
Comparison of heterozygosity in the exome of chromosome X and chromosome 4, the autosome with a comparable sized exome target space per subspecies of gorilla. In each panel y=x is plotted in black. Regression lines were calculated by fitting a linear model to the data for each subspecies.

### Population structure and connectivity in gorillas

Genetic data for chromosome 21 from 17 sampling sites and 158 samples of western lowland gorillas are reported, allowing a more fine-grained evaluation of their population structure. In PCA of western lowland gorillas, we identify three clusters (1-3 numbers indicate each of the detected clusters in Figure 1C; Supplementary Figure 37 and Supplementary Figure 38), consisting of gorillas in the 1) southern part of the western lowland distribution range (Moukalaba-Doudou National Park (Mou) in southern Gabon and Conkouati National Park in the Republic of Congo); 2) central and northern part of the western lowland distribution range (northern Gabon, Cameroon, Guinea Equatorial and the Odzala-Kokoua National Park in the Republic of Congo); 3) north-east extreme of the western lowland distribution range (Dzanga-Ndoki National Park (Dza) in the Central African Republic, and Goualougo (Gou) in the Nouabalé-Ndoki National Park in the Republic of Congo). In admixture analysis, the best inferred K is K=3 where Dza and Gou form a distinct cluster, and the rest of the western lowland populations exhibit a genetic gradient between Gabon and Cameroon (Figure 1D). For this analysis, we retained one gorilla from the Ebo Forest (Ebo) sampling site, despite being above our filtering threshold of 1% human contamination (1.53%), due to its geographic uniqueness. The unique Ebo sample showed admixed patterns between western lowland and Cross River gorillas (Figures 1C,1D). When we perform PCA within each western subspecies, the Ebo Forest sample does not cluster with any of the distinct population clusters (Supplementary Figures 37,40).

We observed low genetic differentiation, as indicated by FST, between the western lowland gorilla sampling sites from Cameroon, which are in population cluster 2 of PCA (Lobéké (Lob), Nki, Messok (Mes), Mambele (Mam), Deng Deng (Den), and La Belgique (Bel)). Among western lowland gorillas, we observed the highest genetic differentiation when comparing gorillas at Dza against any of the gorillas in the Odzala-Kokoua National Park [which consists of the sampling sites Naga-Diba-Dzébé (Nga-Dib-Dze) and Lokoué-Imbalanga (Lok-Imb)], and gorillas at Mou, in southern Gabon (Supplementary Figure 71). The FST values for these comparisons of western lowland populations are similar to the FST estimated between Bwindi and Virunga mountain gorilla populations (Supplementary Table 4). In addition, gorillas in Odzala-Kokoua National Park (*i.e*., those in Nga-Dib-Dze and Lok-Imb), and gorillas in Mou showed high genetic differentiation to Cross River gorillas.

To further assess shared genetic ancestry across gorillas, we applied the method admixfrog (Peter 2020), which allowed us to obtain fragments of chromosome 21 in each individual which are close to a reference panel from whole genome data. Fragments of shared ancestry detected with admixfrog are observed within each gorilla species to some extent (*i.e.*, sharing between eastern lowland gorillas and mountain gorillas; and sharing between Cross River gorillas and western lowland gorillas) (Figure 3A) (see Supplementary section 4). This is in agreement with previous work analysing allele sharing (Scally et al. 2012, Prado et al. 2013, Xue et al. 2015, Pawar et al. 2023, Alvarez-Estape et al. 2023). However, this is the first time the introgressed fragments between the subspecies are identified in this number of wild gorillas. Moreover, the denser sampling design here allows a more fine-grained view of shared ancestry, particularly among western gorillas. For example, we see high proportions of Cross River gorilla fragments among the western lowland gorillas located in the north-eastern distribution, corresponding to the Dza and Gou sampling sites, where 35.9% of fragments detected come from Cross River gorilla ancestry. Whereas western lowland gorillas in the south-eastern range of their distribution (such as the sampling sites, Lok-Imb and Nga-Dib-Dze) appear almost depleted for Cross River ancestry (1.66%) (Figure 3A & Figure 3B). Looking at the tract lengths, the western lowland gorillas at Dza and Gou had the longest Cross River gorilla ancestry tracts, whereas the gorillas at Lok-Imb and Nga-Dib-Dze had the shortest tracts (Supplementary Figure 84). For this analysis, the Ebo gorilla appears to show a mixture of Cross River and western lowland gorilla ancestry (Figure 3A), confirming results from principal component analyses and admixture. The length of the tracts detected for the Ebo gorilla were similar to the tracts detected in gorillas at Dza and Gou sampling sites.

**Figure 3.**
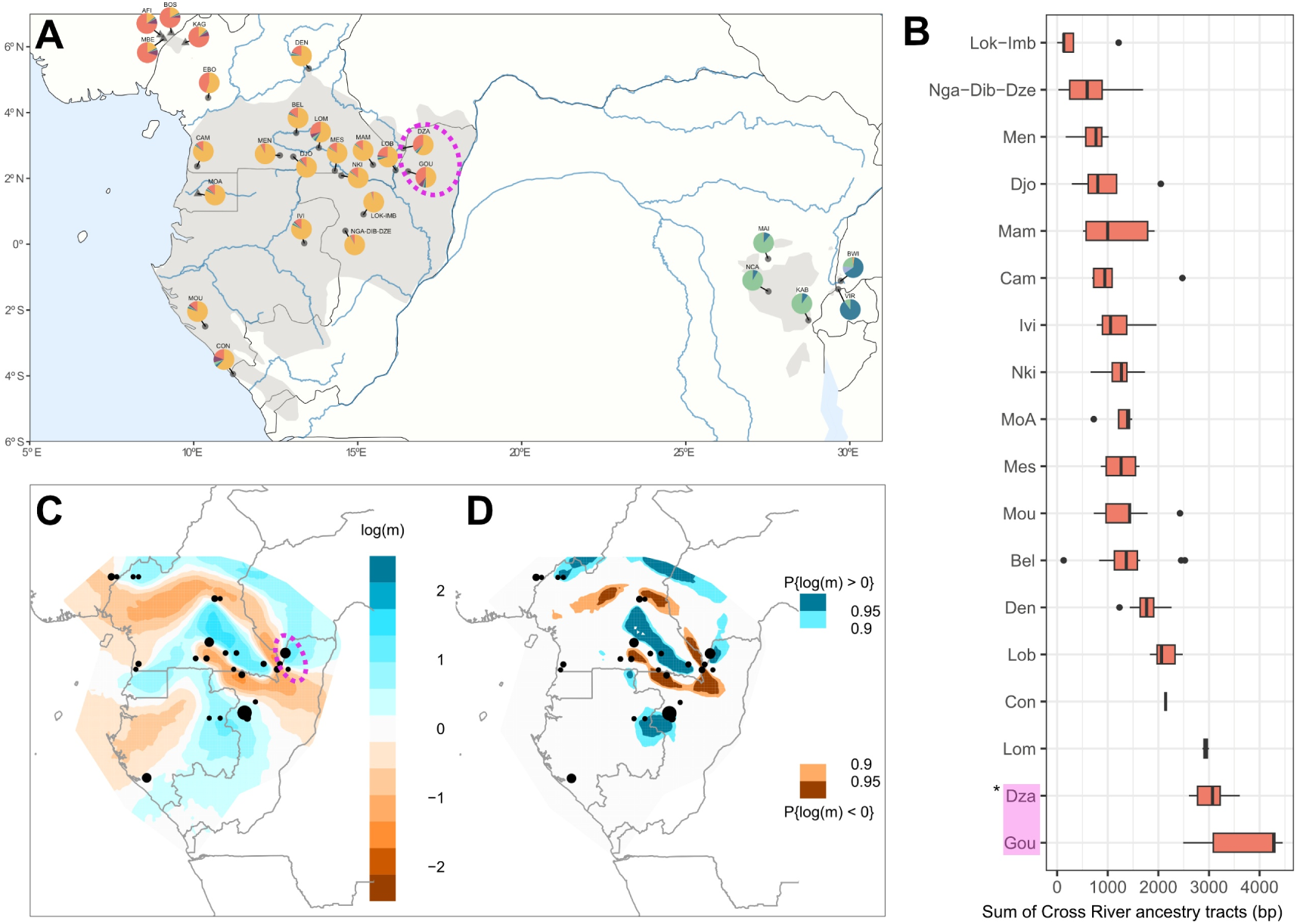
Connectivity and ancestry in gorilla populations. Pink color highlights Dzanga-Ndoki (Dza) and Goualougo (Gou) sites in all plots. A) Mean proportion of ancestry tracts as estimated with admixfrog, per sampling site for the four gorilla subspecies. Each pie-chart represents the mean ancestry of all samples from that site. Ancestry colours: Cross River gorillas (red), eastern lowland gorillas (green), western lowland gorillas (orange) and mountain gorillas (blue). The gorilla distribution range area is colored in grey for the four subspecies. B) The sum of Cross River gorilla ancestry tracts in western lowland gorillas grouped by sampling site. The X axis represents the length sum of all Cross River gorilla ancestry tracts. C) & D) EEMS Estimated Effective Migration Surfaces (EEMS) in the western gorilla species. C) Plotted are the posterior mean migration rates, where the colour indicates deviation from isolation by distance between the sampling sites, blue indicates high migration rate and orange indicates low migration rate. D) Posterior probabilities of the estimated effective migration surfaces; plotted are the values above 0.9 and 0.95 (blue for high migration rate and orange for low migration rate values).

We further assessed patterns of recent connectivity between and within the gorilla subspecies. Using EEMS (Petkova et al. 2016), we find reduced effective migration between the gorilla species, as well as to a lesser extent between the subspecies (Supplementary Figure 85). Within the western gorilla species, EEMS showed reduced migration between the subspecies, separated by the Sanaga River, and between some western lowland populations. Gorillas at Dza and Gou sampling sites, in the north-eastern distribution of western lowland gorillas (pink cluster in Figure 3), appeared genetically isolated from the rest of western lowland populations - matching with their geographical separation by the Sangha River, and its affluents, the Kadei and Ngoko rivers (Figure 3C, 3D). In contrast, these north-eastern western lowland gorillas exhibit higher migration with Cross River gorillas, suggesting a genetic corridor between this cluster of western lowland gorillas and Cross River gorillas (Figure 3C, 3D). In addition, another clear genetic barrier was detected between the sampling sites from Cameroon located above the Ngoko river (also known as the Dja river) (Lob, Mam, Bel, Den, Nki, Mes, Lom) and the sites below the Ngoko river (Djo and Men) and those from Odzala-Kokoua National Park in the Republic of Congo (Lok-Imb, Nga-Dib-Dze and Ivi). An increased effective migration rate between these Cameroon sampling sites was also detected (Lob, Mam, Bel, Den, Nki, Mes, Lom). Another reduction in migration rates between north and south Gabon was detected, which corresponds to the Ogooué river, but this was not statistically significant, probably due to the lower density of sampling sites in this region (Figure 3C, Figure 3D).

### Geolocalization of gorillas

Previous fine-scaled non-invasive sampling of wild chimpanzee populations enabled the geolocalisation of chimpanzees (Fontsere et al. 2022). Here, we predicted the geographic location of gorilla samples using chromosome 21 data (see Supplementary section 5)

First, we applied this methodology to predict geographic coordinates for the 224 georeferenced non-invasively collected samples [192 faecal samples (including 3 faecal samples from Fontsere et al. 2022), 32 hair samples (including hair samples from Alvarez-Estape et al. 2023), consisting of 27 eastern lowland gorillas, 41 mountain gorillas, 18 Cross River gorillas and 138 western lowland gorillas]. We removed the coordinates of the tested sample in a leave-one-out process. This approach was applied to all the samples in the dataset one by one. We calculated the difference between the predicted and geographic location for each sample. Specifically, we used the point distances between the tested samples and their neighbour points within the first two principal components (PC1, PC2) of each subspecies-specific PCA (Figure 4). The four nearest neighbour points of the tested sample were used to predict its geographic coordinates, applying a mean weighted by the Euclidean distances of each point to the tested sample point and the amount of difference explained in PC1 and PC2.

**Figure 4.**
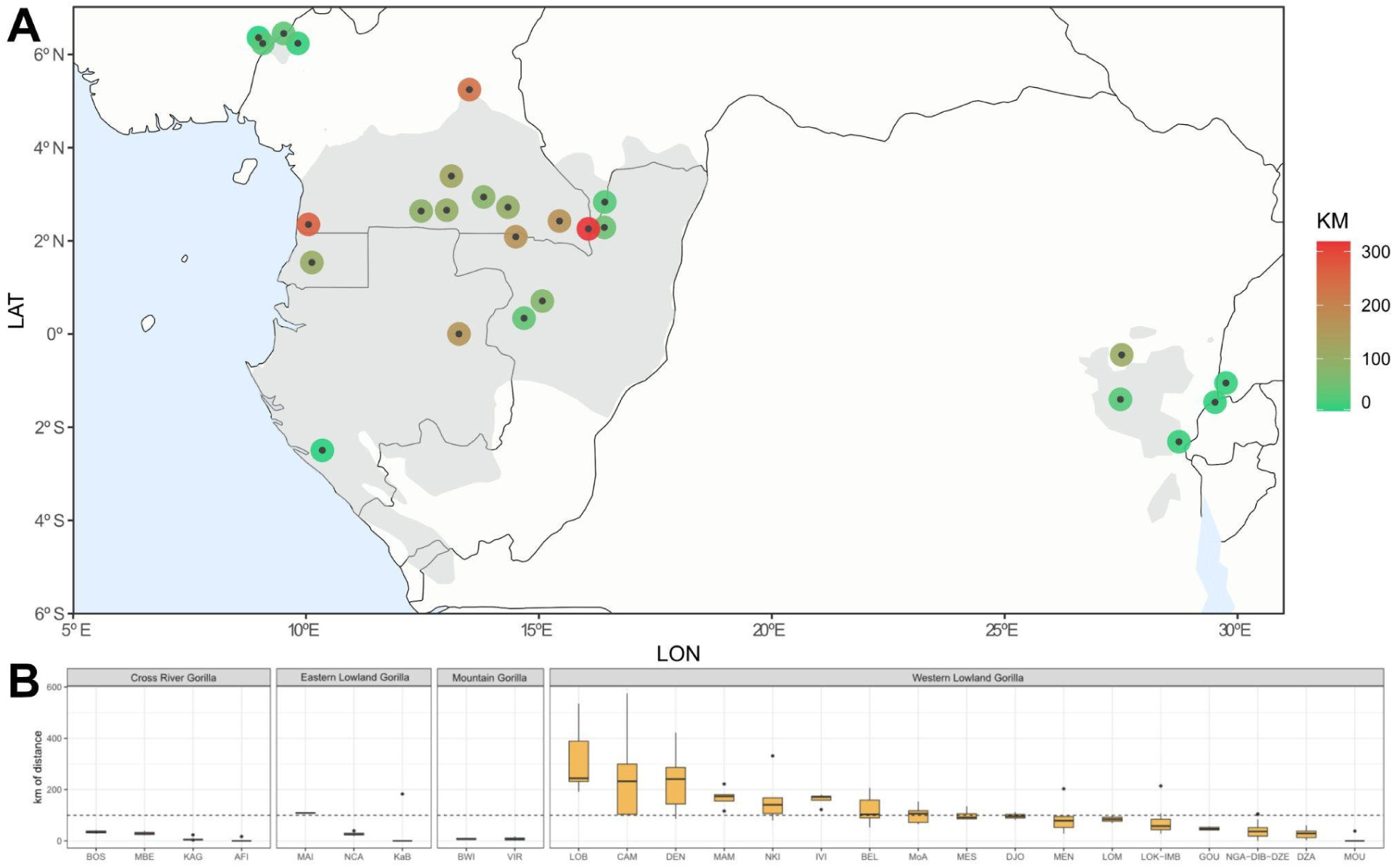
Geolocalization. A. Geographic distribution of tested sites for the geolocalization analysis. Sampling sites are coloured by the mean difference between true and predicted coordinates for all samples from the considered site (in kilometres). B. Box-plots of the difference between true and predicted coordinates for samples per sampling site, separated by subspecies. The y-axis represents the kilometers of difference between the real coordinates of each site and the predicted locations.

For genetically differentiated populations such as the Bwindi and Virunga mountain gorilla populations, the predicted locations were accurate and precise (Figure 4B). This was also the case for the western lowland gorilla population cluster of Dza and Gou (Dza - 28.15 km, and Gou - 47.98 km), and Mou (4.35 km) (Figure 4B). Low prediction accuracy was seen for sites for which the FST values were low between them, but the geographic distance was high to the neighbouring sites. This was the case for the western lowland sites Lob, Mam, Den, Nki, Bel and Mes. The mean kilometres of error when predicting geographic coordinates of all tested samples for all sampling sites considered in this analysis was 65 km.

Second, we aimed to geolocalise previously published wild-born gorilla samples (Prado-Martinez et al. 2013a; Prado-Martinez et al. 2013b; Xue et al. 2015; Alvarez-Estape et al. 2023; Pawar et al. 2023; Kuderna et al. 2023) and the newly sequenced FTA card and museum samples, using the non-invasive georeferenced samples as a reference dataset. We used the same approach as explained above by adding these samples into the non-invasive subspecies-specific PCAs. The mountain gorillas tested were correctly assigned to their corresponding subpopulation, Bwindi or Virunga (Supplementary Figure 91B). We predicted the locations of eastern lowland gorillas whose location was unknown (Amahoro, Itebero, Pinga, Serufuli) (Supplementary Figure 91A). The geographic coordinates for Itebero were predicted near the Nkuba Conservation Area. The geographic coordinates of Amahoro, Pinga and Serufuli were predicted near the Walikale area, with Serufuli being the closest one. We used zoo records (Linn, 2020) to corroborate the geographic predictions we made for the western lowland gorillas (Supplementary Figure 92). Zoo records (Linn 2020) for Coco and Floquet (also named ‘Snowflake’) assign their origins to Equatorial Guinea, as does PCA analysis; we indeed predicted the geographic coordinates for Coco to Equatorial Guinea. For Floquet, we predicted its origin to an area of Cameroon close to Equatorial Guinea. We predicted Katie (B650) and Katie (KB4986) originate in southern Gabon. Likewise, we predicted that Tzambo, Porta, Vila, Iris, Carolyn and Dolly originate from Odzala-Kokoua National Park or its surroundings. None of the tested western lowland gorillas was predicted to be from the area of Dza and Gou. Geographic coordinates for the Cross River gorilla Nyango were predicted between Boshi and Mbe areas, being closer to Boshi (Supplementary Figure 94); in agreement with PCA of Cross River gorillas (Figure 39 Supplementary) and previous studies (Alvarez-Estape et al. 2023).

Finally, we predicted the locations of the FTA card and museum samples following the same approach (Supplementary Figures 95-96). For the FTA cards, the predicted geographic coordinates were compared to the reported origin from where each individual was found (reported in Table 3 Supplementary). For some museum samples, information regarding the country of origin was available and was used to corroborate the predicted geographic coordinates. Of the 12 museum samples tested, we only found discrepancies with the museum metadata provided for one individual, for which the origin reported was the Republic of Congo, and the predicted geographic coordinates were in Cameroon in the northern part of the western lowland distribution (see Supplementary section 5).

## Discussion

Analysis of non-invasively collected samples of wild populations for population genomics is an active research area (Ostridge et al. 2025; Alvarez-Estape et al. 2023; Fontsere et al. 2022; Orkin et al. 2021). In this study, we performed extensive non-invasive sampling of three of the four subspecies of gorilla. This is the first time the target-capture hybridisation protocol established in chimpanzees (Ostridge et al. 2025; Fontsere et al. 2022) has been applied to other species, and we illustrate the feasibility of this approach. Although the estimated divergence between gorillas and humans and chimpanzees is ∼13 mya (Besenbacher et al. 2019), the capture of gorilla chromosome 21 using chimpanzee baits was successful. This is also the first time the capture protocol has been applied beyond faeces to hair samples.

The availability of exome data for chromosome X and chromosome 4 allowed us to investigate differences in effective population sizes of these chromosomes across the gorilla subspecies. As expected, we observed reduced diversity (a proxy for effective population size) on chromosome X compared to chromosome 4. In field studies, western lowland gorillas typically consist of single male breeding groups (with one silverback), whereas in mountain gorillas ∼40% of groups are multimale (Robbins 1999; Robbins et al. 2004; Bradley et al. 2005). We aimed to test whether different mating systems could be observed at the genetic level. While we observed some variation in the regression slopes of heterozygosity of chromosome 4 and chromosome X between the subspecies, we hypothesise that our analysis may be underpowered, and further sampling may disentangle the signal.

With our sampling scheme, we increased genetic data availability, particularly for western lowland gorillas, allowing us to characterise their population structure at fine-scale. We identify three population clusters within western lowland gorillas. In Fünfstück et al. (2014), using fewer sampling sites, two of the three population clusters were also identified (Dzanga-Ndoki National Park with Nouabalé-Ndoki National Park (Gou) and Odzala-Kokoua National Park); this dataset did not include samples from the third population cluster we identified here. The additional cluster we identified (population cluster 1 of PCA, Figure 1C) was formed by sites from southern Gabon, where previous studies showed a shared mitochondrial haplogroup with samples from the Odzala-Kokoua area and low nuclear microsatellite distances with sampling sites in Dzanga-Ndoki National Park (Anthony et al. 2007). Gorillas in the north-eastern population cluster of Dzanga-Ndoki and Gou (Nouabalé-Ndoki), two sampling sites located inside the TriNational Sangha Complex, are genetically differentiated from the rest of the western lowland sampling sites. Population connectivity analysis suggests that this is due to the Sangha River and its affluents, the Kadei and Ngoko rivers, forming a geographic barrier that separates the Dza and Gou gorillas from the rest of the western lowland gorillas. This corroborates observations of the Sangha River in particular, partitioning genetic diversity of western lowland gorillas, in analysis of mitochondrial haplogroups and microsatellites (Anthony et al. 2007; Fünfstück et al. 2014).

This north-eastern cluster of western lowland gorillas (Dza and Gou) is particularly interesting, as they also show the highest levels of shared ancestry with Cross River gorillas. This is in contrast to the rest of the western lowland gorilla populations, for which allele sharing with Cross River gorillas was much lower or not detected (Supplementary Figure 73). This signal of excess allele sharing between specific populations of the two western gorilla subspecies is similar to that identified by Fontsere et al. (2022) between central chimpanzees of Goualougo (sympatric to western lowland gorillas) and Nigeria-Cameroon chimpanzees of Gashaka (distributed near Cross River gorillas). One possible explanation is the existence of a past corridor in this region, which enabled connectivity of different chimpanzee subspecies and different gorilla subspecies. Alternatively, this signal could have been generated by ancestral population structure in western gorillas, which was proposed by the demographic inference of Thalmann et al. (2007). Future work may explore which scenario was more likely in western gorillas.

The recently documented gorilla population in the Ebo forest (Morgan et al. 2003), a key biodiversity area, is geographically intermediate between, and geographically isolated from present-day Cross River gorillas and western lowland gorillas. Morgan et al. (2003) have previously hypothesised that the gorillas at Ebo forest may be a remnant population of a previous wider range of gorillas. Here, we obtained genetic data from one gorilla from the Ebo forest, as it was not possible to obtain enough endogenous DNA from the other samples we collected from this area. This Ebo forest sample appeared mixed or intermediate between Cross River gorillas and western lowland gorillas. We caution that this single gorilla has low coverage (1.42-fold) and relatively high levels of human contamination (1.5%). However, we provide a suggestion of the genetics of the gorillas in the Ebo Forest, beyond microsatellite data (Anthony et al. 2007). Further samples sequenced to good coverage are needed to better characterise the population structure, genetic diversity and taxonomic status of the geographically unique gorilla population at Ebo forest.

Population genomics can be useful in a conservation genomics context, as it can provide the data to geolocate trafficked animals (Oklander and Soto-Calderón 2024). We provide a step towards this for gorillas by applying a geolocalization method using the distance of PCA points to predict a sample’s genetic origin. We first assessed the performance of the method on our dataset of georeferenced non-invasively collected gorilla samples, resulting in 65 kilometers of mean error. Population structure and genetic differentiation affected the ability of the method to predict the geographic locations of the gorilla samples. For instance, location prediction for the western lowland gorillas from Den had poor precision, due to their low genetic differentiation from other sampling sites. Five sampling sites are ∼240-430 km from Den, and the predicted locations of samples from Den are on average 233.56 kilometers from their true coordinates. In contrast, higher levels of population structure can aid the prediction. As mentioned, Dza and Gou are relatively genetically isolated from the other western lowland sampling sites, and so it is easier for the method to predict samples from these locations.

We predicted the geographic origin of 46 wild-born, previously published gorilla genomes (Prado-Martinez et al. 2013a; Prado-Martinez et al. 2013b; Xue et al. 2015; Pawar et al. 2023; Kuderna et al. 2023) using chromosome 21. All mountain gorillas tested were assigned correctly to their respective subpopulation (Virunga or Bwindi). Eastern lowland tested individuals were predicted to be from the Nkuba Conservation Area (NCA) and Walikale. PCA showed a clear separation between the Kahuzi-Biega (KaB) sampling site from the rest (Supplementary Figure 41), as previously reported (Michel et al. 2023). More samples from further sampling sites will be needed to improve population representation of eastern lowland gorillas for future geolocalization analysis. For western lowland gorillas, sites were not equally covered (i.e. few samples at some sampling sites and missing sampling sites across the distribution), likely affecting the ability to predict locations, *e.g.* from Equatorial Guinea or southern Gabon and eastern Cameroon, which are underrepresented in our dataset. Other sampling sites such as Dza and Gou had enough samples for the analysis, but none of the wild-born gorilla samples were predicted to these localities, acting as a true negative case.

We note that previous work by Das and Upadhyai (2019) based on a different algorithm predicted the locations of gorillas from Prado-Martinez et al. (2013a) to their country of origin. Here, using our newly generated georeferenced chromosome 21 data, we expand the reference dataset, allowing us to predict location to a finer-scale resolution than per country, for areas well covered by our sampling scheme. For example, the samples Katie-B650 and Katie-KB4986 exemplify this improvement, where we predicted their location to southern Gabon, in accordance with population structure results. Whereas, Das and Upadhyai (2019), assigned Katie-B650 and Katie-KB4986 to the extremity of the Congolese-Cameroon cline due to missing representatives from the southern Gabon. In addition, we were able to infer the geographic location of rescued gorillas sampled using FTA cards technology and gorilla museum samples from European natural history collections (Supplementary Figure 95-96). These results are a step for future work in this field, which would be useful for detecting hotspots of illegal trafficking and for localising museum specimens lacking complete metadata.

## Limitations

Target-capture hybridisation of gorilla chromosome 21 was performed using RNA baits designed on the chimpanzee assembly and previously used in Fontsere et al. (2022). Divergence may affect the specificity of the capture; specifically, capture specificity decreases as divergence from the baits increases (Jin et al. 2012). We observe that the gorilla chromosome 21 data has relatively high levels of missingness even in samples with a mean coverage of 5X-10X.

We performed target-capture hybridisation of the exome for the same gorilla faecal samples as for chromosome 21, but a technical problem during bait generation led to only partial capture of the whole exome (for details, see Supplementary Information). Nonetheless, the exome of chromosome X and several autosomal chromosomes were successfully captured, and we therefore make use of the high number of samples to analyse chromosome X: autosome comparisons across the gorilla subspecies.

Non-invasively collected samples, such as faeces and hair, have lower quality, lower endogenous DNA content and higher amounts of contamination than invasive samples, such as blood. Our filtering criteria and quality controls aim to limit the impact of potential contamination, which to some extent limits the sample size for some analyses.

Here, we increased the availability of genetic georeferenced data for wild gorilla populations. However, sampling gaps remain, and some sampling sites are represented by few individuals, which future work should address to gain a more complete genetic representation across the range of wild gorillas. This would likely improve our ability to geolocate gorillas from currently underrepresented regions.

## Conclusion

Non-invasively collected samples from wild gorillas allowed us to genetically characterise western lowland gorillas at the population level. We detected three distinct populations of western lowland gorillas and found that the Sangha River and its affluents, the Kadei and Ngoko rivers acted as geographic barriers to gene flow. This will be useful for the development of conservation strategies for these critically endangered populations. Specifically, we recommend that connectivity should be maintained within the three clusters of western lowland gorillas identified. We also detected past gene flow between Cross River gorillas and north-eastern western lowland gorillas, which may have been facilitated by a corridor also used by chimpanzees. Finally, we predicted geographic locations for wild-born gorillas as a proof of concept for future applications to geolocalise trafficked or rescued gorillas.

## METHODS

### Sample collection

For this study, 265 faecal samples and 15 hair samples were collected in a non-invasive manner from 26 sampling sites across the geographic ranges of three of the four extant gorilla subspecies (mountain gorillas, western lowland gorillas and eastern lowland gorillas). Additionally, 12 western lowland gorillas were sampled using FTA card samples from 10 sites, and 11 gorilla museum specimens. This was possible due to extensive collaboration with researchers in the field over several years. A full list of field sites is listed in Table 1 Supplementary. Research approval and sample permits were obtained from all countries involved, including country-specific permits. Faecal samples are exempt from Convention on the Trade in Endangered Species of Wild Fauna and Flora (CITES) permits; however, CITES permits were obtained for the hair and FTA card samples.

### Sample preparation and DNA extraction of faecal and hair samples

Faecal samples were prepared in a clean-hood that had been cleaned with bleach-water-ethanol. All faecal sample storage before DNA extraction is described in Table 3 in the Supplementary. We used 500mg (for fresh) and 100mg (for dried) faecal samples for DNA extraction. DNA was extracted from the faecal samples following 2 different approaches, depending on the sample provider (Table 3 Supplementary). The first one was using the QIAamp Fast DNA Stool Mini Kit (Qiagen) protocol, and the second one was following the same procedure as Michel et al. 2022. Hair samples were cleaned with 70% EtOH solution and cut into small pieces, and DNA was extracted from the hairs using the QIAamp DNA Investigator Kit (Qiagen). Faecal and hair samples were sonicated using a Q800R3 sonicator (Qsonica, USA) to obtain fragments of approximately 200 base pairs. Faecal samples were sonicated at 40% amplitude in pulse mode (15 seconds on / 15 seconds off) for a total active sonication time of 15 minutes. Hair samples, on the other hand, were sonicated at 20% amplitude without pulse mode for 15 minutes.

### Target Space Design

The chromosome 21 RNA baits used here had been designed in (Fontsere et al. 2022) using the PanTro4 assembly with 3x tiling density and fit into two Tier 5 custom arrays from SureSelect Agilent. The exome RNA baits used here were designed by SureSelect Agilent based on the panTro6 assembly with 3x tiling density, into three Tier 5 custom arrays.

### Library preparation, capture and sequencing of chromosome 21 and the exome for gorilla faecal and hair samples

Libraries were generated for the faecal and hair samples following the “BEST” protocol (Carøe et al. 2017), including minor modifications applied in (Fontsere et al. 2022). Shallow sequencing was performed (producing 4GB per library) to calculate the endogenous DNA content of each library. Unique in-line barcodes were added to each prepared library before capture, following Fontsere et al. (2022). We calculated the endogenous DNA of each newly sequenced sample by dividing the final number of filtered reads by the total of sequenced reads following https://github.com/claudefa/Mapping_Capture_Experiments/tree/main/hDNA. Libraries were pooled based on the endogenous DNA of each sample following Fontsere et al. (2022), Fontsere et al. (2021b), and Hernandez-Rodriguez et al. (2018) to achieve equi-endogenous pooling. Each pool was then divided into 4 aliquots to have two replicates for each capture procedure (chromosome 21 and exome).

The pooled libraries were captured using the capture protocol previously established in chimpanzees (Fontsere et al. 2021b; Fontsere et al. 2022). Captured libraries were sequenced on an Illumina NovaSeq 6000 with 2×150 bp setup at the National Center of Genomic Analysis (CNAG, Barcelona, Spain).

### DNA extraction, library preparation and sequencing of FTA card and museum gorilla samples

3 mm diameter punches were cut from the FTA cards, and DNA was extracted using the QIAamp DNA Investigator Kit (Qiagen). Libraries were made following the “BEST” protocol (Carøe et al. 2017), including minor modifications applied in Fontsere et al. (2022).

Tooth powder was extracted from 11 museum samples from Vienna (NMW 7136 & NMW 3111/ST 665), Berlin (ZMB_Mam_83519 & ZMB_Mam_37523), Frankfurt (59158, 96550, 1132, 17826 & 16180), Salzburg (HNS-Mam-S-0525) and Bonn (ZFMK_MAM2015-0479) museum collections. 50 mg of the tooth powder per sample was used for DNA extraction following the protocol described in Dabney et al. 2013, in an ancient DNA laboratory at UPF, following all required cleaning steps (bleach, ethanol and post-usage UV lights) and wearing specific clothes to avoid contamination. Libraries were made following the single-chain reaction (SCR) protocol described in Kapp et al. (2021).

We performed whole genome sequencing of the FTA card and museum samples, using an Illumina NovaSeq 6000 with 2 x 150 bp setup at the National Center of Genomic Analysis (CNAG, Barcelona, Spain).

### Data Processing faecal and hair chr21 and exome captures

We demultiplexed sequencing reads of each hybridisation pool originating from faecal and hair samples using Sabre (https://github.com/najoshi/sabre) using the unique in-line barcodes. Reads were trimmed (of Illumina adapters), filtered and merged using FASTP (Chen et al. 2018) (v0.23.2). We mapped merged and paired (not merged) reads to the human reference genome hg38 (to avoid reference bias between gorilla species, which may occur if the gorilla reference genome was used) using BWA mem (Li et al. 2009a) (0.7.12-r1039). Read groups were added using AddOrREplaceReadGroups with Picard (https://broadinstitute.github.io/picard/) (1.95). We merged all bam files of each unique sample using samtools (Li et al. 2009b) (1.15), where the same library had been captured more than once and where the same captured library had been sequenced in different lanes. Duplicates and reads shorter than 35 bp were removed from the merged bam files using the MarkDuplicates option of Picard (https://broadinstitute.github.io/picard/) (1.95). Secondary alignments and reads with mapping quality below 30 were also removed using samtools (Li et al. 2009b) (1.15). We intersected the resulting reads to the chromosome 21 or exome target space respectively, using bedtools (Quinlan and Hall 2010) (2.29.0).

### Data Processing Museum and FTA card whole genomes

Whole genome paired reads generated from museum samples were collapsed using FASTP software using the “--merge” option applying an overlap length of 11 (--overlap_len_require 11) allowing 2 differences (--overlap_diff_limit 2), trimming polyg (--trim_poly_g) with a minimum length of 10 (--poly_g_min_len 10), and filtering by length and quality (--length_required 30 & --cut_mean_quality 20). The collapsed reads were mapped to the human reference genome (hg38) using BWA aln (aln -l 16500 -t 8 -n 0.01 -o 2) and samse commands.

Raw reads sequenced from FTA card samples were trimmed from Illumina adapters using FASTP software (Chen et al. 2018) (v0.23.2). Merged and unmerged reads were mapped to the human reference genome (hg38) using BWA mem (Li et al. 2009a) (0.7.12-r1039).

For both museum and FTA card samples, we included read group information using the AddOrReplaceReadGroups option of Picard (https://broadinstitute.github.io/picard/) (1.95). All bam files from unique samples were merged using samtools (Li et al. 2009b) (1.15). Duplicates were removed using MarkDuplicates option of Picard (https://broadinstitute.github.io/picard/) (1.95). Secondary alignments and reads with mapping quality below 25 or 30 (Museum and FTA card, respectively) were removed using samtools (Li et al. 2009b) (1.15).

### Quality control

We calculated the amount of sequenced, mapped and filtered reads for all newly sequenced samples (chromosome 21, exome, FTA card and museum samples). We evaluated the success of the capture protocol in gorillas by calculating the library complexity, enrichment factor, sensitivity and specificity, as described in Fontsere et al (2021b) (See Supplementary Section 1).

We calculated the depth of coverage for all newly sequenced samples using mosdepth (Pedersen and Quinlan 2018) (0.3.3-GCC-11.2.0). We assessed potential human contamination using HuConTest (Kuhlwilm et al. 2021) which evaluates the percentage of human-like alleles (as opposed to gorilla-like alleles) for each sample. We discarded samples with less than 0.5X of mean coverage and more than 1% of human contamination.

### Previously published gorilla data

We downloaded 84 previously published high-coverage whole genome data for gorillas from the 4 subspecies (Prado-Martinez et al. 2013a; Prado-Martinez et al. 2013b, Xue et al. 2015; Pawar et al. 2023; Alvarez-Estape et al. 2023; Kuderna et al. 2023) from ENA and mapped to the human reference genome hg38. The chromosome 21 on-target space from these samples was extracted using intersectBed from bedtools (Quinlan and Hall 2010) for subsequent analysis. The exome on-target space was extracted from samples of Alvarez-Estape et al. 2023 following the same approach.

### Genotype Likelihoods

Genotype likelihoods were calculated using ANGSD (0.935) (Korneliussen et al. 2014) for the whole dataset (of the newly sequenced and published genomes), both for chr21 and exome data, with the GATK model (-GL 2). We used these flags: -uniqueOnly 1 -remove_bads 1 - only_proper_pairs 1 -trim 0 -C 50 -GL2 -minMapQ 25 -minQ 20 -doGlf 2 -SNP_pval 1e-6.

### Relatedness

Kinship relations were assessed per gorilla sampling site using NgsRelate (Korneliussen et al. 2015) for each pair of individuals that passed our filtering criteria (coverage > 0.5X and human contamination < 1%). We filtered out identical and first-degree relatives. One individual of each first-degree related pair detected was removed, keeping the sample with higher coverage or fewer further kinship relations

### Population Structure

Principal Component Analysis (PCA) was constructed using PCAngsd (Meisner and Albrechtsen 2018) (v1.21). We performed PCA at different levels; for all samples, per species, per subspecies and within western lowland gorilla groups as population structure was detected. We applied MAF filtering to remove singletons.

Admixture was assessed using NGSadmix (Skotte et al. 2013). We also applied MAF filtering to remove singletons. We performed 25 different runs for each of the 10 Ks. The best K was inferred using the best likelihoods of each run and K.

We calculated genetic distances between subspecies and between populations using ANGSD (Korneliussen et al. 2014) (0.935). At the subspecies level, each subspecies was downsampled to 18 individuals (to match the number of available ELG in the dataset). At the population level, we randomly downsampled each sampling site to 8 samples. We considered only those sampling sites with >=8 gorillas, and so we merged sites closer than 25 km (see Supplementary Table 3).

Site allele frequency likelihood was calculated for each site using ANGSD (Korneliussen et al. 2014) with flags: -dosaf 1 -gl 1 -uniqueOnly 1 -remove_bads 1 -only_proper_pairs -trim 0 -C 50 -baq 1 -minMapQ 20 -minQ 20. The 2D-site frequency spectrum was calculated using the realSFS, and the command “realSFS fst index” was used to obtain fst binary files. The command “realSFS fst stats” was used to obtain FST values for each comparison.

We calculated heterozygosity per sample for all samples (for chromosome 21 and the exomes of chromosome X and chromosome 4) using realSFS within ANGSD (Korneliussen et al. 2014) (0.935) and flags: -dosaf 1 -gl 1 -uniqueOnly 1 -remove_bads 1 -only_proper_pairs 1 - trim 0 -minMapQ 25 -minQ 20 -setMaxDepth 200 -doCounts 1 -C 50 -baq 1. For the comparison of the heterozygosity of chromosome X to chromosome 4, we retained only females in the analysis. We downsampled samples >1.5X to 1.5X to avoid bias in coverage (for chromosome 21).

### Gene flow

Ancestry fragments across chromosome 21 were inferred using admixfrog (Peter 2020) (0.7.2). Admixfrog uses a Hidden Markov Model to infer ancestry fragments from low-coverage and contaminated data. We considered the sources of admixture: Cross River gorillas, western lowland gorillas, eastern lowland gorillas and mountain gorillas. The reference file was created using *admixfrog-ref* with –states (CRG WLG ELG MNG ANC HUM). We used the human reference genome as the ancestral state and for contamination. Sample input files were prepared using *admixfrog-bam* on the bam files. Admixfrog was run to infer the ancestry fragments across chromosome 21 for all samples using the *admixfrog* command.

We assessed the possibility of gene flow using D-statistics in ANGSD (Abbababa) (Korneliussen et al. 2014) (0.935). We used the “Abababa2-multipop” command to include multiple individuals for each studied population. As above, for the FST analysis, we randomly downsampled each sampling site to 8 samples and merged sites closer than 25 km. The flags used to calculate the D-statistics were: -doMajorMinor 1 -doMaf 1 -minMaf 0.01 -useLast 1 - rmTrans 0 -blockSize 5000000 -uniqueOnly 1 -remove_bads 1 -minMapQ 25 -minQ 20. A size file was used, indicating the number of individuals of each population.

### Variant Calling

For samples with more than 5X coverage, we used GATK haplotypecaller with -ERC GVF option to call all sites for all samples (McKenna et al. 2010) (4.2.5.0). A gVCFs files were combined using GATK CombineGVCFs. The combined gVCF file was genotyped for all non-variant sites using GATK GenotypeGVCFs.

The resulting VCF file was filtered to keep just the variants. In addition, a callable region was calculated using the public high-quality and high-coverage genomes, filtering variants due to genotype quality (min 30) and coverage (6-40X). All sites outside this callable region were removed from the final set of variants in the VCF file.

Filters were applied accordingly to each analysis. In order to define proper filters for this dataset, statistics of the final variants were performed. Genotype quality was measured for all variants in order to evaluate the overall quality of the variants. Faecal sample genotype quality was expected to be lower when compared to high-quality samples such as blood or tissue. Depth of coverage was extracted for all positions using the --site-mean-depth option of VCFtools (Danecek et al. 2011). Missing data for each sample and each site was calculated from the VCF file using the --missing-indv and --missing-site options of VCFtools (Danecek et al. 2011).

### Called variants: population structure

Detection of heterozygous variants is influenced by sample coverage. To assess this in the gorilla faecal samples, we downsampled samples to either 6X, 10X, and 20X. We applied the following filters: 30 as minimum variant quality and a minimum genotype quality of 10. In addition, we filtered out all positions outside the callable region. We checked the individual missingness after all the filtering in order to remove samples containing high missing values (>40% missing data). Finally, we calculated heterozygosity in the called genotypes at 6X, 10X and 20X.

The number of heterozygous positions inside the callable region for each sample was counted and divided by the callable region length. This was the measure we used as heterozygosity to study genetic diversity. We used bcftools view -s (Danecek et al. 2021) to extract each sample from the VCF file and -R option to extract the positions inside the callable region. We used bcftools view -i ‘GT=”het”’ to extract heterozygote positions and then count all of them.

### Called variants: population connectivity

Connectivity and genetic barriers among gorilla populations were assessed using Estimated Effective Migration Surfaces (EEMS) (Petkova et al. 2016). EEMS infers the migration rates across an established landscape by comparing observed genetic dissimilarities to those expected under a spatial model.

Here we used samples with a coverage >=5X, low levels of missingness (<40%) and for which their geographic coordinate information was available. We also filtered out variants with more than 20% of missing genotypes, genotype quality below 20 and depth of coverage below 3, as well as only including the on-target space and biallelic sites.

The required coordinate input file was created with the sample IDs and their corresponding geographic coordinates. We also generated a boundary file defining the studied area, expanding the distribution ranges given in IUCN shapefiles for the gorilla species (UNEP-WCMC & IUCN 2016, IGCP & WCS 2019) to include possible past corridors. A params file was also generated specifying: the number of samples, positions used, output directory, and input file prefixes required for the analysis.

The VCF file was converted to PLINK format to create a dissimilarity matrix using the provided bed2diff script (https://github.com/dipetkov/eems). We ran the EEMS script using the params file (generated above), containing the path to all input files needed.

### Geolocalisation

We used the subspecies-specific PCAs, constructed using genotype likelihoods, to geolocate samples of each subspecies. For this analysis, we used all georeferenced samples which passed filtering thresholds (less than 0.5X of mean coverage and more than 1% of human contamination & non-related individuals). We used the point distances in the first two principal components of each subspecies PCA, using the “get.knnx” function from the FNN package (Beygelzimer et al. 2024) in R. The four nearest neighbour points of the tested sample were used to predict the geographic coordinates of the test sample, in a leave-one-out process. We applied a mean weighted by the Euclidean distances of each point to the tested sample and the amount of difference explained in the first two components used.

Applying a leave-one-out process, geographic coordinates for the 224 georeferenced non-invasively collected samples (192 faecal samples (including 3 faecal samples from Fontsere et al. 2022), 32 hair samples (including hair samples from Alvarez-Estape et al. 2023), consisting of 27 eastern lowland gorillas, 41 mountain gorillas, 18 Cross River gorillas and 138 western lowland gorillas) were predicted. The difference between the real and the predicted geographic location was measured for each sample and visualised using distribution plots to evaluate the overall prediction of all samples for each sampling site. FTA cards, museum samples and the publicly available genomes (Prado-Martinez et al. 2013a; Prado-Martinez et al. 2013b; Xue et al. 2015; Pawar et al. 2023; Kuderna et al. 2023) were also used to predict their respective geographic coordinates using the 224 georeferenced non-invasively collected samples. Geolocalization plots for chromosome 21 of the wild-born publicly available gorillas were generated.

## DECLARATIONS

All coauthors have approved this manuscript for publication

Conflict of interest statement. None declared.

### Data availability

The datasets generated and analysed during the current study are available in the ENA repository, [give ENA ID].

## Acknowledgements

We thank Yasmin Moebius, Mizuki Murai, Manasseh Eno-Nku, Martijn Ter Heegde, Amelia Meier, Hilde Vanleeuwe, Jean Claude Dengui, Paul Telfer, Abel Nzeheke, Luc Roscelin Tédonzong, Sonia Nicholl, Theophile Desarmeaux, Vianet Mihindou, Marcel Ketchan Eyong and Chad N. Evans for field sample collection, logistical and laboratory support. We thank Naoki Matsuura for logistical support for the fieldwork. We thank Agence Nationale des Parcs Nationaux, Gabon; Centre National de la Recherche Scientifique (CENAREST), Gabon; L’Institut Congolais pour la Conservation de la Nature (ICCN) and local landowners and community members, rangers and field assistants, gorilla trackers in DRC;; Ministère des Forêts et de la Faune, Cameroon; Ministère de l’Enseignement Supérieur, Central African Republic; Ministère de la Recherche Scientifique et de l’Innovation Technologique, Cameroon; Project Grands, Singes, Cameroon; Rwanda Development Board; Uganda National Council for Science and Technology (UNCST), Uganda; Uganda Wildlife Authority, Uganda; and WWF Central African Republic and Primate Habituation Programme staff for supporting the field research that allowed us to obtain the non-invasively collected samples. We thank the WWF Central African Republic and Primate Habituation Programme staff for the sample collection logistics. We thank the Ministère de la Recherche Scientifique et de l’Innovation and Ministère des Forêts et de la Faune for the permits, and the Antwerp Zoo Society, Belgium and Project Grands Singes, Cameroon for the sample collection logistics. We thank the Rwanda Development Board for the permits, and the Gorilla Doctors, Dian Fossey Gorilla Fund and Virunga National Park for the sample collection. We thank the Pan African Programme: The Cultured Chimpanzee (PanAf) for providing gorilla samples from: Uganda (Bwindi), Cameroon (Campo Ma’an, La belgique (Dja)), Nigeria (Mbe mountains), Republic of Congo (Conkouati, Goualougo) and Gabon (Ivindo, Loango, Lopé, and Mts de Cristal). We thank the Zoologisches Forschungsmuseum A. Koenig, Leibniz-Institut zur Analyse des Biodiversitätswandels in Bonn, in particular Eva Bärmann, Jan Decher, and Christian Montermann at the Section Theriology; thanks to Irina Ruf and Katrin Krohmann at the Mammalogy collection at Senckenberg Forschungsinstitut und Naturmuseum Frankfurt/M.; thanks to Frank Zachos and Alexander Bibl at Naturhistorisches Museum Wien; thanks to Robert Lindner at Haus der Natur – Museum für Natur und Technik in Salzburg; to Christiane Funk and Frieder Mayer at the mammalian collection at Museum für Naturkunde/Leibniz-Institut für Evolutions- und Biodiversitätsforschung in Berlin. Institutional support to CNAG was provided by the Spanish Government (Ministry of Science, Innovation and Universities) and the Generalitat de Catalunya through the Departament de Recerca i Universitats and the Departament de Salut. We thank Aida M. Andrés, Marc de Manuel and Ben Evans for helpful discussions.

## Author contributions

T.M.B. E.L. and M.K conceived the study. D.C., N.I., U.N., K.G., E.B., E.A.O., A.K.B., K.S., B.P., T.F., N.T., D.M, E.A.A., J.W., N.W.N., E.D., E.N., I.I., E.B., A.D., K.J.J., T.D., M.R., J.H., A.C.G., V.H., C.O., D.M., C.S., M.B., G.M., K.L., P.D., H.K., C.B., E.I., M.B., G.I.B., T.S.E., J.D.O, M.P., K.P., B.M., M.A., R.P., O.C., B.N., L.B., F.P., A.N., J.Y., M. I.-M., Y.T., S.F., T.B., U.M., B.N., F.P., B.L., and V.R. performed fieldwork, collected samples, provided or helped to have access to gorilla samples. I.G.G., M.G. provided access to sequencing data and laboratory resources. I.R.G. and H.P. performed the analysis. M.A.E., M.A., P.A.M. and S.R.P. performed experimental laboratory work. M.A.E., H.P. and I.R.G. performed data curation. T.M.B., E.L. and M.K. provided supervision. H.J.O., J.P.M., C.F., S.C.E. provided analytical support. I.R.G. and H.P. wrote the paper with input from all authors.

## References

Alvarez-Estape, Marina, Harvinder Pawar, Claudia Fontsere, Amber E. Trujillo, Jessica L. Gunson, Richard A. Bergl, Magdalena Bermejo, et al. ‘Past Connectivity but Recent Inbreeding in Cross River Gorillas Determined Using Whole Genomes from Single Hairs’. Genes 14, no. 3 (18 March 2023): 743. 10.3390/genes14030743.

Anthony, Nicola M., Mireille Johnson-Bawe, Kathryn Jeffery, et al. ‘The Role of Pleistocene Refugia and Rivers in Shaping Gorilla Genetic Diversity in Central Africa’. Proceedings of the National Academy of Sciences 104, no. 51 (2007): 20432–36. 10.1073/pnas.0704816105.

Battey, Cj, Peter L Ralph, and Andrew D Kern. ‘Predicting Geographic Location from Genetic Variation with Deep Neural Networks’. eLife 9 (8 June 2020): e54507. 10.7554/eLife.54507.

Bergl, R.A., Dunn, A., Fowler, A., Imong, I., Ndeloh, D., Nicholas, A. & Oates, J.F. 2016. Gorilla gorilla ssp. diehli. The IUCN Red List of Threatened Species 2016: e.T39998A102326240.10.2305/IUCN.UK.2016-2.RLTS.T39998A17989492.en

Bermejo, Magdalena, José Domingo Rodríguez-Teijeiro, Germán Illera, Alex Barroso, Carles Vilà, and Peter D. Walsh. ‘Ebola Outbreak Killed 5000 Gorillas’. Science 314, no. 5805 (8 December 2006): 1564–1564. 10.1126/science.1133105.

Besenbacher, Søren, Christina Hvilsom, Tomas Marques-Bonet, Thomas Mailund, and Mikkel Heide Schierup. ‘Direct Estimation of Mutations in Great Apes Reconciles Phylogenetic Dating’. Nature Ecology & Evolution 3, no. 2 (February 2019): 286–92. 10.1038/s41559-018-0778-x.

Beygelzimer, Alina, Sham Kakadet, John Langford, Sunil Arya, David Mount, and Shuai Li. 2024. FNN: Fast Nearest Neighbor Search Algorithms and Applications. R package version 1.1.4.1. https://CRAN.R-project.org/package=FNN.

Bradley, Brenda J., Martha M. Robbins, Elizabeth A. Williamson, H. Dieter Steklis, Netzin Gerald Steklis, Nadin Eckhardt, Christophe Boesch, and Linda Vigilant. ‘Mountain Gorilla Tug-of-War: Silverbacks Have Limited Control over Reproduction in Multimale Groups’. Proceedings of the National Academy of Sciences 102, no. 26 (28 June 2005): 9418–23. 10.1073/pnas.0502019102.

Caillaud, Damien, Florence Levréro, Romane Cristescu, Sylvain Gatti, Maeva Dewas, Mélanie Douadi, Annie Gautier-Hion, Michel Raymond, and Nelly Ménard. ‘Gorilla Susceptibility to Ebola Virus: The Cost of Sociality’. Current Biology: CB 16, no. 13 (11 July 2006): R489–491. 10.1016/j.cub.2006.06.017.

Carøe, Christian, Shyam Gopalakrishnan, Lasse Vinner, Sarah S. T. Mak, Mikkel Holger S. Sinding, José A. Samaniego, Nathan Wales, Thomas Sicheritz-Pontén, and M. Thomas P. Gilbert. ‘Single-tube Library Preparation for Degraded DNA’. Edited by Susan Johnston. Methods in Ecology and Evolution 9, no. 2 (February 2018): 410–19. 10.1111/2041-210X.12871.

Chen, Shifu, Yanqing Zhou, Yaru Chen, and Jia Gu. ‘Fastp: An Ultra-Fast All-in-One FASTQ Preprocessor’. Bioinformatics (Oxford, England) 34, no. 17 (1 September 2018): i884–90. 10.1093/bioinformatics/bty560.

Dabney, Jesse, Michael Knapp, Isabelle Glocke, Marie-Theres Gansauge, Antje Weihmann, Birgit Nickel, Cristina Valdiosera, et al. ‘Complete Mitochondrial Genome Sequence of a Middle Pleistocene Cave Bear Reconstructed from Ultrashort DNA Fragments’. Proceedings of the National Academy of Sciences 110, no. 39 (24 September 2013): 15758–63. 10.1073/pnas.1314445110.

Danecek, Petr, Adam Auton, Goncalo Abecasis, Cornelis A. Albers, Eric Banks, Mark A. DePristo, Robert E. Handsaker, et al. ‘The Variant Call Format and VCFtools’. Bioinformatics (Oxford, England) 27, no. 15 (1 August 2011): 2156–58. 10.1093/bioinformatics/btr330.

Danecek, Petr, James K. Bonfield, Jennifer Liddle, John Marshall, Valeriu Ohan, Martin O. Pollard, Andrew Whitwham, et al. ‘Twelve Years of SAMtools and BCFtools’. GigaScience 10, no. 2 (16 February 2021): giab008. 10.1093/gigascience/giab008.

Das, Ranajit, and Priyanka Upadhyai. ‘Application of the Geographic Population Structure (GPS) Algorithm for Biogeographical Analyses of Wild and Captive Gorillas’. BMC Bioinformatics 20, no. 1 (5 February 2019): 35. 10.1186/s12859-018-2568-5.

Estrada, Alejandro, Paul A. Garber, Anthony B. Rylands, Christian Roos, Eduardo Fernandez-Duque, Anthony Di Fiore, K. Anne-Isola Nekaris, et al. ‘Impending Extinction Crisis of the World’s Primates: Why Primates Matter’. Science Advances 3, no. 1 (6 January 2017): e1600946. 10.1126/sciadv.1600946.

Fontsere, Claudia, Peter Frandsen, Jessica Hernandez-Rodriguez, Jonas Niemann, Camilla Hjorth Scharff-Olsen, Dominique Vallet, Pascaline Le Gouar, et al. ‘The Genetic Impact of an Ebola Outbreak on a Wild Gorilla Population’. BMC Genomics 22, no. 1 (11 October 2021): 735. 10.1186/s12864-021-08025-y.

Fontsere, Claudia, Marina Alvarez-Estape, Jack Lester, Mimi Arandjelovic, Martin Kuhlwilm, Paula Dieguez, Anthony Agbor, et al. ‘Maximizing the Acquisition of Unique Reads in Noninvasive Capture Sequencing Experiments’. Molecular Ecology Resources 21, no. 3 (April 2021): 745–61. 10.1111/1755-0998.13300.

Fontsere, Claudia, Martin Kuhlwilm, Carlos Morcillo-Suarez, Marina Alvarez-Estape, Jack D. Lester, Paolo Gratton, Joshua M. Schmidt, et al. ‘Population Dynamics and Genetic Connectivity in Recent Chimpanzee History’. Cell Genomics 2, no. 6 (8 June 2022): None. 10.1016/j.xgen.2022.100133.

Fünfstück, Tillmann, Mimi Arandjelovic, David B. Morgan, Crickette Sanz, Thomas Breuer, Emma J. Stokes, Patricia Reed, et al. ‘The Genetic Population Structure of Wild Western Lowland Gorillas (*Gorilla Gorilla Gorilla*) Living in Continuous Rain Forest’. American Journal of Primatology 76, no. 9 (September 2014): 868–78. 10.1002/ajp.22274.

Fünfstück, Tillmann, and Linda Vigilant. ‘The Geographic Distribution of Genetic Diversity within Gorillas’. American Journal of Primatology 77, no. 9 (September 2015): 974–85. 10.1002/ajp.22427.

Grubb, Peter, Thomas M. Butynski, John F. Oates, Simon K. Bearder, Todd R. Disotell, Colin P. Groves, and Thomas T. Struhsaker. ‘Assessment of the Diversity of African Primates’. International Journal of Primatology 24, no. 6 (December 2003): 1301–57. 10.1023/B:IJOP.0000005994.86792.b9.

Hernandez-Rodriguez, Jessica, Mimi Arandjelovic, Jack Lester, Cesare de Filippo, Antje Weihmann, Matthias Meyer, Samuel Angedakin, et al. ‘The Impact of Endogenous Content, Replicates and Pooling on Genome Capture from Faecal Samples’. Molecular Ecology Resources 18, no. 2 (March 2018): 319–33. 10.1111/1755-0998.12728.

Hickey, J.R., Basabose, A., Gilardi, K.V., Greer, D., Nampindo, S., Robbins, M.M. & Stoinski, T.S. 2018. Gorilla beringei ssp. beringei. The IUCN Red List of Threatened Species 2018: e.T39999A17989719. 10.2305/IUCN.UK.2018-2.RLTS.T39999A17989719.en

IGCP and WCS (International Gorilla Conservation Programme and Wildlife Conservation Society). 2019. Gorilla beringei (spatial data). The IUCN Red List of Threatened Species. Version 2025-1. https://www.iucnredlist.org.

Imong, I., M. M. Robbins, R. Mundry, R. Bergl, and H. S. Kühl. ‘Distinguishing Ecological Constraints from Human Activity in Species Range Fragmentation: The Case of Cross River Gorillas: Species Range Fragmentation’. Animal Conservation 17, no. 4 (August 2014): 323–31. 10.1111/acv.12100.

Jin, Xin, Mingze He, Betsy Ferguson, Yuhuan Meng, Limei Ouyang, Jingjing Ren, Thomas Mailund, et al. ‘An Effort to Use Human-Based Exome Capture Methods to Analyze Chimpanzee and Macaque Exomes’. PLOS ONE 7, no. 7 (27 July 2012): e40637. 10.1371/journal.pone.0040637.

Kapp, Joshua D, Richard E Green, and Beth Shapiro. ‘A Fast and Efficient Single-Stranded Genomic Library Preparation Method Optimized for Ancient DNA’. Edited by Robert C Fleischer. Journal of Heredity 112, no. 3 (24 May 2021): 241–49. 10.1093/jhered/esab012.

Korneliussen, Thorfinn Sand, Anders Albrechtsen, and Rasmus Nielsen. ‘ANGSD: Analysis of Next Generation Sequencing Data’. BMC Bioinformatics 15, no. 1 (25 November 2014): 356. 10.1186/s12859-014-0356-4.

Korneliussen, Thorfinn Sand, and Ida Moltke. ‘NgsRelate: A Software Tool for Estimating Pairwise Relatedness from next-Generation Sequencing Data’. Bioinformatics 31, no. 24 (15 December 2015): 4009–11. 10.1093/bioinformatics/btv509.

Kuderna, L.F.K. et al. (2023) ‘A global catalog of whole-genome diversity from 233 primate species,’ Science, 380(6648), pp. 906–913. 10.1126/science.abn7829.

Kuhlwilm, Martin, Claudia Fontsere, Sojung Han, Marina Alvarez-Estape, and Tomas Marques-Bonet. ‘HuConTest: Testing Human Contamination in Great Ape Samples’. Edited by David Enard. Genome Biology and Evolution 13, no. 6 (8 June 2021): evab117. 10.1093/gbe/evab117.

Lester, Jack D., Linda Vigilant, Paolo Gratton, et al. ‘Recent Genetic Connectivity and Clinal Variation in Chimpanzees’. Communications Biology 4, no. 1 (2021): 1–11. 10.1038/s42003-021-01806-x.

Li, Heng, and Richard Durbin. ‘Fast and Accurate Short Read Alignment with Burrows-Wheeler Transform’. Bioinformatics (Oxford, England) 25, no. 14 (15 July 2009): 1754–60. 10.1093/bioinformatics/btp324.

Li, Heng, Bob Handsaker, Alec Wysoker, Tim Fennell, Jue Ruan, Nils Homer, Gabor Marth, Goncalo Abecasis, Richard Durbin, and 1000 Genome Project Data Processing Subgroup. ‘The Sequence Alignment/Map Format and SAMtools’. Bioinformatics (Oxford, England) 25, no. 16 (15 August 2009): 2078–79. 10.1093/bioinformatics/btp352.

Linn S.N., Bender U. (2022). International studbook for Western Lowland Gorilla (Gorilla gorilla gorilla). 2022 edition. Zoo Frankfurt, Germany.

Maisels, F., Strindberg, S., Breuer, T., Greer, D., Jeffery, K. & Stokes, E. 2018. Gorilla gorilla ssp. gorilla (amended version of 2016 assessment). The IUCN Red List of Threatened Species 2018

Malinsky, Milan, Michael Matschiner, and Hannes Svardal. ‘Dsuite - Fast *D* -statistics and Related Admixture Evidence from VCF Files’. Molecular Ecology Resources 21, no. 2 (February 2021): 584–95. 10.1111/1755-0998.13265.

McKenna, Aaron, Matthew Hanna, Eric Banks, Andrey Sivachenko, Kristian Cibulskis, Andrew Kernytsky, Kiran Garimella, et al. ‘The Genome Analysis Toolkit: A MapReduce Framework for Analyzing next-Generation DNA Sequencing Data’. Genome Research 20, no. 9 (September 2010): 1297–1303. 10.1101/gr.107524.110.

McManus, Kimberly F., Joanna L. Kelley, Shiya Song, Krishna R. Veeramah, August E. Woerner, Laurie S. Stevison, Oliver A. Ryder, et al. ‘Inference of Gorilla Demographic and Selective History from Whole-Genome Sequence Data’. Molecular Biology and Evolution 32, no. 3 (March 2015): 600–612. 10.1093/molbev/msu394.

Meisner, Jonas, and Anders Albrechtsen. ‘Inferring Population Structure and Admixture Proportions in Low-Depth NGS Data’. Genetics 210, no. 2 (October 2018): 719–31. 10.1534/genetics.118.301336.

Michel, Alice, Riana Minocher, Peter-Philip Niehoff, Yuhong Li, Kevin Nota, Maya A. Gadhvi, Jiancheng Su, Neetha Iyer, Amy Porter, Urbain Ngobobo-As-Ibungu, Escobar Binyinyi, Radar Nishuli Pekeyake, Laura Parducci, Damien Caillaud, and Katerina Guschanski. 2023. “Isolated Grauer’s Gorilla Populations Differ in Diet and Gut Microbiome.” Molecular Ecology 32, no. 23 (December): 6523–6542. 10.1111/mec.16663

Morgan, Bethan J., Chris Wild, and Atanga Ekobo. ‘Newly Discovered Gorilla Population in the Ebo Forest, Littoral Province, Cameroon’. International Journal of Primatology 24, no. 5 (1 October 2003): 1129–37. 10.1023/A:1026288531361.

Oklander, Luciana Inés, and Iván Darío Soto-Calderón. ‘Applications of Primate Genetics for Conservation and Management’. Annual Review of Anthropology 53, no. 1 (21 October 2024): 371–95. 10.1146/annurev-anthro-041422-114003.

Orkin, Joseph D., Michael J. Montague, Daniela Tejada-Martinez, Marc De Manuel, Javier Del Campo, Saul Cheves Hernandez, Anthony Di Fiore, et al. ‘The Genomics of Ecological Flexibility, Large Brains, and Long Lives in Capuchin Monkeys Revealed with fecalFACS’. Proceedings of the National Academy of Sciences 118, no. 7 (16 February 2021): e2010632118. 10.1073/pnas.2010632118.

Ostridge, Harrison J., Claudia Fontsere, Esther Lizano, Daniela C. Soto, Joshua M. Schmidt, Vrishti Saxena, Marina Alvarez-Estape, et al. ‘Local Genetic Adaptation to Habitat in Wild Chimpanzees’. Science 387, no. 6730 (10 January 2025): eadn7954. 10.1126/science.adn7954.

Pawar, Harvinder, Aigerim Rymbekova, Sebastian Cuadros-Espinoza, Xin Huang, Marc de Manuel, Tom van der Valk, Irene Lobon, et al. ‘Ghost Admixture in Eastern Gorillas’. Nature Ecology & Evolution 7, no. 9 (September 2023): 1503–14. 10.1038/s41559-023-02145-2.

Pedersen, Brent S., and Aaron R. Quinlan. ‘Mosdepth: Quick Coverage Calculation for Genomes and Exomes’. Bioinformatics (Oxford, England) 34, no. 5 (1 March 2018): 867–68. 10.1093/bioinformatics/btx699.

Perry, George H., John C. Marioni, Páll Melsted, and Yoav Gilad. ‘Genomic-Scale Capture and Sequencing of Endogenous DNA from Feces’. Molecular Ecology 19, no. 24 (December 2010): 5332–44. 10.1111/j.1365-294X.2010.04888.x.

Peter, Benjamin M. ‘100,000 Years of Gene Flow between Neandertals and Denisovans in the Altai Mountains’. bioRxiv, 15 March 2020. 10.1101/2020.03.13.990523.

Petkova, Desislava, John Novembre, and Matthew Stephens. ‘Visualizing Spatial Population Structure with Estimated Effective Migration Surfaces’. Nature Genetics 48, no. 1 (January 2016): 94–100. 10.1038/ng.3464.

Plumptre, A., Nixon, S., Caillaud, D., Hall, J.S., Hart, J.A., Nishuli, R. & Williamson, E.A. 2016. Gorilla beringei ssp. graueri. The IUCN Red List of Threatened Species 2016: e.T39995A102328430. 10.2305/IUCN.UK.2016-2.RLTS.T39995A17989838.en

Prado-Martinez, Javier, Peter H. Sudmant, Jeffrey M. Kidd, Heng Li, Joanna L. Kelley, Belen Lorente-Galdos, Krishna R. Veeramah, et al. ‘Great Ape Genetic Diversity and Population History’. Nature 499, no. 7459 (July 2013): 471–75. 10.1038/nature12228.

Prado-Martinez, Javier, Irene Hernando-Herraez, Belen Lorente-Galdos, Marc Dabad, Oscar Ramirez, Carlos Baeza-Delgado, Carlos Morcillo-Suarez, et al. ‘The Genome Sequencing of an Albino Western Lowland Gorilla Reveals Inbreeding in the Wild’. BMC Genomics 14, no. 1 (31 May 2013): 363. 10.1186/1471-2164-14-363.

Quinlan, Aaron R., and Ira M. Hall. ‘BEDTools: A Flexible Suite of Utilities for Comparing Genomic Features’. Bioinformatics 26, no. 6 (15 March 2010): 841–42. 10.1093/bioinformatics/btq033.

Robbins, M. M. ‘Male Mating Patterns in Wild Multimale Mountain Gorilla Groups’. Animal Behaviour 57, no. 5 (May 1999): 1013–20. 10.1006/anbe.1998.1063.

Robbins, Martha M., Magdelena Bermejo, Chloé Cipolletta, Florence Magliocca, Richard J. Parnell, and Emma Stokes. ‘Social Structure and Life-history Patterns in Western Gorillas (*Gorilla Gorilla Gorilla*)’. American Journal of Primatology 64, no. 2 (October 2004): 145–59. 10.1002/ajp.20069.

Robbins, Martha M., Markye Gray, Katie A. Fawcett, Felicia B. Nutter, Prosper Uwingeli, Innocent Mburanumwe, Edwin Kagoda, et al. ‘Extreme Conservation Leads to Recovery of the Virunga Mountain Gorillas’. PLOS ONE 6, no. 6 (8 June 2011): e19788. 10.1371/journal.pone.0019788.

Scally, Aylwyn, Julien Y. Dutheil, LaDeana W. Hillier, Gregory E. Jordan, Ian Goodhead, Javier Herrero, Asger Hobolth, et al. ‘Insights into Hominid Evolution from the Gorilla Genome Sequence’. Nature 483, no. 7388 (7 March 2012): 169–75. 10.1038/nature10842.

Scally, Aylwyn, Bryndis Yngvadottir, Yali Xue, Qasim Ayub, Richard Durbin, and Chris Tyler-Smith. ‘A Genome-Wide Survey of Genetic Variation in Gorillas Using Reduced Representation Sequencing’. PLOS ONE 8, no. 6 (4 June 2013): e65066. 10.1371/journal.pone.0065066.

Skotte, Line, Thorfinn Sand Korneliussen, and Anders Albrechtsen. ‘Estimating Individual Admixture Proportions from Next Generation Sequencing Data’. Genetics 195, no. 3 (1 November 2013): 693–702. 10.1534/genetics.113.154138.

Strindberg, Samantha, Fiona Maisels, Elizabeth A. Williamson, Stephen Blake, Emma J. Stokes, Rostand Aba’a, Gaspard Abitsi, et al. ‘Guns, Germs, and Trees Determine Density and Distribution of Gorillas and Chimpanzees in Western Equatorial Africa’. Science Advances 4, no. 4 (6 April 2018): eaar2964. 10.1126/sciadv.aar2964.

Thalmann, O, A Fischer, F Lankester, S Paabo, and L Vigilant. ‘The Complex Evolutionary History of Gorillas: Insights from Genomic Data’. Molecular Biology and Evolution 24, no. 1 (16 October 2006): 146–58. 10.1093/molbev/msl160.

Thalmann, Olaf, Daniel Wegmann, Marie Spitzner, Mimi Arandjelovic, Katerina Guschanski, Christoph Leuenberger, Richard A. Bergl, and Linda Vigilant. ‘Historical Sampling Reveals Dramatic Demographic Changes in Western Gorilla Populations’. BMC Evolutionary Biology 11, no. 1 (1 April 2011): 85. 10.1186/1471-2148-11-85.

UNEP-WCMC and IUCN (International Union for Conservation of Nature). 2016. Gorilla gorilla (spatial data). The IUCN Red List of Threatened Species. Version 2025-1. https://www.iucnredlist.org.

Van Der Valk, Tom, David Díez-del-Molino, Tomas Marques-Bonet, Katerina Guschanski, and Love Dalén. ‘Historical Genomes Reveal the Genomic Consequences of Recent Population Decline in Eastern Gorillas’. Current Biology 29, no. 1 (January 2019): 165–170.e6. 10.1016/j.cub.2018.11.055.

Van Der Valk, T., Jensen, A., Caillaud, D. et al. Comparative genomic analyses provide new insights into evolutionary history and conservation genomics of gorillas. BMC Ecol Evo 24, 14 (2024). 10.1186/s12862-023-02195-x

White, Lauren C., Claudia Fontsere, Esther Lizano, David A. Hughes, Samuel Angedakin, Mimi Arandjelovic, Anne-Céline Granjon, et al. ‘A Roadmap for High-Throughput Sequencing Studies of Wild Animal Populations Using Noninvasive Samples and Hybridization Capture’. Molecular Ecology Resources 19, no. 3 (May 2019): 609–22. 10.1111/1755-0998.12993.

Xue, Yali, Javier Prado-Martinez, Peter H. Sudmant, Vagheesh Narasimhan, Qasim Ayub, Michal Szpak, Peter Frandsen, et al. ‘Mountain Gorilla Genomes Reveal the Impact of Long-Term Population Decline and Inbreeding’. Science 348, no. 6231 (10 April 2015): 242–45. 10.1126/science.aaa3952.

