## Supplementary information for "Non-invasive genomic sampling uncovers novel connectivities and origins of confiscated gorillas"

Sections

1. Sample collection and data generation

1.1 Capture protocol

1.2 Assessment of the performance of the target capture hybridisation

2. Quality control

2.1 Coverage and human contamination

2.2 Variant calling metrics

2.3 Relatedness

3. Population structure

3.1. Principal component analysis

3.2. Admixture

3.3. Heterozygosity

3.4 FST

4. Population connectivity and gene flow

4.1 D statistics

4.2 Fragments of shared ancestry

4.3 EEMS

5. Geolocalisation

### 1. Sample collection and data generation

Sample collection

280 non-invasive samples (265 faecal samples, 15 hair samples) were sampled from 26 sampling sites across the distribution ranges of eastern lowland gorillas, mountain gorillas and western lowland gorillas. This was possible due to extensive collaboration with researchers in the field over several years. Wild gorilla samples were collected under the research umbrellas of various programs, including the Pan African Programme: The Cultured Chimpanzee (PanAf: Uganda (Bwindi), Cameroon(Campo Ma'an, La Belgique (Dja)), Nigeria (Mbe mountains), Republic of Congo (Conkouati, Goualougo) and Gabon ( Ivindo, Loango, Lopé, and Mts de Cristal). A full list of field sites is listed in Table 1 Supplementary, and storage conditions in Table 3 Supplementary. In addition, 12 western lowland gorillas were sampled using FTA cards®, which collect, stabilize and protect nucleic acids using blood drops at room temperature (Smith & Burgoyne, 2004). The FTA cards were sampled from rescued individuals; the location where the individuals were found was available. 11 tooth samples were collected from eastern and western gorilla specimens at 5 different natural history museums (Museum Koenig Bonn, Senckenberg Forschungsinstitut und Naturmuseum Frankfurt, Haus der Natur Salzburg, Museum für Naturkunde Berlin and Naturhistorisches Museum Vienna). For 9 out of the 11 museum samples, the region of origin information was available in the metadata provided by the museum collections. Specific geographic coordinates were not available for the museum or FTA card samples, unlike for the georeferenced faecal and hair samples. In total, 303 new samples were collected for this study from 28 different sampling sites (Figure 1 Supplementary; Figure 2 Supplementary).

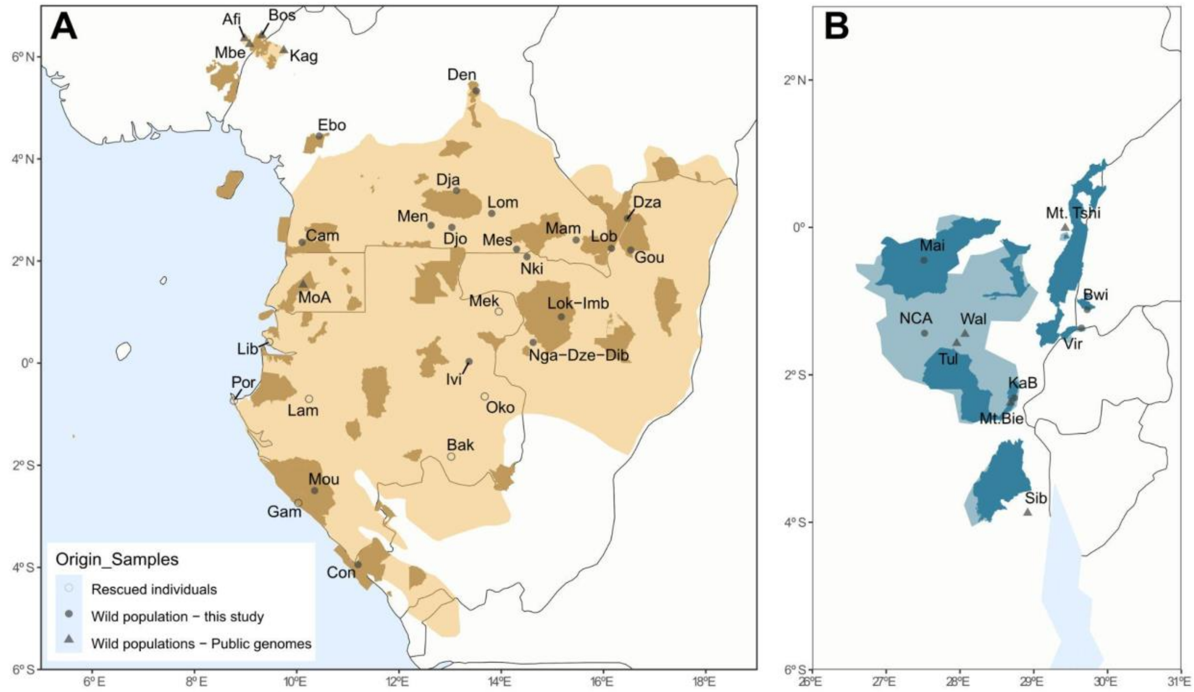


Figure 1 Supplementary. Map of the distribution ranges of the two gorilla species (A-western gorillas & B-eastern gorillas). Plotted are the sampling locations for the newly sequenced non-invasive gorilla samples and the sampling locations for previously published Cross River gorillas (Alvarez-Estape et al. 2023) and wild-born eastern gorillas (Xue et al. 2015; Pawar et al. 2023), for which we extracted chromosome 21 and the exome. Filled circles indicate sites with available geographic origin information (specific coordinates), and empty circles indicate sites where rescued gorillas were found (FTA card samples). Triangles indicate sites for which all samples came from public data. Distribution ranges are coloured yellow for western gorillas and blue for eastern gorillas. Each sampling site is labelled with an identifier, and the full names are given in Tables 1 and 3 Supplementary. National parks are highlighted in dark colors for both gorilla species.

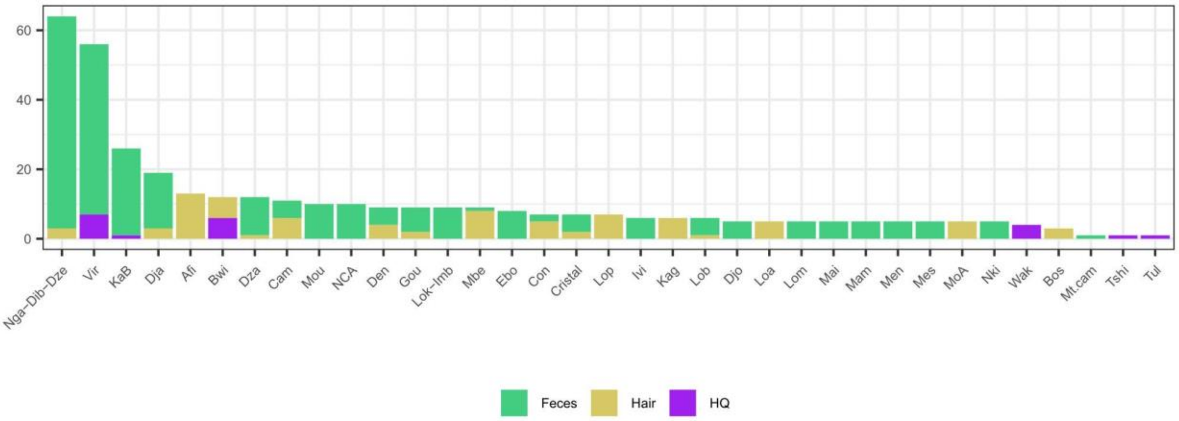


Figure 2 Supplementary. Number of samples per site: public data and collected non-invasive samples have been used in this summary plot. Each color represents a sample type or origin of the data.

| SITE | AREA<br>(National Park/Reserve) | Country | Number of samples | SUBSPECIES | DATA ORIGIN |
|---|---|---|---|---|---|
| Afi Mountain Wildlife Sanctuary | Afi Mountain Wildlife Sanctuary | Nigeria | 13 | Cross River Gorilla | Public data (Alvarez-Estape et al. 2023) |
| Bwindi | Bwindi Impenetrable Forest | Uganda | 9 | Mountain Gorilla | This study & Public data (Pawar et al. 2023) |
| Campo | Campo Ma'an National Park | Cameroon | 6 | Western Lowland Gorilla | This study |
| Conkouati | Conkouati-Douli National Park | Republic of Congo | 3 | Western Lowland Gorilla | This study & Public data (Fontsere et al. 2022) |
| Boshi Extension | Cross River National Park | Nigeria | 3 | Cross River Gorilla | Public data (Alvarez-Estape et al. 2023) |
| Deng Deng | Deng Deng National Park | Cameroon | 9 | Western Lowland Gorilla | This study & Public data (Alvarez-Estape et al. 2023) |
| Diba - Odzala National Park | Odzala National Park | Republic of Congo | 2 | Western Lowland Gorilla | This study |
| La Belgique | Dja Faunal Reserve | Cameroon | 18 | Western Lowland Gorilla | This study & Public data (Fontsere et al. 2022) |
| Djoum | Dja-Et-Lobo departament | Cameroon | 5 | Western Lowland Gorilla | This study |
| Dzanga-Ndoki | Dzanga-Sangha National Park | Central African Republic | 12 | Western Lowland Gorilla | This study & Public data (Alvarez-Estape et al. 2023) |
| Dzébé - Odzala National Park | Odzala National Park | Republic of Congo | 12 | Western Lowland Gorilla | This study |
| Ebo Forest | Ebo Wildlife Reserve | Cameroon | 8 | Western Gorilla species | This study |
| Goualougo | Nouabalé-Ndoki National Park | Republic of Congo | 9 | Western Lowland Gorilla | This study |
| Imbalanga - Odzala National Park | Odzala National Park | Republic of Congo | 3 | Western Lowland Gorilla | This study |
| Ivindo | Ivindo National Park | Gabon | 6 | Western Lowland Gorilla | This study |
| Kagwene Gorilla Sanctuary | Kagwene Gorilla Sanctuary | Cameroon | 6 | Cross River Gorilla | Public data (Alvarez-Estape et al. 2023) |
| Kahuzi-Biega NP | Kahuzi-Biega National Park | Democratic Republic of Congo | 26 | Eastern Lowland Gorilla | This study & Public data (Xue et al. 2015) |
| Lobéké | Lobéké National Park | Cameroon | 6 | Western Lowland Gorilla | This study & Public data (Alvarez-Estape et al. 2023) |
| Lokoué - Odzala National Park | Odzala National Park | Republic of Congo | 6 | Western Lowland Gorilla | This study |
| Lomié | periphery of the Dja Faunal Reserve | Cameroon | 5 | Western Lowland Gorilla | This study |
| Lope | Lopé National Park | Gabon | 1 | Western Lowland Gorilla | This study |
| Maiko | Maiko National Park | Democratic Republic of Congo | 5 | Eastern Lowland Gorilla | This study |
| Mambele | around Lobéké National Park | Cameroon | 5 | Western Lowland Gorilla | This study |
| Mbe Mountains | Mbe Mountains Community Forest | Nigeria | 5 | Cross River Gorilla | This study & Public data (Alvarez-Estape et al. 2023 |
| Mengamé | The Mengamé Gorilla Sanctuary | Cameroon | 5 | Western Lowland Gorilla | This study |
| Messok | Messok Dja Forest | Cameroon | 5 | Western Lowland Gorilla | This study |
| Monte Alen | Monte Alén National Park | Equatorial Guinea | 5 | Western Lowland Gorilla | Public data (Alvarez-Estape et al. 2023) |
| Monts de Cristal | Crystal Mountains National Park | Gabon | 5 | Western Lowland Gorilla | This study |
| Moukalaba-Doudau NP | Moukalaba-Doudau National Park | Gabon | 10 | Western Lowland Gorilla | This study |
| Mount Tshiaberimu | Virunga National Park | Democratic Republic of Congo | 1 | Eastern Lowland Gorilla | Public data (Pawar et al. 2023) |
| Ngaga Camp | Odzala National Park | Republic of Congo | 50 | Western Lowland Gorilla | This study & Public data (Alvarez-Estape et al. 2023) |
| Nki | Nki National Park | Cameroon | 5 | Western Lowland Gorilla | This study |
| Nkuba Conservation Area | Nkuba Conservation Area | Democratic Republic of Congo | 10 | Eastern Lowland Gorilla | This study |
| Tulakwa | North Kivu Province | Democratic Republic of Congo | 1 | Eastern Lowland Gorilla | Public data (Prado-Martinez et al. 2013) |
| Virunga | Virunga National Park | Rwanda | 56 | Mountain Gorilla | This study & Public data (Xue et al. 2015) |
| Walikale | North Kivu Province | Democratic Republic of Congo | 4 | Eastern Lowland Gorilla | Public data (Xue et al. 2015) |
| Museum samples- unknown real origin | - | | 8 | Western Lowland Gorilla & Eastern Lowland Gorilla | This study |
| Mamfe (musuem sample) | Near Mamfe city | Cameroon | 1 | Cross River Gorilla | This study |
| Sibatwa (museum sample) | Itombwe Nature Reserve | Democratic Republic of Congo | 1 | Eastern Lowland Gorilla | This study |
| Rescued gorillas - unknown real origin | - | - | 12 | Western Lowland Gorilla | This study |
| Public genomes unknown origin | - | - | 26 | Western Lowland Gorilla | This study & Public data (Prado-Martinez et al. 2013) |

Table 1 Supplementary. Summary sampling sites included in the project, indicating new sampling sites and public data included.

DNA extraction and library preparation were performed for the 303 newly collected samples (explained in the methods section). Library preparation failed for 19 faecal samples; therefore, these samples were discarded from the final dataset. One capture pool also failed and was removed from the final dataset.

Eight samples from the Ebo forest were collected and processed following the same procedure used for the general dataset. It was only possible to extract DNA, prepare libraries and capture Chr21 and exome for four of the initial samples.

### 1.1 Capture protocol

The capture protocol applied here followed the methodology proposed in Fontsere et al. 2021b. To determine the pooling strategy for capture, we quantified the endogenous DNA (host DNA, hDNA) of each sample, grouping the libraries in different pools for shallow sequencing (~3GB/pool). The percentage of endogenous DNA content is the gorilla endogenous DNA concentration divided by total DNA concentration (Fontsere et al. 2021b). Samples with less than 0.02 of endogenous DNA content were not expected to be successfully captured (Hernandez-Rodriguez et al. 2018; Fontsere et al. 2021b). Still, some pools having less than 0.02 of endogenous content were captured as some samples were from interesting sampling sites where few samples had been obtained, such as Ebo forest, Maiko and Nkuba Conservation Area. Pooling was performed equi-endogenously to avoid capturing bias, where the sample with the highest endogenous DNA content included in a pool at a maximum can be double that of the library with the lowest endogenous DNA, following (Fontsere et al. 2021b). 4 µg of DNA were required per pool as starting material for the target capture hybridisation as described in Fontsere et al. 2021b. We pooled 297 libraries into 21 pools (Table 2 Supplementary). Each pool was divided into two to capture the chromosome 21 and exome of the same samples, and subsequently subdivided into two replicates (a total of four divisions from one original pool). We performed two consecutive rounds of target capture hybridisation for all pools, as recommended by Hernandez-Rodriguez et al. (2018). 1 pool (11 libraries) failed the capture process and, in consequence, was not sequenced (this pool contained samples from Ebo forest, Maiko, Virunga and Nkuba Conservation Area).

Endogenous DNA

| Capture pool | n. libs | %hDNA | Gb / sample | Subproject |
|---|---|---|---|---|
| cP25 | 5 | 20,19 - 8,53 | 12 | BATCH_2 |
| cP26 | 7 | 9,75 - 4,92 | 8 | BATCH_2 |
| cP27 | 17 | 4,17 - 2,24 | 4 | BATCH_2 |
| cP28 | 30 | 2,04 - 1,10 | 4 | BATCH_1 |
| cP29 | 30 | 1,22 - 0,73 | 3 | BATCH_1 |
| cP30 | 30 | 0,74 - 0,55 | 2 | BATCH_2 |
| cP31 | 27 | 0,56 - 0,45 | 2 | BATCH_2 |
| cP32 | 32 | 0,48 - 0,35 | 2 | BATCH_2 |
| cP33 | 29 | 0,33 - 0,24 | 2 | BATCH_2 |
| cP34 | 22 | 0,25 - 0,16 | 2 | BATCH_2 |
| cP35 | 19 | 0,16 - 0,12 | 2 | BATCH_3 |
| cP36 | 11 | 0,12 - 0,09 | 2 | BATCH_3 |
| cP37 | 10 | 0,09 - 0,06 | 1 | BATCH_4 |
| cP38 | 10 | 0,05 - 0,03 | 1 | BATCH_4 |
| cP39 | 10 | 0,03 - 0,02 | 1 | BATCH_4 |
| cP57 | 4 | 1,91-1,06 | 4 | BATCH_5 |
| cP58 | 5 | 0,58-0,32 | 2 | BATCH_5 |
| cP59 | 4 | 0,18-0,06 | 2 | BATCH_6 |
| cP60 | 4 | 0,04-0,01 | 1 | BATCH_6 |
| cP61 | 7 | 53,01-26,4 | 12 | BATCH_7 |
| cP62 | 8 | 21,58-9,30 | 12 | BATCH_7 |

Table 2 Supplementary. Summary table of the endogenous DNA content (%hDNA) for each pool, number of libraries (within a pool) and GB sequenced after capture. Also given are the subproject identifiers for each capture pool.

As reported by (Maeda et al. 2023; Alvarez-Estape et al. 2023; Hernandez-Rodriguez, J. et al. 2018; Fontsere et al. 2021b), we observe hair samples have higher endogenous content than faecal samples (Figure 3 Supplementary).

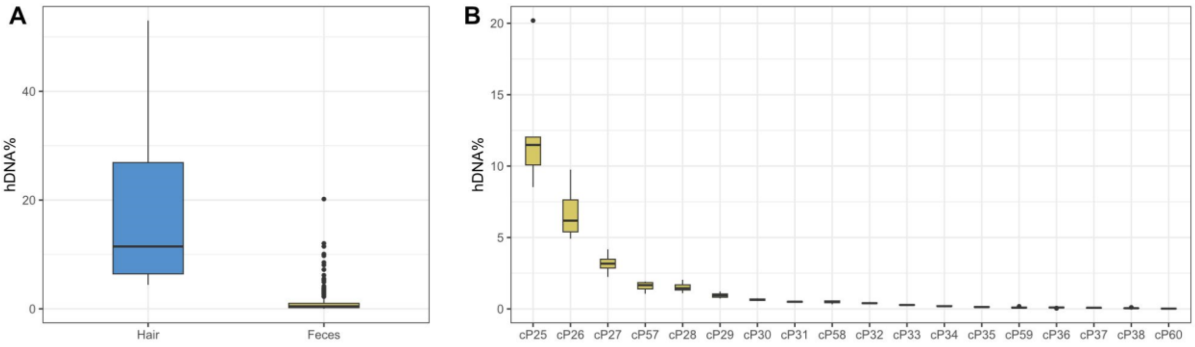


Figure 3 Supplementary. Endogenous content. A) Endogenous content per sample type (endogenous DNA is equivalent to host DNA, here hDNA). B) Endogenous content per faecal sample pool.

Chromosome 21 capture was performed using RNA baits designed by SureSelect on the chimpanzee assembly (PanTro4) and previously used in (Fontsere et al. 2022). The chromosome 21 total size design is 22.358 Mbp. For the exome capture, we also followed the capture protocol established in chimpanzees (Fontsere et al. 2021b) but using a novel capture design for the exome RNA baits, designed by SureSelect Agilent using the panTro6 assembly with 3x tiling density, which fit into three Tier 5 custom arrays. The exome total size design is 35.70 Mbp.

#### 1.2 Assessment of the performance of the target capture hybridisation

Summary metrics

Capture performance was evaluated following Fontsere et al. (2022), whereby we assessed the raw reads produced at each capture against the mapped reads (including duplicates), mapped reads without duplicates, reliable reads, and the on-target reads; the following plots are separated by sequencing batch (Figures 4-9 Supplementary). Downstream analysis will focus on the on-target reads. Reliable reads are the mapped reads without duplicates and filtered by not primary alignments. The on-target reads are the reliable reads which are inside the capture target space.

We see concordance between the summary metrics for chromosome 21 and the exome sequenced for the same sample.

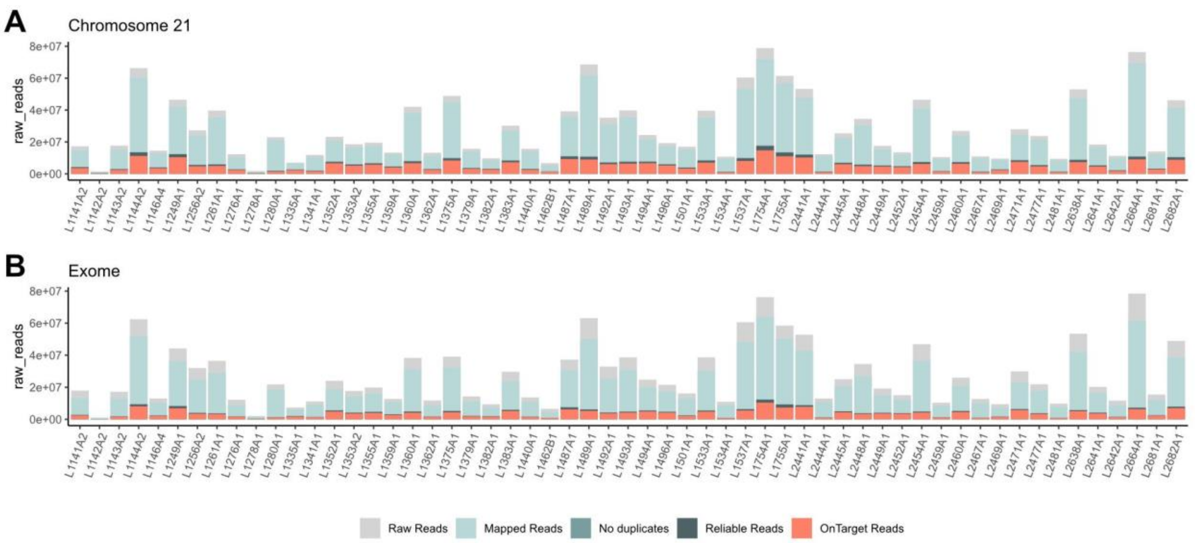


Figure 4 Supplementary. Summary metrics subproject batch 1. Summary reads per library are visualised, for chromosome 21 in panel A and for the exome in panel B. Plotted are the raw reads (grey), mapped reads with duplicates (light blue), mapped reads without duplicates (blue), reliable reads (dark blue), and on-target reads (orange).

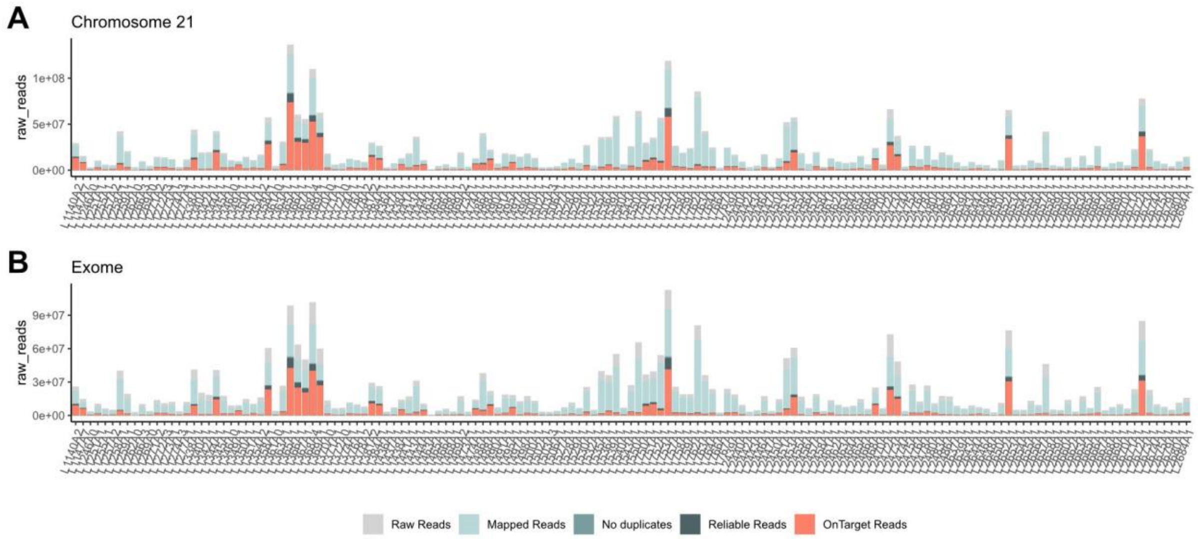


Figure 5 Supplementary. Summary metrics subproject batch 2. Summary reads per library are visualised, for chromosome 21 in panel A and for the exome in panel B. Plotted are the raw reads (grey), mapped reads with duplicates (light blue), mapped reads without duplicates (blue), reliable reads (dark blue), and on-target reads (orange).

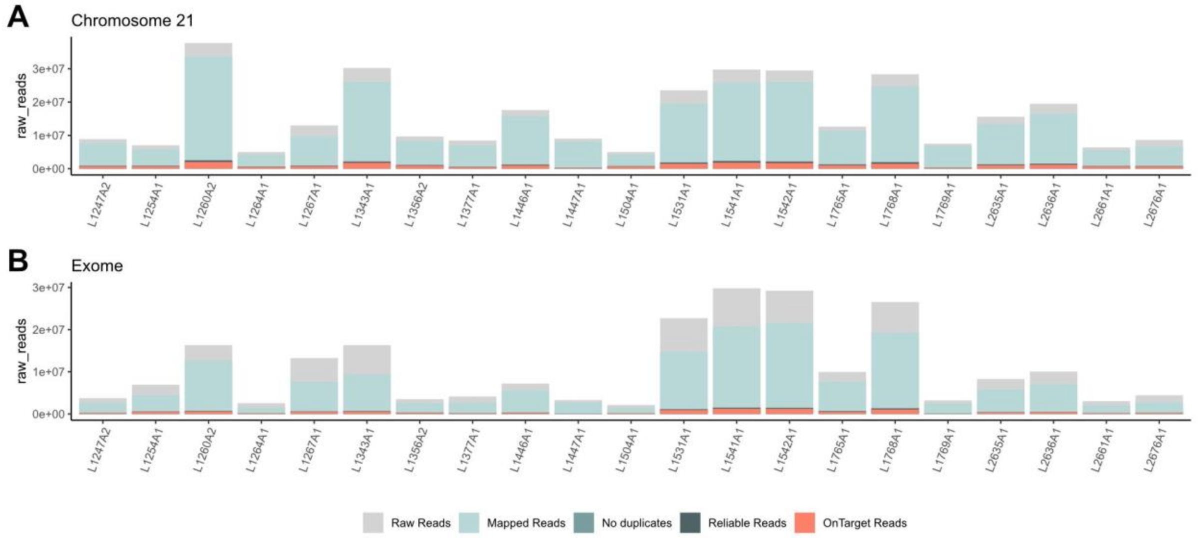


Figure 6 Supplementary. Summary metrics subproject batch 3. Summary reads per library are visualised, for chromosome 21 in panel A and for the exome in panel B. Plotted are the raw reads (grey), mapped reads with duplicates (light blue), mapped reads without duplicates (blue), reliable reads (dark blue), and on-target reads (orange).

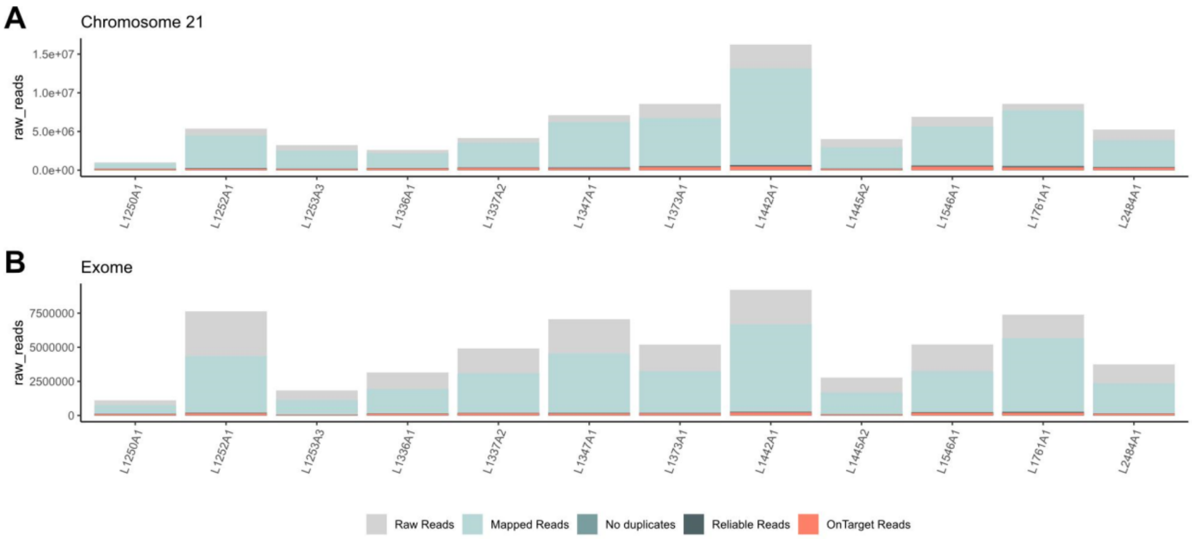


Figure 7 Supplementary. Summary metrics subproject batch 4. Summary reads per library are visualised, for chromosome 21 in panel A and for the exome in panel B. Plotted are the raw reads (grey), mapped reads with duplicates (light blue), mapped reads without duplicates (blue), reliable reads (dark blue), and on-target reads (orange).

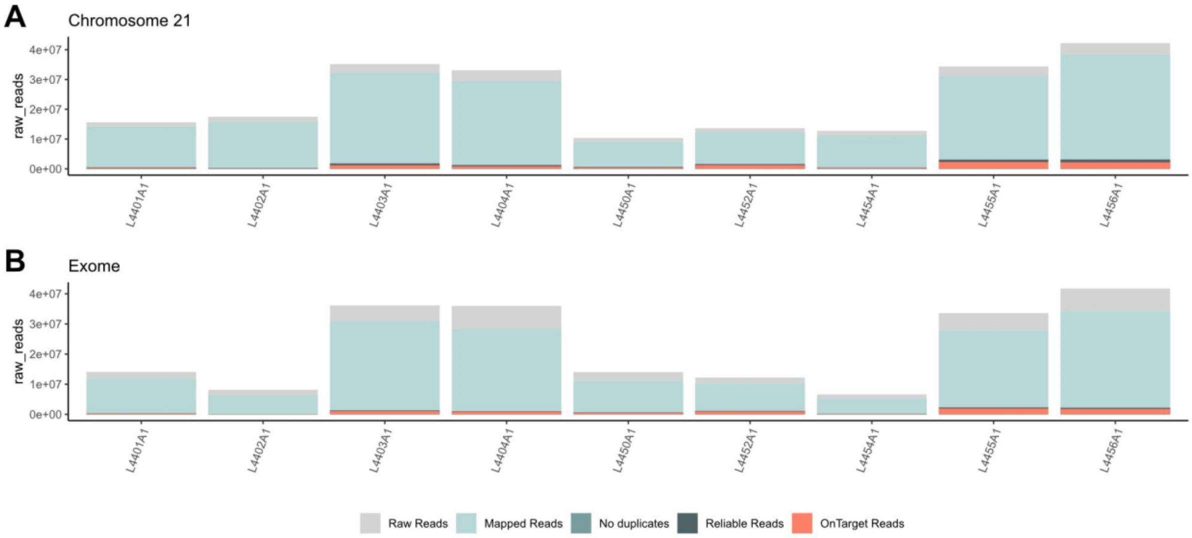


Figure 8 Supplementary. Summary metrics subproject batch 5. Summary reads per library are visualised, for chromosome 21 in panel A and for the exome in panel B. Plotted are the raw reads (grey), mapped reads with duplicates (light blue), mapped reads without duplicates (blue), reliable reads (dark blue), and on-target reads (orange).

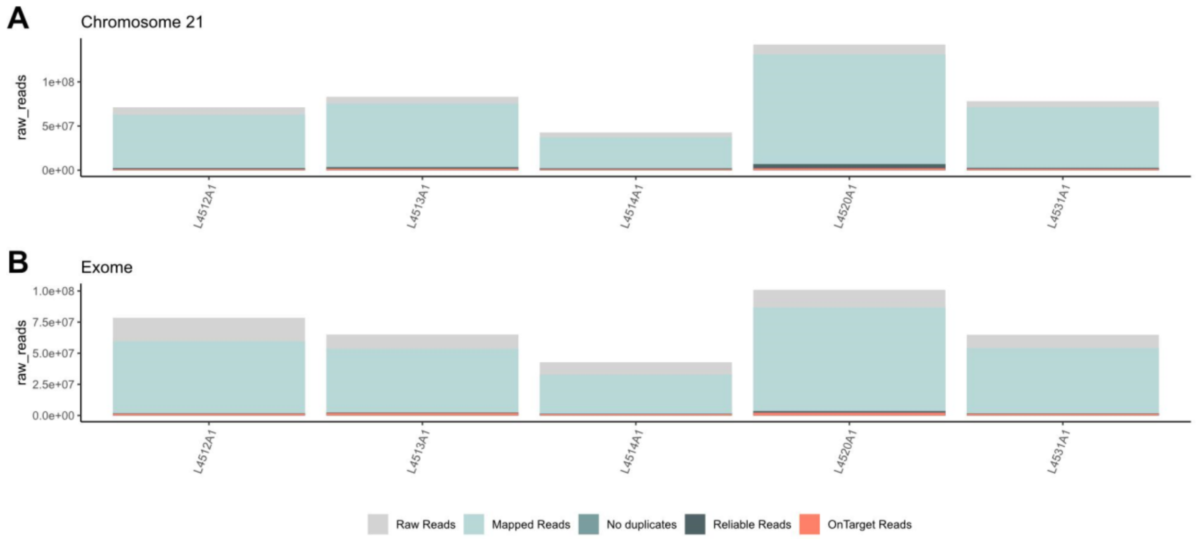


Figure 9 Supplementary. Summary metrics subproject batch 7 - hair samples. Summary reads per library are visualised, for chromosome 21 in panel A and for the exome in panel B. Plotted are the raw reads (grey), mapped reads with duplicates (light blue), mapped reads without duplicates (blue), reliable reads (dark blue), and on-target reads (orange).

We further assessed the performance of the target capture hybridisation by calculating the enrichment factor, capture specificity, library complexity and capture sensitivity, following the definitions of (Fontsere et al. 2021b; Hernandez-Rodriguez et al. 2018).

<u>Enrichment factor</u>

We evaluated the enrichment obtained for chromosome 21 and the exome of the same samples, where the enrichment factor is defined as the ratio of reads inside the capture target space to the total sequenced reads divided by the ratio of the capture target space to the whole genome size. Endogenous DNA of the sample and the enrichment factor post-capture were positively correlated, as expected from previous studies (Fontsere et al. 2021b) (Figure 10-11 Supplementary). The enrichment factor in the chromosome 21 captures was different for faecal and hair samples. Hair samples showed lower values of enrichment factor, even though these samples had higher values of endogenous DNA (Figure 10 Supplementary).

We saw higher enrichment of chromosome 21 than the exome (Figure 12A Supplementary), when using the entire designed exome space to calculate it. The lower enrichment factor in the exome data, compared to the chromosome 21 data, can be explained by the missing chromosomes (see section 2.1 for further details) in the exome capture compared to the design of the exome on-target space, as the space design is considered when calculating the enrichment factor. When calculating the enrichment factor using the real captured exome target space, the values are similar to chromosome 21 (Figure 12B Supplementary).

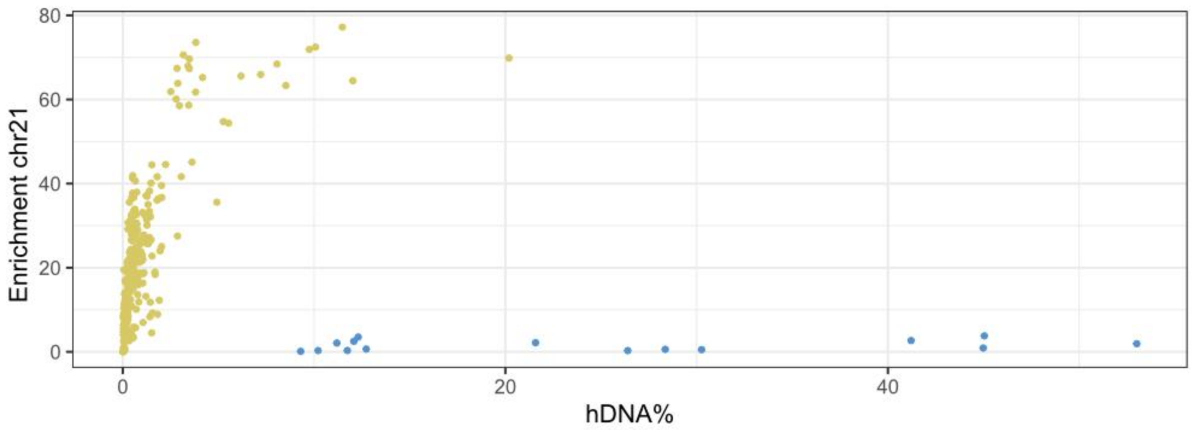


Figure 10 Supplementary. Enrichment factor chromosome 21 capture vs endogenous DNA. Each dot represents a library, and the color is the sample type (green=faeces & blue=hairs).

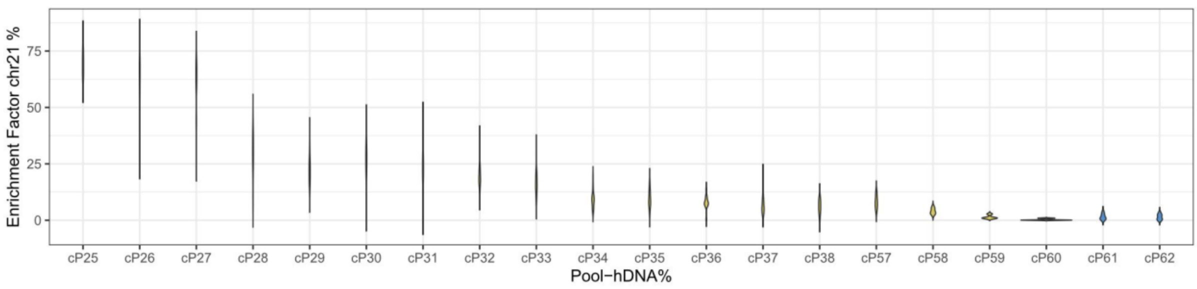


Figure 11 Supplementary. Enrichment factor of chromosome 21 by capture pool. Each color represents the sample type (green=faeces, blue=hairs). Pools are ordered by endogenous content value for faecal samples, which were pooled equi-endogenously.

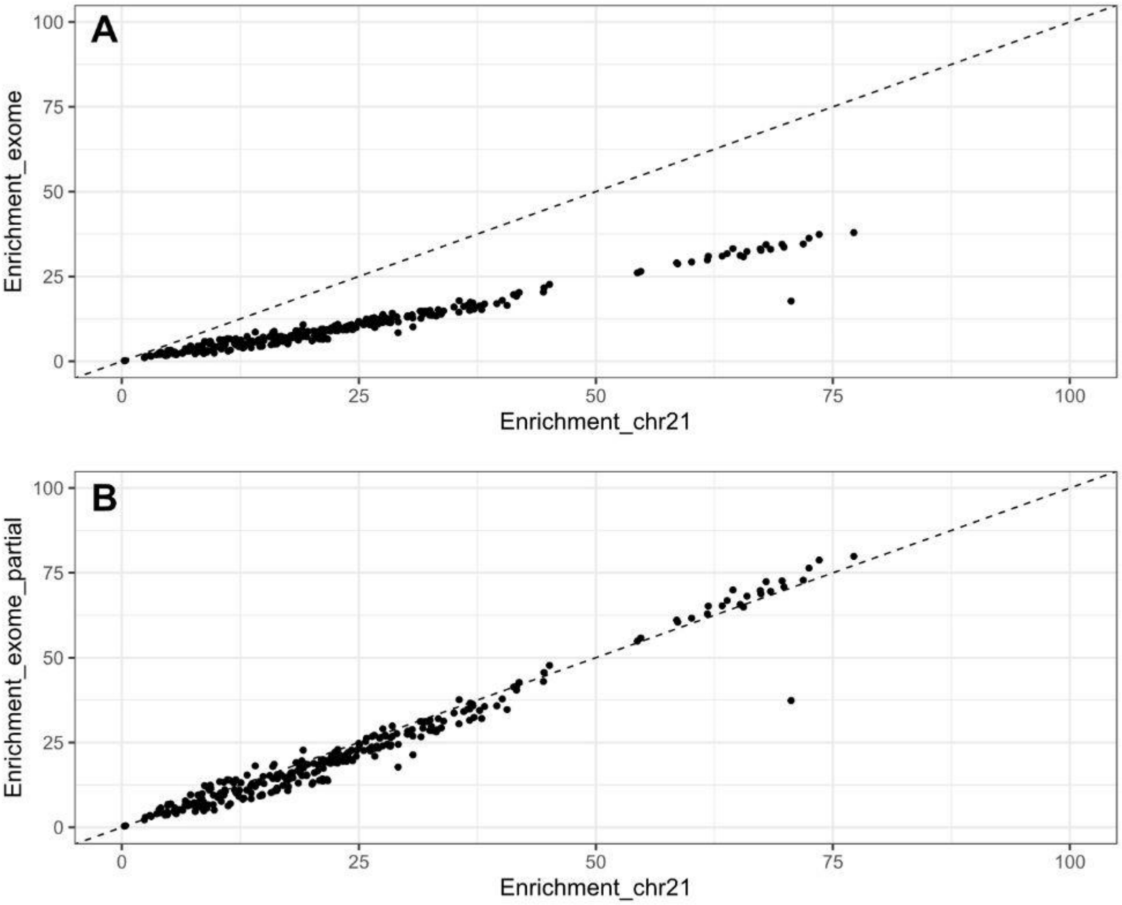


Figure 12 Supplementary. Correlation between enrichment after capture obtained for chromosome 21 versus the exome for the same library preparations. A) Enrichment factor using the designed exome target space. B) Enrichment factor using the successfully captured exome target space.

Library complexity

We assessed the complexity of the libraries produced for chromosome 21 and the exome of the same samples, where library complexity is the number of reliable reads divided by the mapped reads, including duplicates (Fontsere et al. 2021b; Hernandez-Rodriguez et al. 2018).

As expected, library complexity was higher for samples which had higher values of endogenous DNA (Figure 13-14 Supplementary), except for hair samples, which show the same poorer capture performance as seen in the enrichment factor section.

Library complexity between chromosome 21 and exome data did not differ as much as in the enrichment factor due to the fact that the size of the target space design is not considered in the measure (Figure 15 Supplementary).

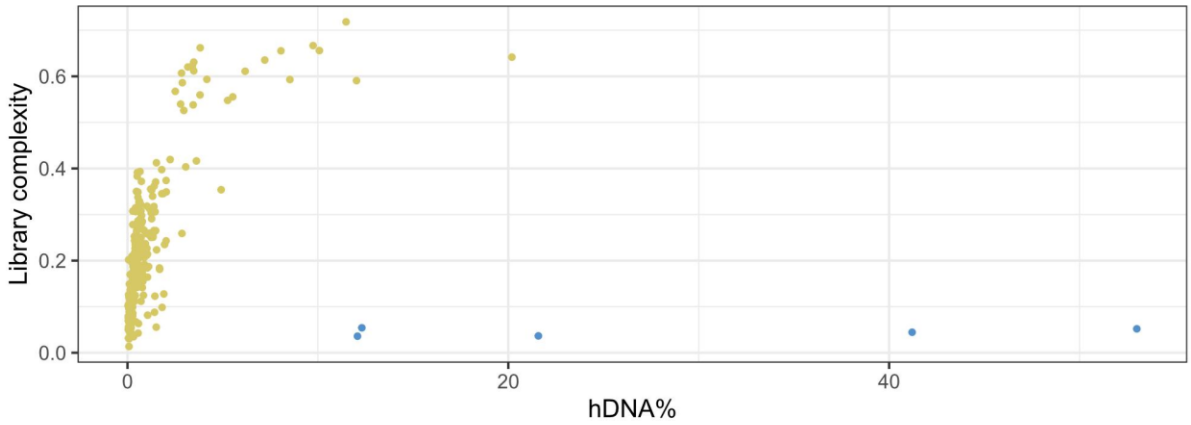


Figure 13 Supplementary. Library complexity vs endogenous DNA. Each dot represents a unique library, and each color represents the sample type (green = faeces, blue = hairs).

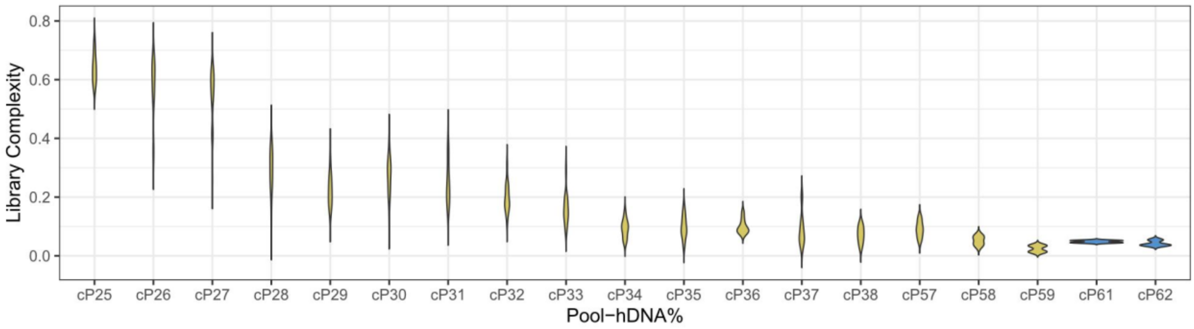


Figure 14 Supplementary. Library complexity for each capture pool. Each color represents the sample type (green = faeces, blue = hairs). Pools are ordered by endogenous content value for faecal samples, which were pooled equi-endogenously.

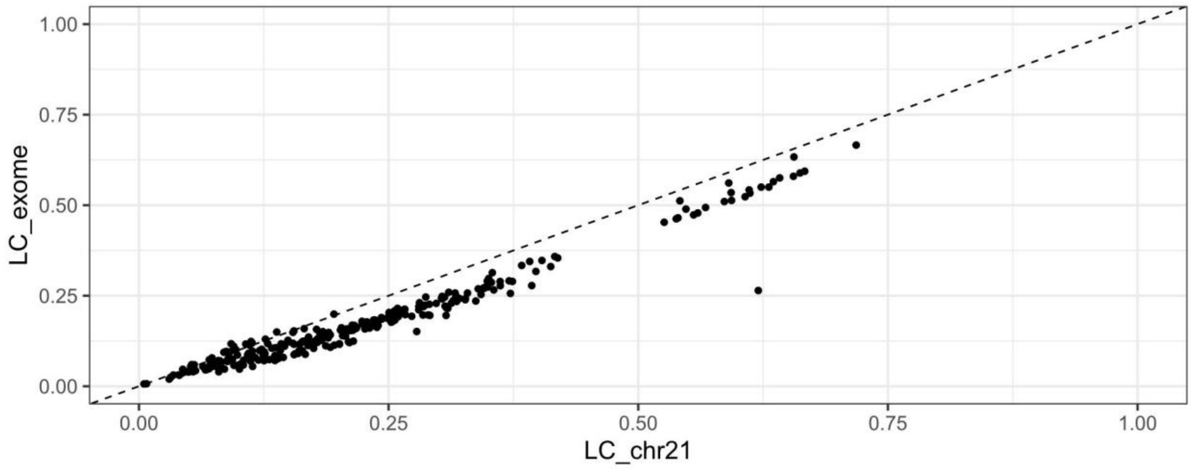


Figure 15 Supplementary. Correlation between library complexity for chromosome 21 versus the exome for the same samples (the same library preparations).

Specificity

Capture specificity is the ratio of on-target reads to reliable reads (Fontsere et al. 2021b; Hernandez-Rodriguez et al. 2018).

The specificity values were higher than 80% for the majority of samples (Figure 16 Supplementary) and capture pools (Figure 17 Supplementary), as expected from Fontsere et al. 2021b. Samples with lower endogenous content showed lower specificity values between 80-60%. Hair samples also showed lower levels of specificity, reflecting a possible capture failure of these batches. Capture specificity between chromosome 21 and exome data was similar (Figure 18 Supplementary).

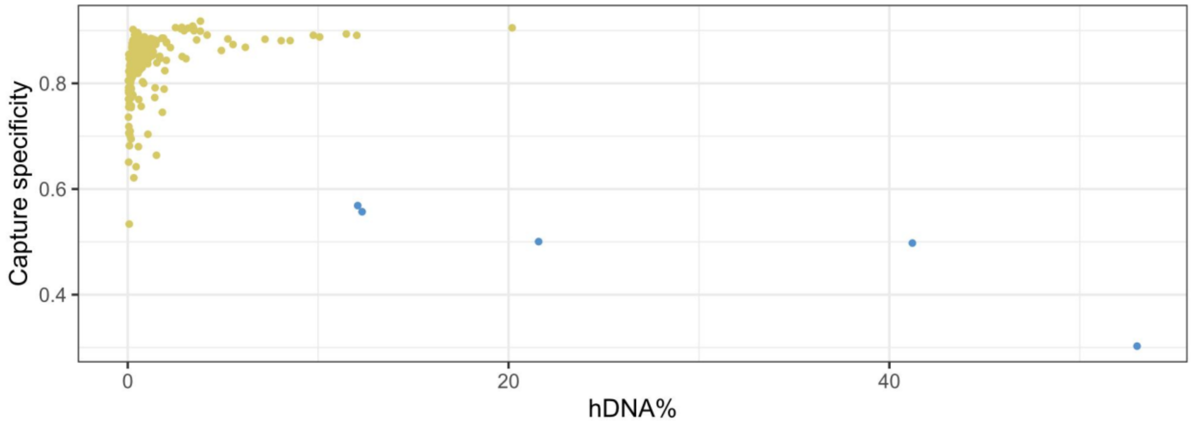


Figure 16 Supplementary. Capture specificity vs endogenous DNA. Each dot represents a unique library, and each color represents the sample type (green = faeces, blue = hairs).

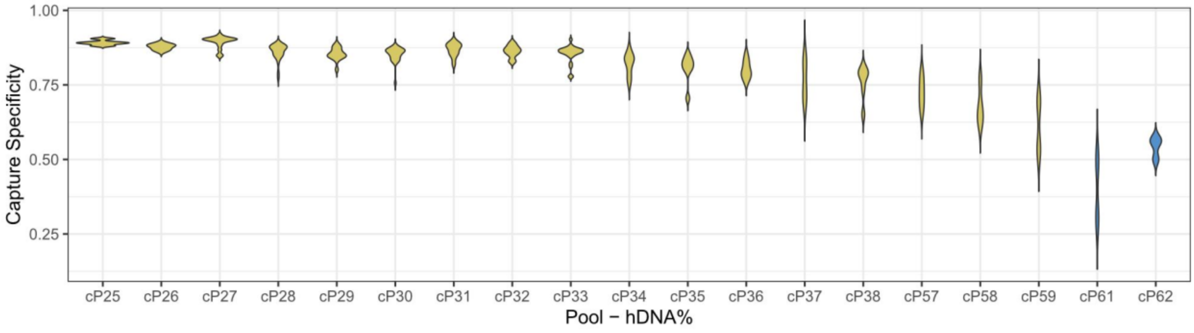


Figure 17 Supplementary. Capture specificity for each capture pool. Each color represents the sample type (green = faeces, blue = hairs). Pools are ordered by endogenous content value for faecal samples, which were pooled equi-endogenously.

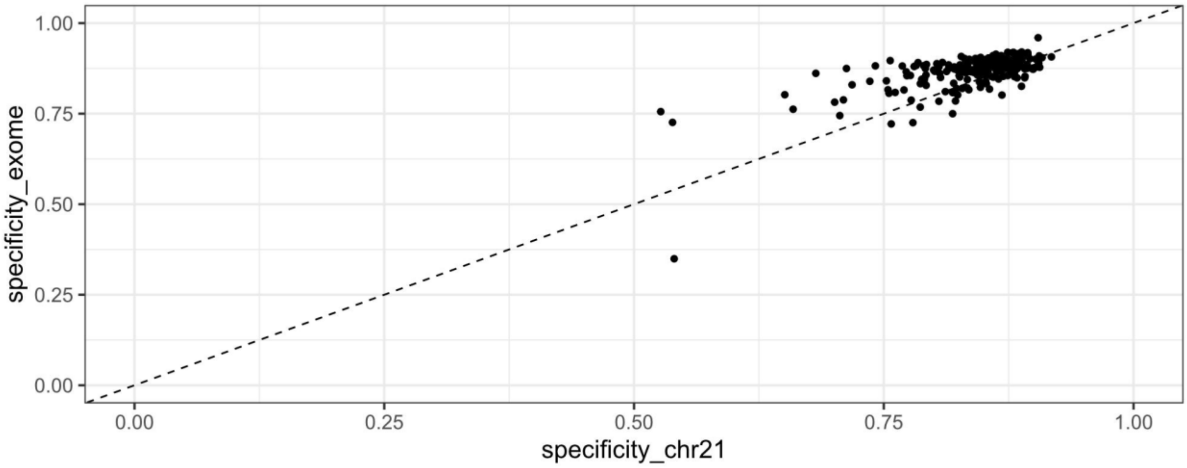


Figure 18 Supplementary. Capture specificity chromosome 21 captures vs capture specificity exome captures. Each dot represents a library.

Sensitivity

We consider capture sensitivity as the number of target regions with average coverage of at least 5 (DP5) or 50 (DP50), divided by the number of target regions (DP5 = Figure 19 Supplementary).

Capture sensitivity measures were similar when comparing 5X and 50X depth of coverage (Figure 20-21 Supplementary). The measures were also affected by the endogenous content of each sample, as expected (Figure 20-21 Supplementary).

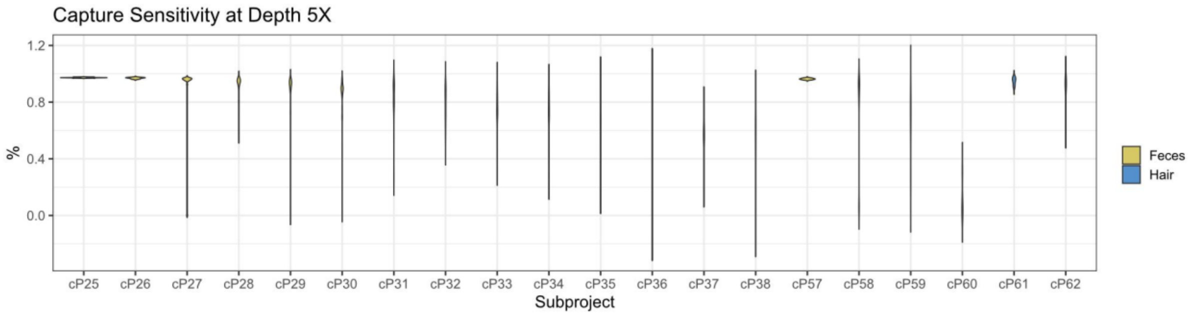


Figure 19 Supplementary. Capture sensitivity at 5DP for each capture pool. Each color represents the sample type (green = faeces, blue = hairs). Pools are ordered by endogenous content value for faecal samples, which were pooled equi-endogenously.

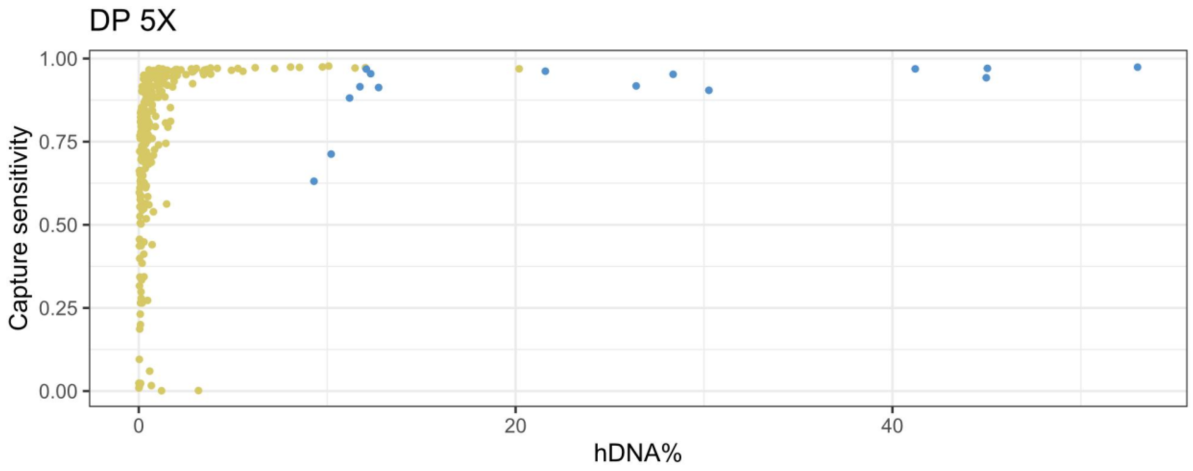


Figure 20 Supplementary. Capture sensitivity at DP5 of chromosome 21 captures vs endogenous content. Each dot represents a unique library, and each color represents the sample type (green = faeces, blue = hairs).

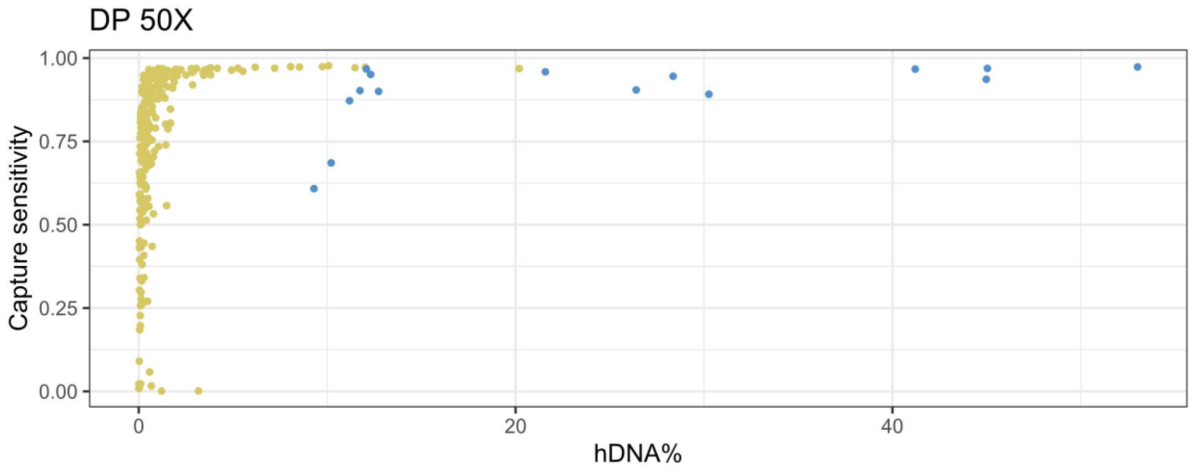


Figure 21 Supplementary. Capture sensitivity at DP50 of chromosome 21 captures vs endogenous content. Each dot represents a unique library, and each color represents the sample type (green = faeces, blue = hairs).

### 2. Quality control

#### 2.1 Coverage and human contamination

Chromosome 21 depth of coverage

We calculated the coverage in the on-target space for each of the newly sequenced non-invasive samples (Figure 22 Supplementary).

As described in methods, we removed samples with low-coverage (<0.5X coverage) and samples with more than 1% human contamination as estimated using the HuConTest (Kuhlwilm et al. 2021). Mean coverage in the on-target space varied depending on sample type due to different sequencing depths (publicly available genomes) and whether the samples had been target-captured (faecal and some hair samples) (Figure 22A Supplementary). As can be seen in Figure 22B Supplementary, all targeted regions were successfully covered.

We retained 278 chromosome 21 samples with >=0.5X average coverage and <1% human contamination. Due to its geographic uniqueness, one sample from Ebo forest (mean coverage=1.42X, human contamination=1.537%) was included in PCA, admixture, and admixfrog analysis (the rest were excluded due to low coverage and high human contamination). A total of 13 samples were removed due to low coverage.

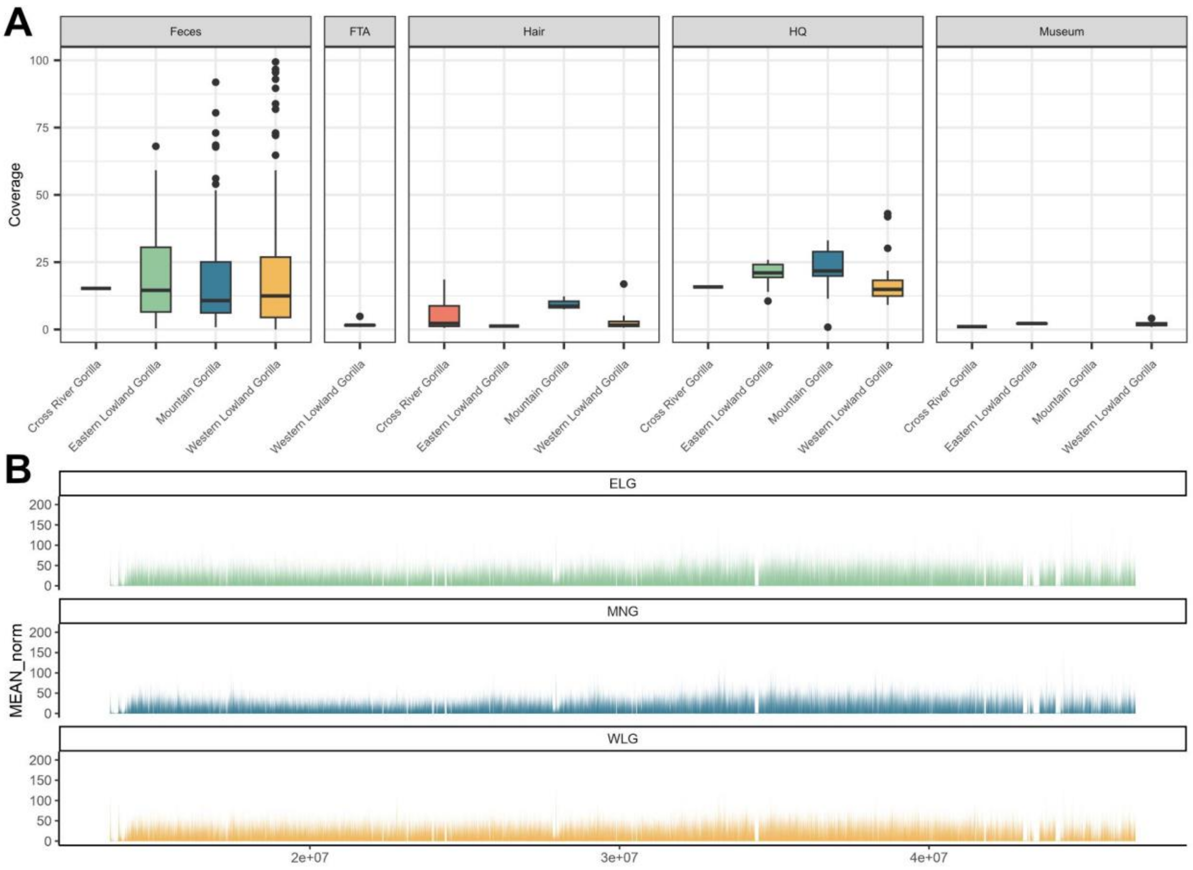


Figure 22 Supplementary. Coverage by subspecies and sample type. Each subspecies is represented by one unique color (Cross River gorillas in red, eastern lowland gorillas in green, western lowland gorillas in yellow and mountain gorillas in blue). A) Coverage for chromosome 21 samples per sampling site, including non-invasive samples for Cross River gorillas from Alvarez-Estape et al. (2023). Box plots are coloured by subspecies (green eastern lowland gorillas, blue mountain gorillas, orange western lowland gorillas, red Cross River gorillas). B) Coverage across chromosome 21 plotted per subspecies. X axis represents bases across chromosome 21.

<u>Human contamination</u>

Human contamination calculated in chromosome 21 data was equally affecting the four subspecies. 41 samples were excluded due to human contamination in chromosome 21 data (samples with less than 0.5X of coverage were previously discarded) (Figure 23 Supplementary).

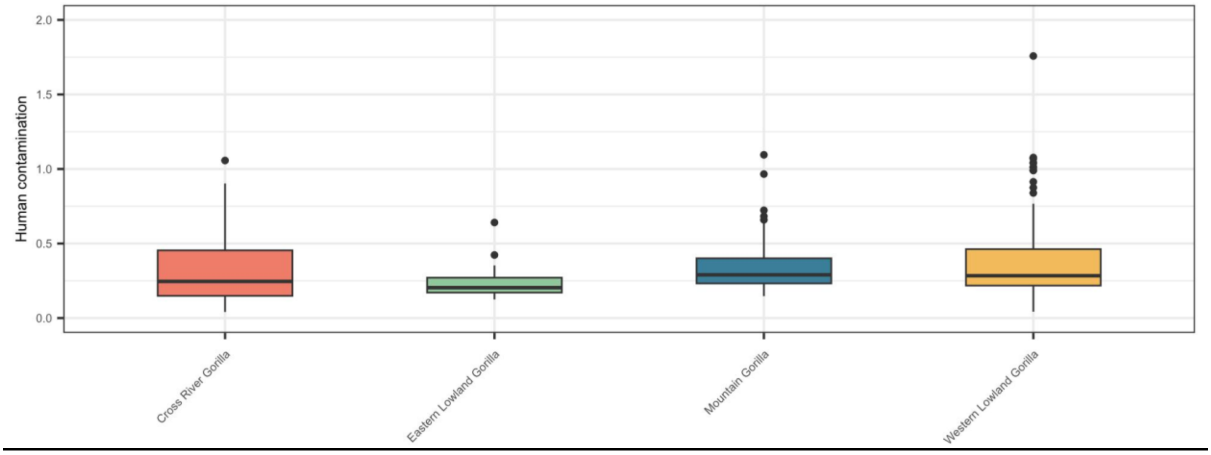


Figure 23 Supplementary. Human contamination by subspecies. Each subspecies is represented by one unique color (Cross River gorillas in red, eastern lowland gorillas in green, western lowland gorillas in yellow and mountain gorillas in blue).

<u>Exome depth of coverage</u>

We calculated coverage in the exome on-target space (Figure 24 Supplementary). We observe three coverage distributions (low, medium, high) across chromosomes for the captured exome samples, corresponding to the three sets of baits for each of the custom arrays. Unfortunately, due to a manufacturing error, one of the sets of RNA baits (containing chromosomes 1, 10-16) from the exome design was not produced and so not captured (regions with low coverage), while another set of the baits (containing chromosomes 2-9) was duplicated (regions with high coverage), causing these differences in coverage.

The exomes of chromosome X and chromosomes 2-9, 17-22 were successfully captured across the on-target exome space of the respective chromosomes. Therefore, we focus our analysis of the gorilla exome to chromosome X to an autosome comparisons (in the main), and to a general characterisation of the partial exome on-target space (representing 60.4% of genes in the exome target space) in the Supplementary.

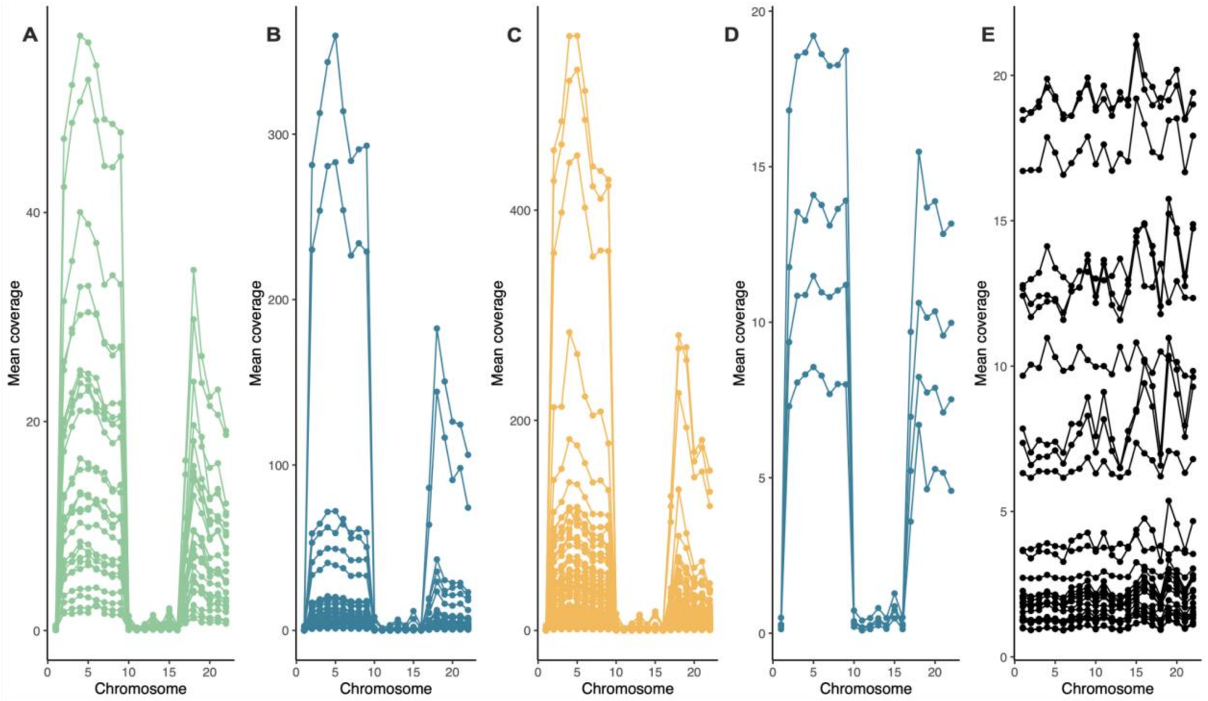


Figure 24 Supplementary. Average coverage per chromosome in the autosomal exome on-target space for captured faecal samples of ELG (A), MG (B) and WLG (C) and captured hair samples of MG (D). Also shown is the coverage across the autosomes for previously published CRG WGS samples (Alvarez-Estape et al. 2023), subset to the exome on-target space (E).

We applied the same filters to the partial exome dataset as for the chromosome 21 dataset (>=0.5X average coverage and < 1% human contamination), retaining 27 eastern lowland gorillas, 45 mountain gorillas, 132 western lowland gorillas and 16 Cross River gorillas.

### 2.2 Damage Patterns

Museum specimens often contain fragmented and chemically altered DNA due to age, environmental exposure, and storage conditions. In order to assess and visualise DNA damage patterns in the 11 museum samples, we applied the MapDamage bioinformatics tool (Jónsson et al. 2013), see (Figure 25 Supplementary; Figure 26 Supplementary.

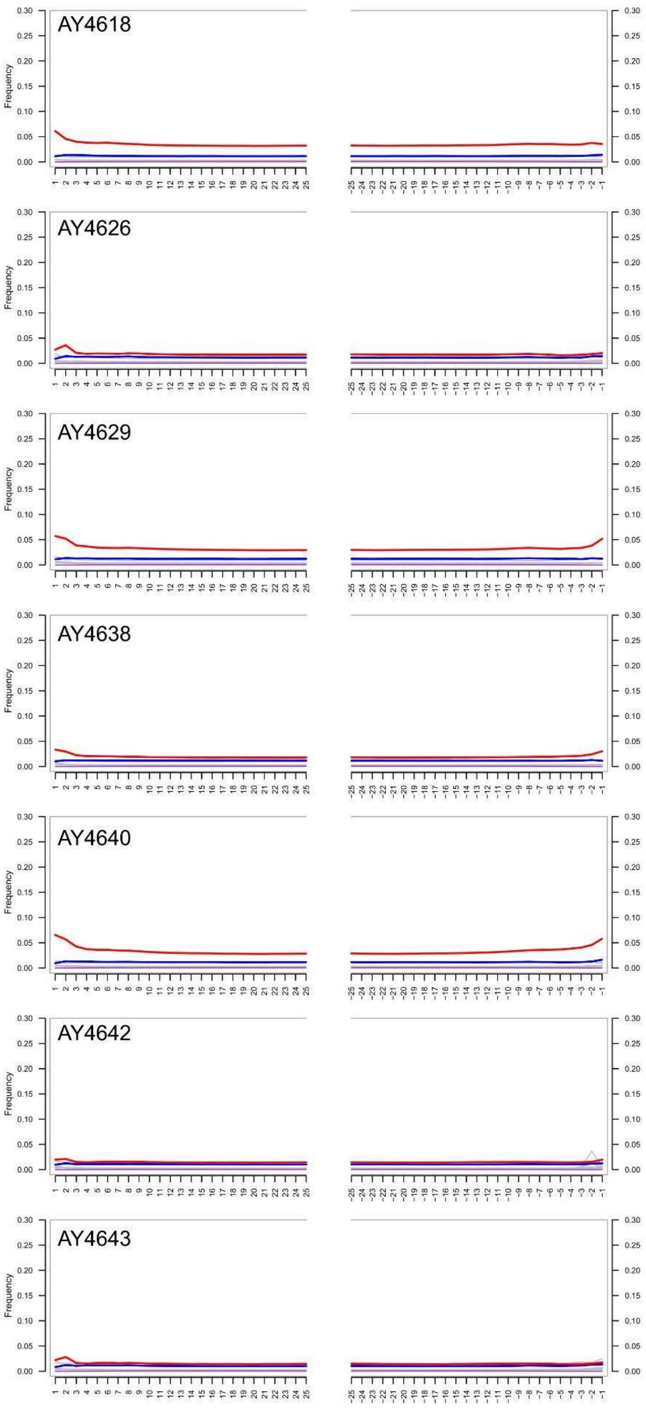


Figure 25 Supplementary. Damage patterns analysis of ancient DNA extracted from museum specimens using MapDamage for batch 5. The Fragmisincorporation plot visualizes nucleotide misincorporation patterns at the 5′ and 3′ ends of DNA fragments.

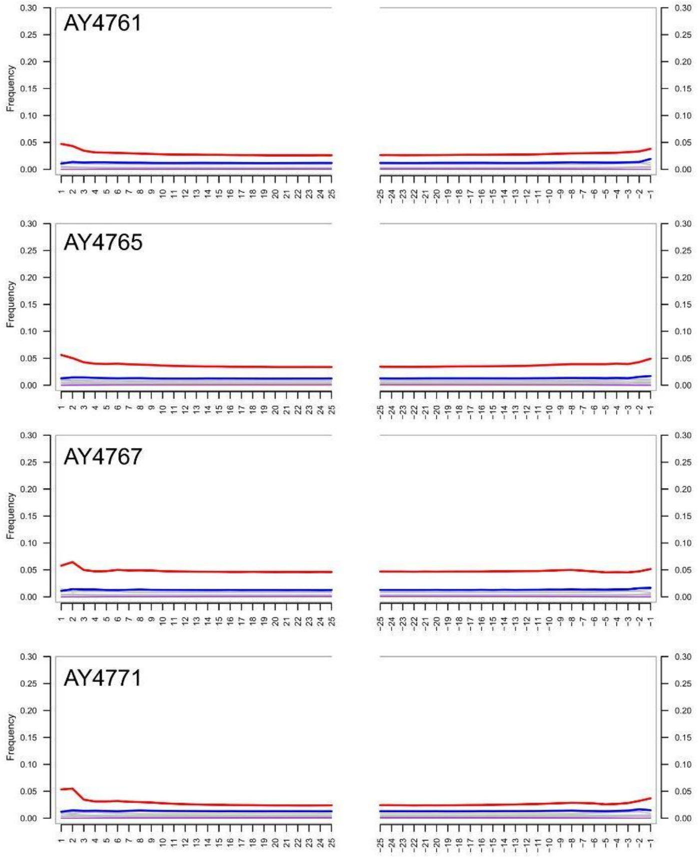


Figure 26 Supplementary. Damage patterns analysis of ancient DNA extracted from museum specimens using MapDamage for batch 6. The Fragmisincorporation plot visualizes nucleotide misincorporation patterns at the 5′ and 3′ ends of DNA fragments.

### 2.3 Relatedness

Kinship relations were assessed using ANGSD (Korneliussen et al. 2014) and NgsRelate (Korneliussen et al. 2015) per gorilla sampling site for each pair of individuals that passed our filtering criteria (coverage > 0.5X and human contamination < 1%). This analysis was run per sample site, as population-specific allele frequencies were used.

First-degree relatives, including monozygotic twins and duplicated individuals, were removed to prevent potential bias in population analyses. Coverage and the number of related pairs involved were used as the main criteria to remove individuals from each related pair. In total, 25 samples were removed due to first-degree relatedness or possible duplicates. Relatedness between pairs of individuals is represented in the plots below for all sites containing at least one third-degree relationship (Figure 27-31 Supplementary).

#### Cross River gorillas

For Cross River gorillas, where we extracted chromosome 21 of the whole genome data generated in (Alvarez-Estape et al. 2023), first and second degree relatedness results (Figure

28 Supplementary) were as expected in accordance with the related pairs found in Alvarez-Estape et al. 2023 using the whole genomes, except for one pair. Taking only the chromosome 21 on-target space, Afi20 and Afi17 were found as first-degree related pairs, in contrast with Alvarez-Estape et al. 2023 where they were described as second-degree related.

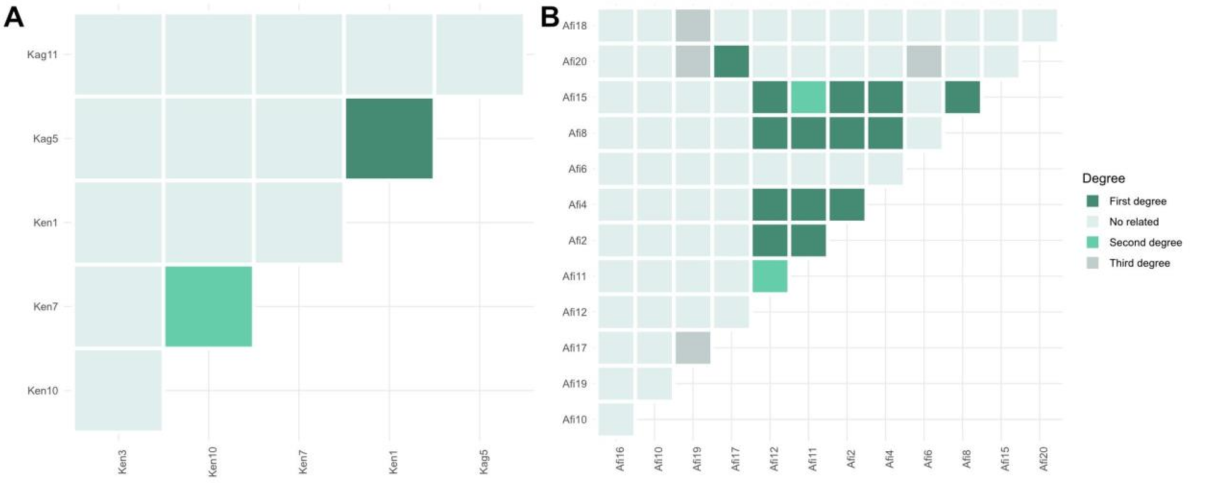


Figure 27 Supplementary. Relatedness pairs of Cross River gorilla sites. A) Kagwene Gorilla Sanctuary B) Afi Mountain Wildlife Sanctuary.

Afi Mountain Wildlife Sanctuary samples removed were: Afi17, Afi12, Afi8, Afi15, Afi11. Only one sample from Kagwene Gorilla Sanctuary was removed (Kag5). CRNP-Boshi Extension and Mbe Mountains sites did not show any relatedness between individuals.

**Eastern lowland gorillas**

Kahuzi-Biega National Park (KaB)

One sample from Kahuzi-Biega (KaB) was removed (L1546A1) due to first-degree relatedness (Figure 29 Supplementary).

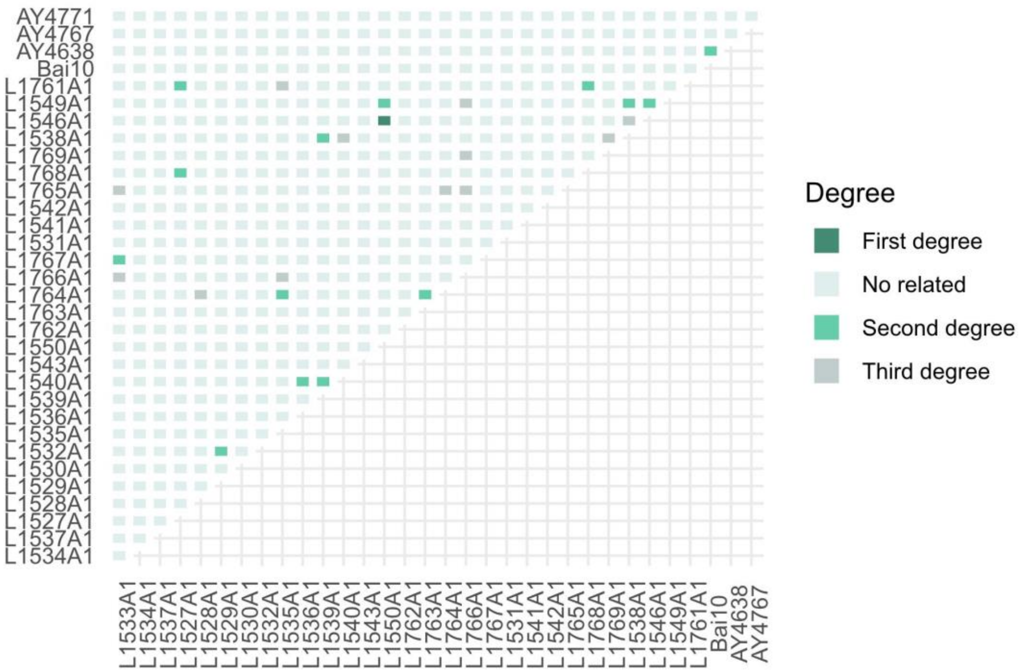


Figure 28 Supplementary. Relatedness pairs of eastern lowland gorilla sites at Kahuzi-Biega National Park.

The sampling sites, Maiko (Mai) and Nkuba Conservation Area (NCA), did not show any first, second or third-degree relatedness.

#### Mountain gorillas

From Bwindi, none of the three hair samples were related. For the Virunga site, 2 samples were removed (Figure 30 Supplementary): L2646A1, L2680A1.

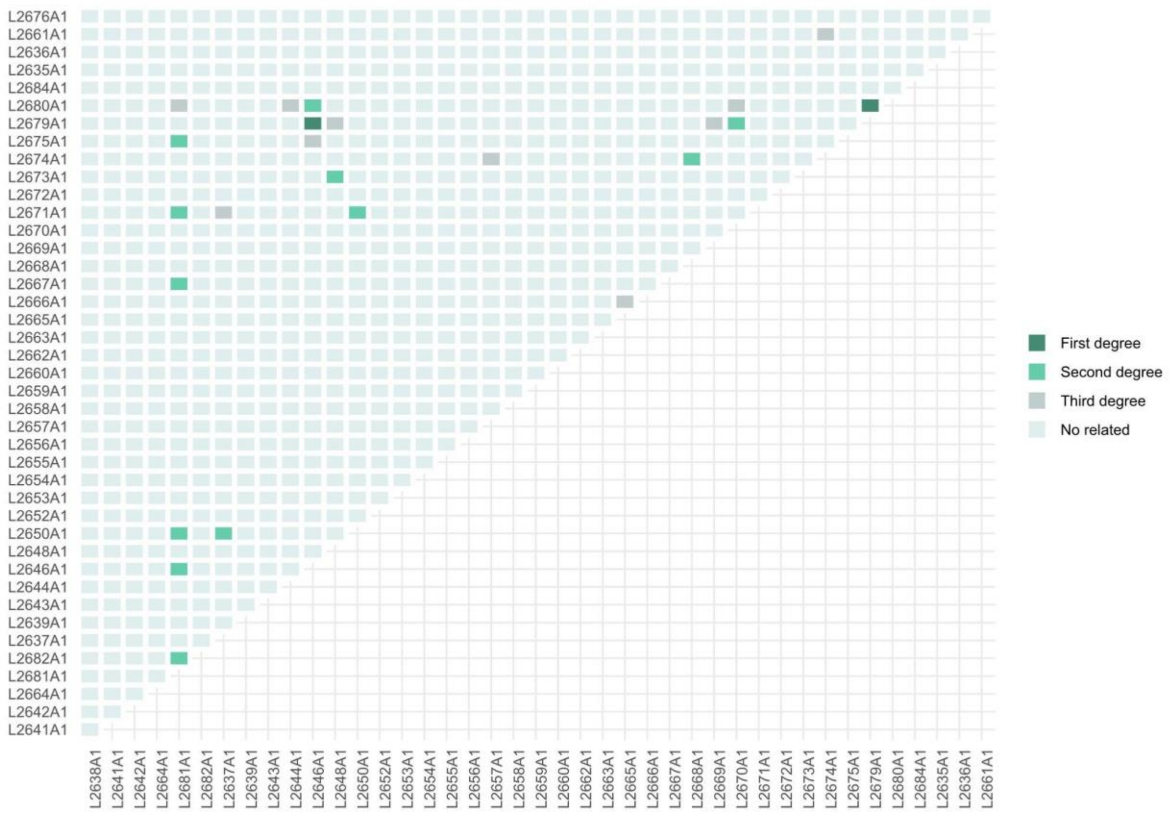


Figure 29 Supplementary. Relatedness pairs of the Virunga site - mountain gorillas.

**Western lowland gorillas**

For the western lowland gorilla subspecies, just 5 sites presented duplicated individuals or first to third-degree related pairs.

Ngaga Camp, Diba and Dzébé (Nga-Dib-Dze)

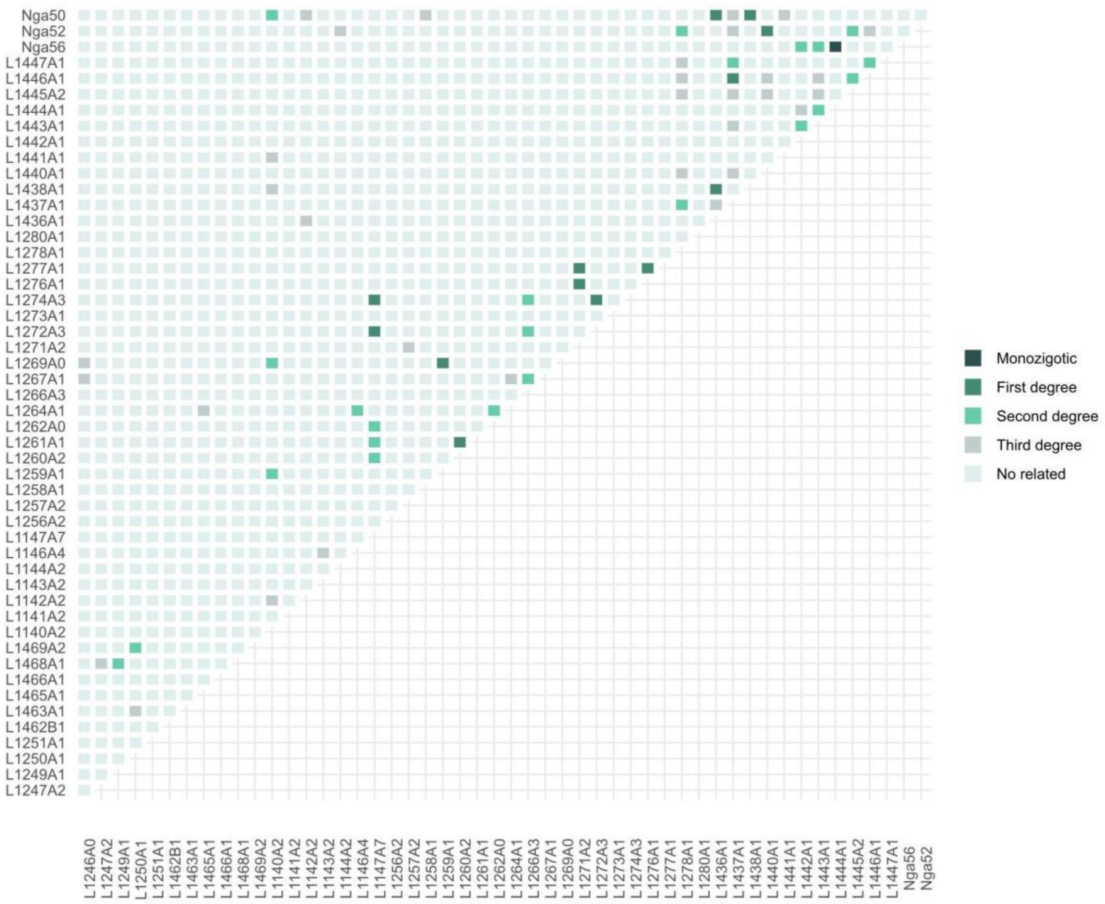


---

Figure 30 Supplementary. Relatedness pairs Ngaga camp, Diba and Dzébé western lowland gorillas.

For the Nga-Dib-Dze site, 11 samples were removed (Figure 31 Supplementary): L1272A3, L1274A3, L1269A0, L1260A2, L1276A1, L1277A1, L1436A1, Nga50, L1437A1, Nga52, Nga56.

La Belgique, Lobéké, Lomié and Mambele

We removed one sample from La Belgique (Bel); L1504A1 (Figure 32A Supplementary). No samples were removed from Lobéké (Lob); 1 second-degree relationship was found (Figure 32B Supplementary). Three samples from Lomié (Lom) were removed (Figure 32C Supplementary): L1366A1, L1367A1, L1369A. No samples were removed from Mambele (Mam); 1 third-degree relationship was found (Figure 32D Supplementary). No other relatedness pairs were found in the rest of the western lowland sites: Dzanga-Ndoki (Dza), Campo (Cam), Deng Deng (Den), Djoum (Djo), Lokoué-Imbalanga (Lok-Imb), Mengamé (Men), Monte Alen (MoA), Messok (Mes), Moukalaba-Doudau (Mou), Nki, Ivindo (Ivi) and Goualougo (Gou).

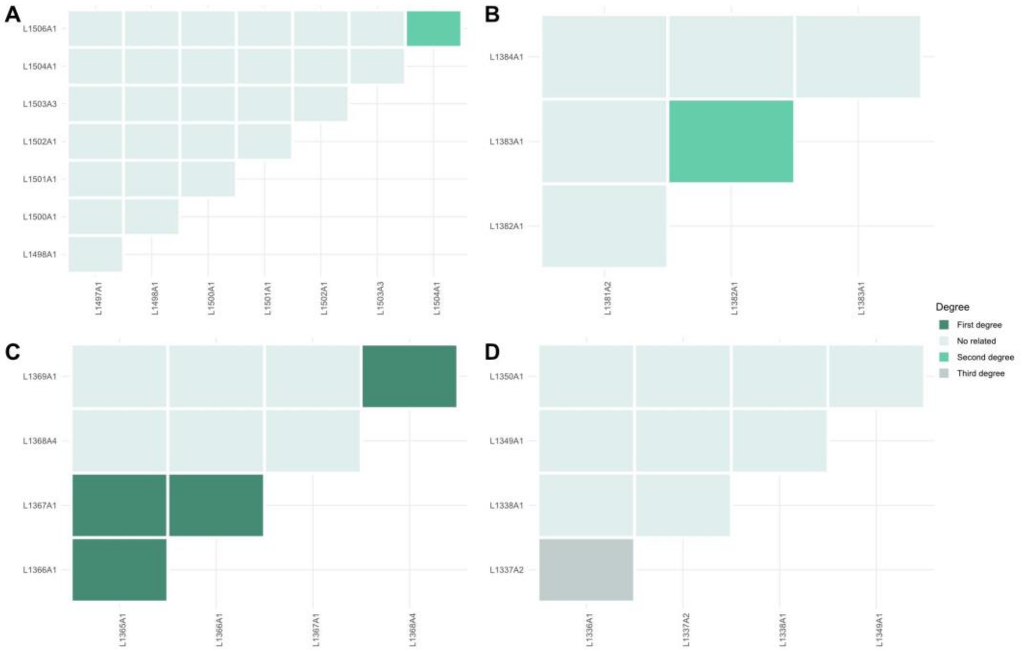


Figure 31 Supplementary. Relatedness pairs of western lowland gorilla sites. A) Bel B) Lob C) Lom D) Mam.

### 2.4 Variant calling metrics

In order to define proper filters for calling genotypes in this dataset, statistics of the called variants were calculated.

Genotype quality was measured for all variants in order to evaluate the overall quality of the variants (Figure 32 Supplementary). Finally, a threshold of a minimum of 20 was applied to filter out variants.

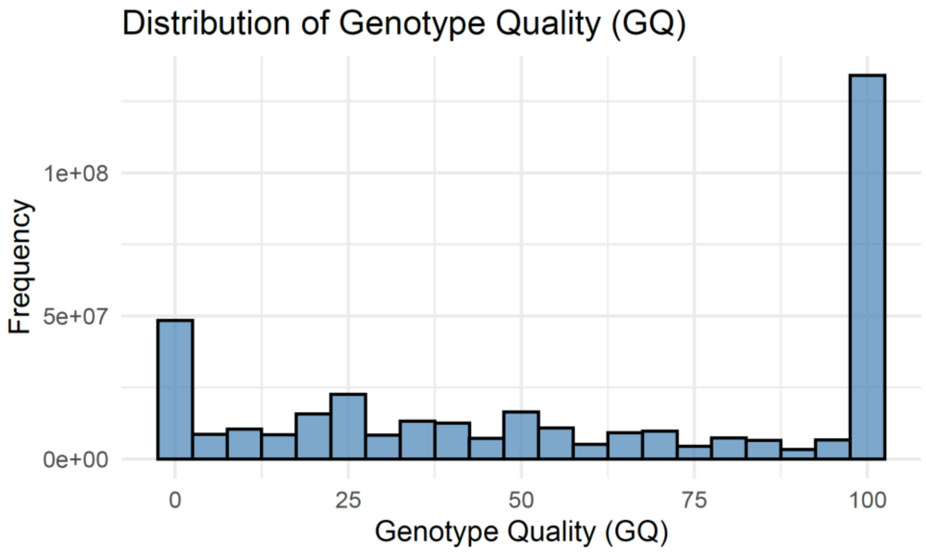


Figure 32 Supplementary. Genotype quality distribution for all called variants (samples > 5X)

Missing data for each sample and each site was calculated from the VCF file using the --missing-indv and --missing-site options of VCFtools (Danecek et al. 2011). Missing data versus coverage per sample was plotted (Figure 33 Supplementary).

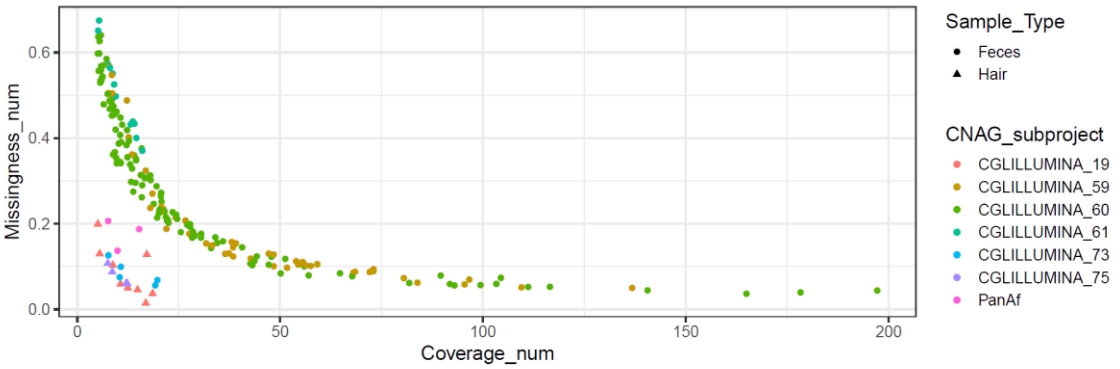


Figure 33 Supplementary. Coverage against missingness for the called variants (samples > 5X), per sample type and sequencing batch (CNAG subproject).

Depth of coverage was extracted for all positions using the --site-mean-depth option of VCFtools (Danecek et al. 2011).

### 3. Population structure

Population structure was assessed using ANGSD (Korneliussen et al. 2014) and pcANGSD (Meisner and Albrechtsen 2018) and NGSadmix (Skotte et al. 2013), which uses as input genotype likelihoods, for both chromosome 21 and exome captures.

#### 3.1. Principal component analysis

We performed principal component analysis (PCA) of chromosome 21 and the partial exome capture data using PCAngsd (Meisner and Albrechtsen 2018) (v1.21).

Chromosome 21:

We performed PCA at different levels: species, subspecies and population level. For this analysis, all samples (< 1% human contamination and >= 0.5X average coverage) were used (faecal, hair, FTA card and museum samples), including chromosome 21 extracted from the publicly available genomes (Figure 34 Supplementary). The Ebo forest sample was included in the following analyses. No sample type bias was seen in any of the principal component analyses performed using chromosome 21 data (Figure 34-43 Supplementary).

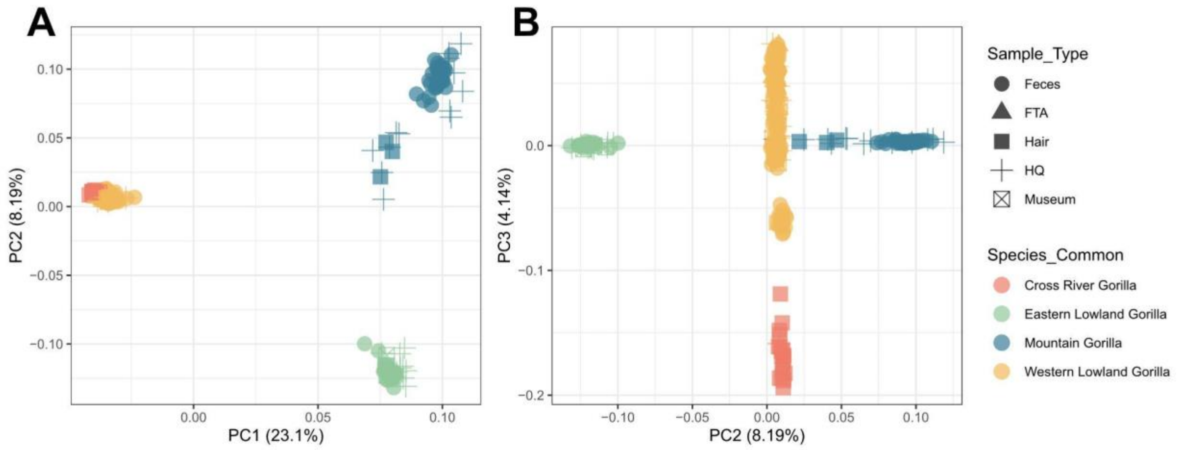


Figure 34 Supplementary. Principal Component Analysis gorilla genus (including all samples which passed the filters and the Ebo sample). 299 samples and 159.260 SNPs were used for this analysis. Each dot represents a unique sample, and each color represents the subspecies of each sample (eastern lowland gorillas - green, mountain gorillas - blue, western lowland gorillas- yellow and Cross River gorillas - red). Shapes represent the different sample types used (faecal samples, FTA cards, hair samples, museum samples, and publicly available genomes for which chromosome 21 was extracted).

Western gorilla species:

The principal component analysis of western gorillas clearly differentiated the two subspecies (highlighted in two different colors - yellow and red), as expected. Inside the western lowland subspecies, three main clusters can be distinguished, one of them clustering closer to the Cross River gorilla subspecies for the first component (Figure 35 Supplementary).

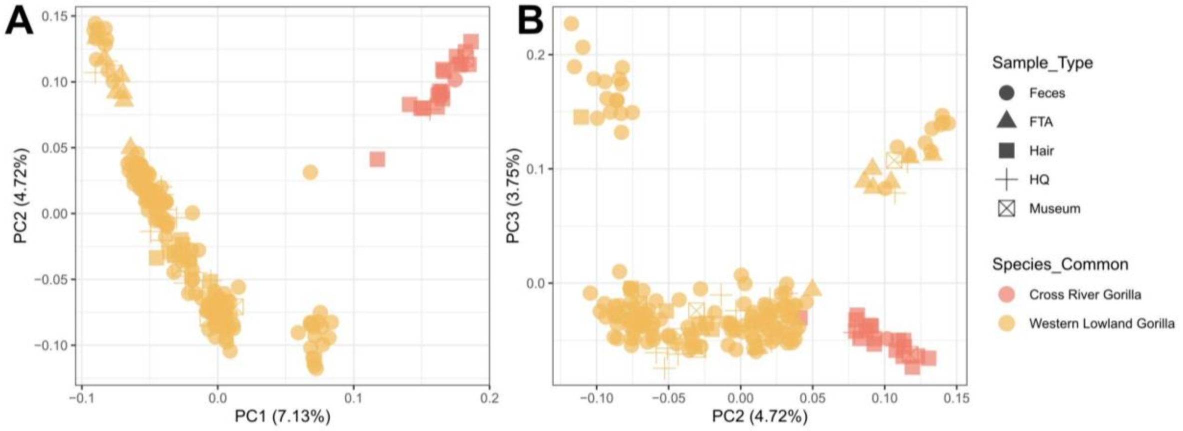


Figure 35 Supplementary. Principal Component Analysis of western gorillas (including Ebo forest sample). 205 samples and 181.947 SNPs were used for this analysis. Each dot represents a unique sample, and each color represents the subspecies of each sample (western lowland gorillas- yellow and Cross River gorillas - red). Shapes represent the different sample types used (faecal samples, FTA cards, hair samples, museum samples, and publicly available genomes for which chromosome 21 was extracted).

When we colour the western gorillas by country of origin, in the first principal component, the Republic of Congo sites (represented in pink color) are clustered separately in the three main groups (Figure 35 Supplementary). The Republic of Congo comprises the eastern part of the western lowland distribution, also being the southwest part of the distribution range. We have different sites from this country (Gou - northern part of the country, Odzala National Park sites (Nga-Dib-Dze and Lok-Imb) and Conkouati (Con) - southern part of the country). Con site clustered with all southern sites from Gabon. Gou site clustered with Dzanga-Ndoki National Park (Dza). The rest of the Republic of Congo sites are from the Odzala National Park (east to Gabon), which clustered together with all northern Gabon sites and Cameroon sites. We identified the L4450A1 sample as an outlier of the Gou sampling site, and therefore was removed from further analyses.

The unique Ebo sample used for this analysis did not cluster with any of the two western subspecies (highlighted with a red arrow) (Figure 36 Supplementary).

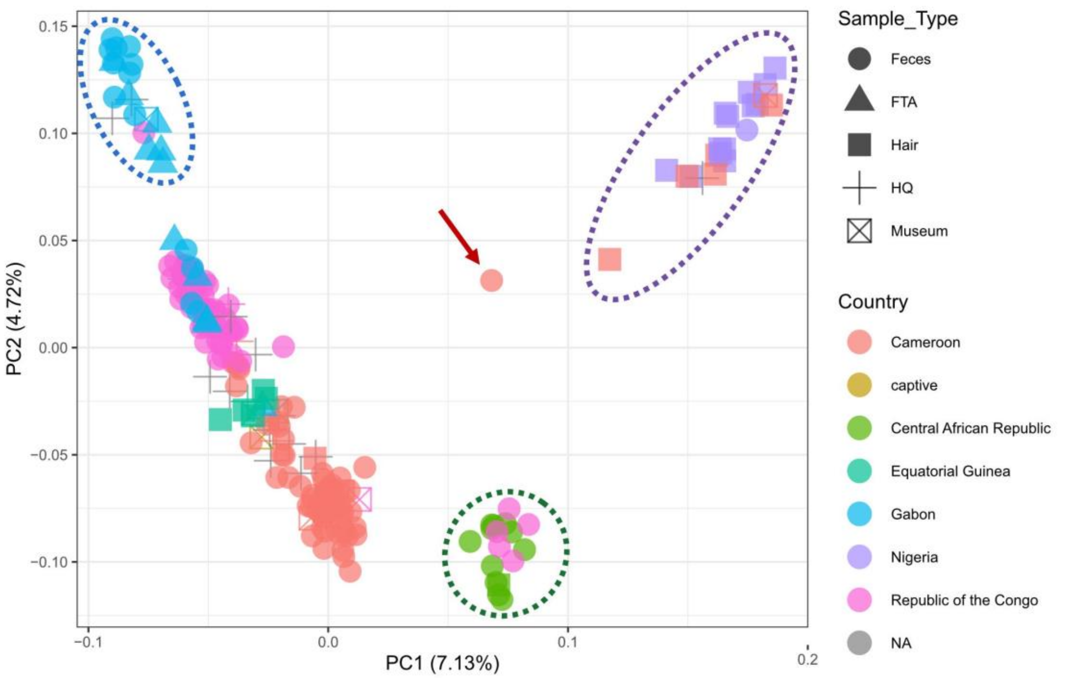


Figure 36 Supplementary. Principal Component Analysis of western gorillas (including the Ebo forest sample). the red arrow indicates the Ebo forest sample. The purple circle indicates the Cross River gorilla cluster. The green circle indicates the Dzanga-Ndoki and Gou cluster. The blue circle highlighted the southern cluster sites. Each dot represents a unique sample, and each color represents the country origin of each sample.

Western lowland gorilla subspecies:

Western lowland gorilla substructure was detected in the principal component analyses of only western lowland gorillas. For this analysis, we also included the unique Ebo sample and the 7 museum and 12 FTA card samples with metadata associated as western lowland gorillas. The three main population groups seen above were also detected independently of the sample type (Figure 37 Supplementary). The first component follows geographic order from north-east to south-west: from Dzanga-Ndoki National Park and the Republic of Congo (Gou) to the south of Gabon and the Republic of Congo (Conkouati), passing through Cameroon, Equatorial Guinea and the rest of Gabon in order. The second component divides the main group of western lowland gorilla sites from Gabon, Cameroon, Equatorial Guinea and part of the Republic of Congo from the other two clusters of sites from south of Gabon and the Republic of Congo (Con) and from Dzanga-Ndoki National Park and the Republic of Congo (Gou).

Previous work by Fünfstück et al. (2014) identified three main clusters within western lowland gorillas, using microsatellites; the clusters they identified were between Lob, Odzala National Park area and Dzanga-Ndoki National Park. Using our denser sampling dataset on chromosome 21 data, the PCA clusters (Supplementary Figure X) separate southern Gabon (not present in Fünfstück et al. (2014)), Dza-Gou (present in Fünfstück et al. (2014)), and the rest of the western lowland gorilla populations (including both Odzala and Lobeke, which were also present in Fünfstück et al. (2014)).

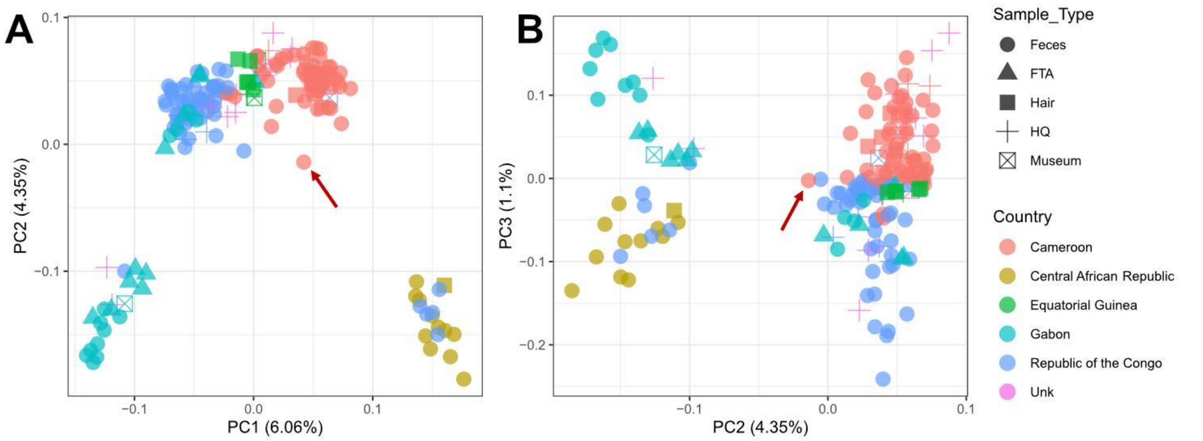


Figure 37 Supplementary. Principal Component Analysis of western lowland gorillas (including the Ebo forest sample - highlighted with a red arrow). Each dot represents a unique sample, and each color represents the country of origin of each sample.

For the FTA and museum samples from the western lowland gorilla subspecies, sample origin or rescued area information was available for the majority of them. The first and second principal components verified the origin information obtained from these samples at the country level, as each sample clustered near samples from the same country (Figure 38 Supplementary).

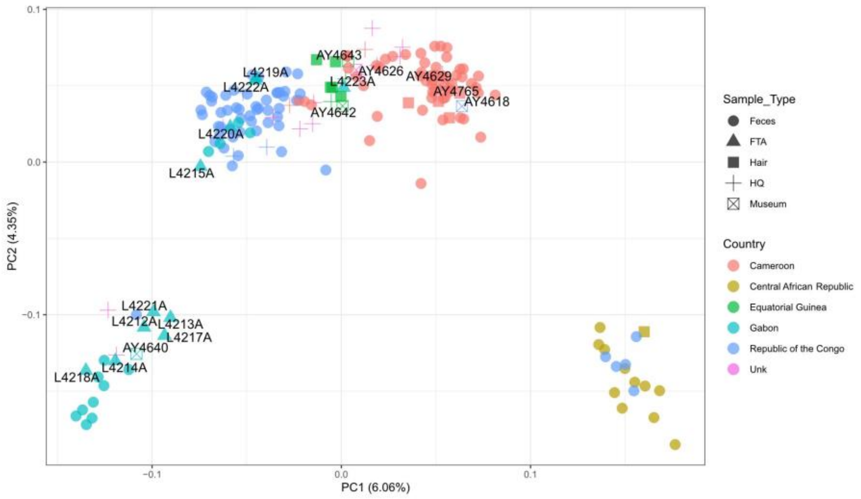


Figure 38 Supplementary. Principal Component Analysis of western lowland gorillas (including Ebo forest sample), including labels for FTA and museum samples. Each dot represents a unique sample and, each color represents the site of the sample.

Each western lowland gorilla cluster detected in the subspecies principal components analysis was deeper analysed independently. Cameroon-Gabon-R.Congo showed a clear geographic distribution, not showing any clear independent cluster (Figure 39A Supplementary). Dza and Gou sites showed differences between the two sampling sites; the first component separates Mongambe from Bai Hokou gorilla social groups within the Dza site (Figure 39B Supplementary). Finally, the samples from the south of Gabon did not show a specific pattern of clustering related to geography; first and second components separated samples from Mou (Figure 39C Supplementary). In this case, the majority of samples were from the same site (Mou) or were FTA cards for which geographic coordinates were not available (just origin information).

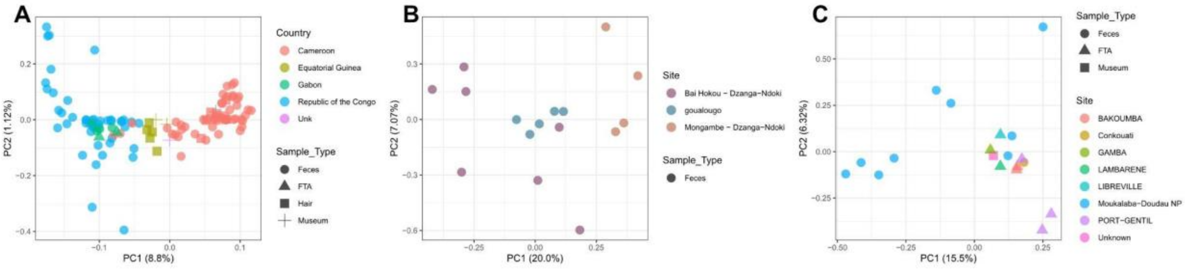


Figure 39 Supplementary. Principal Component Analysis of western lowland gorilla groups ( including FTA and museum samples). Each dot represents a unique sample, and each color represents the site of the sample or the country of origin.

Cross River gorilla subspecies:

Principal component analysis of Cross River gorillas (Figure 40 Supplementary) was performed using chromosome 21 of previously published data (Alvarez-Estape et al. 2023;

Prado-Martinez et al. 2013; Fontsere et al. 2022) and the newly sequenced sample from Ebo forest and one museum sample labelled to be from the Mamfe sampling site, from which genomic data had not been previously sampled (blue cross sign). The population clusters were consistent with those described in Alvarez-Estape et al. 2023, the newly sequenced museum specimen from Mamfe separated from Kag in the first component, and in the second component between gorillas at Afi and those in Bos-Mbe. We recommend further sampling of Cross River gorillas at Mamfe to refine our understanding of the genetics of the gorillas at this sampling site, which is relatively far from the other populations of Cross River gorillas.

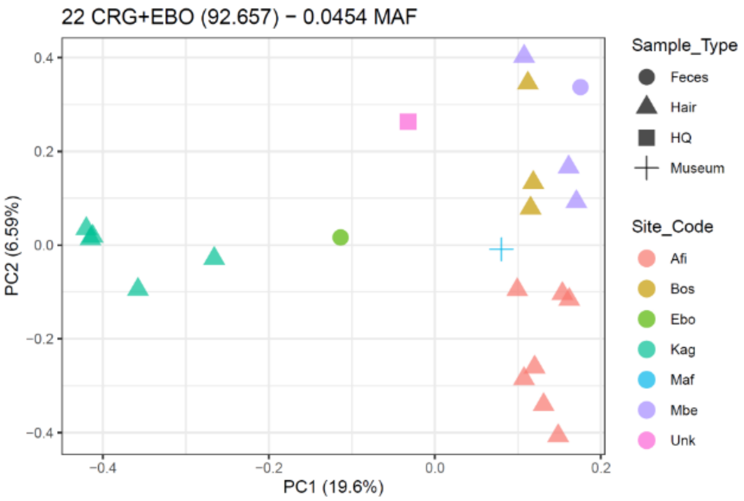


Figure 40 Supplementary. Principal Component Analysis Cross River gorillas (including Ebo forest sample). Each dot represents a unique sample, and each color represents the site of the sample.

Eastern gorilla species:

The principal component analysis of eastern gorillas clearly differentiated the two subspecies (highlighted in two different colors - green and blue), as expected. The second component differentiates the two isolated mountain gorilla populations, Bwindi and Virunga, highlighted in dashed circles. The orange circle for the Bwindi population, and the purple circle for the Virunga population. The third component explains genetic differences among the eastern lowland gorillas (Figure 41 Supplementary).

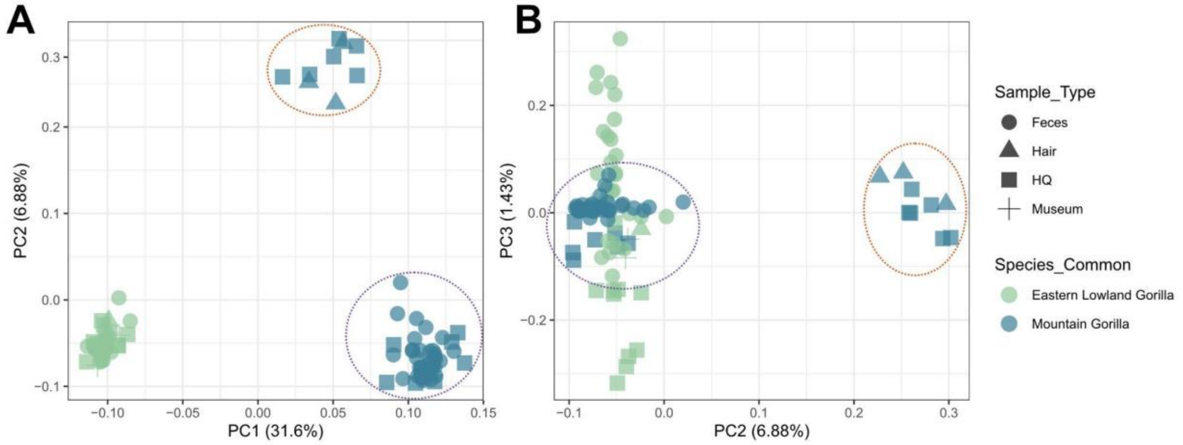


Figure 41 Supplementary. Principal Component Analysis Eastern gorilla species. The purple circle highlights the Virunga population, and the orange circle highlights the Bwindi population.

Eastern lowland gorilla subspecies:

Principal component analysis for eastern lowland gorillas showed geographic clustering in the first component, separating samples from the Kahuzi-Biega National Park from the rest of the sites, except for the publicly available whole genome  from Kahuzi-Biega (Figure 42 Supplementary). Geographic coordinates were available for all faecal samples, but only region of origin information was available for the majority of eastern public genomes. For example, for the Itebero sample (red square), the metadata information indicated origin from Kahuzi-Biega National Park, but specific geographic coordinates are not provided. Kahuzi-Biega National Park has an extension of 6000 km2 (UNEP-WCMC 2025), meaning that this sample could be from a very distinct region of the National Park compared to the rest of the faecal samples from this site, and this is what is suggested from PCA. The second component separates Maiko, Nkuba Conservation Area, Tulakwa, and the public sample from Kahuzi-Biega from the rest of the public genomes from unknown precedence, Walikale area and Mount Tshiaberimu (Figure 42 Supplementary). Three samples from Walikale have been added to this PCA, showing different patterns of clustering in the second component. More samples from different regions will be needed to clarify the possible population structure inside the eastern lowland gorillas, which have the second largest distribution range of the gorilla subspecies (Plumptre et al. 2016).

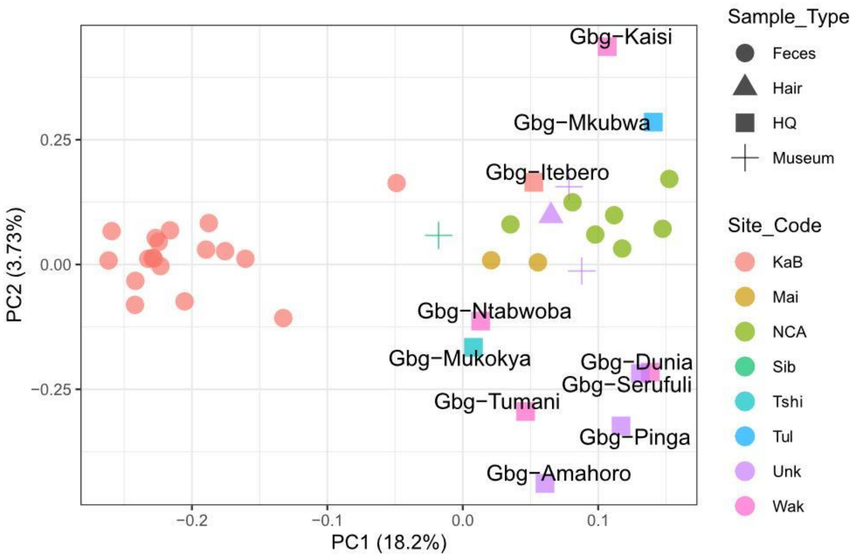


Figure 42 Supplementary. Principal Component Analysis eastern lowland gorilla species, including publicly available genomes (sample IDs indicated in the plot).

Mountain gorilla subspecies:

Mountain gorillas were divided into the two well-known isolated populations in the first component of the PCA, Virunga and Bwindi, as expected. The second component showed differences inside the Bwindi and Virunga populations (Figure 43 Supplementary).

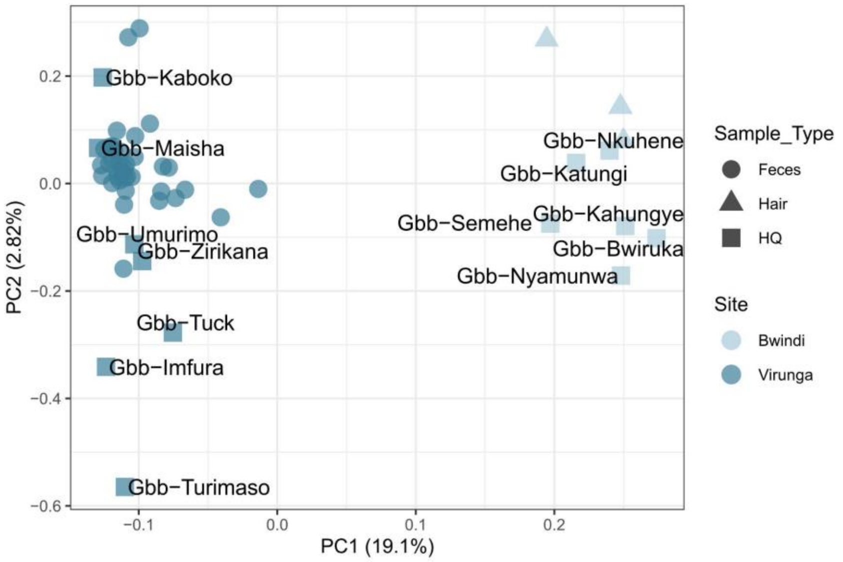


Figure 43 Supplementary. Principal Component Analysis mountain gorilla species, including publicly available genomes (sample IDs indicated in the plot).

Partial exome:

The partial exome was also used to study possible population structure within the gorilla genus following the same approach as used for chromosome 21, requiring 25% of individuals to be covered per nucleotide site. We also performed PCA for each subspecies

We also performed PCA using post QC filtering by < 1% human contamination, >= 0.5X average coverage and no self or first-degree relatives (237 samples, 232 faecal samples, 5 hair samples). In PCA for the western lowland gorilla subspecies, we also identified one sample, L4450A1, which falls outside its population cluster (for the Gou sampling site) and was removed from the downstream analyses.

In PCA of the partial exome on-target space, the newly sequenced gorilla samples clearly separate by subspecies, as expected (Figure 44 Supplementary) and in accordance with chromosome 21 data.

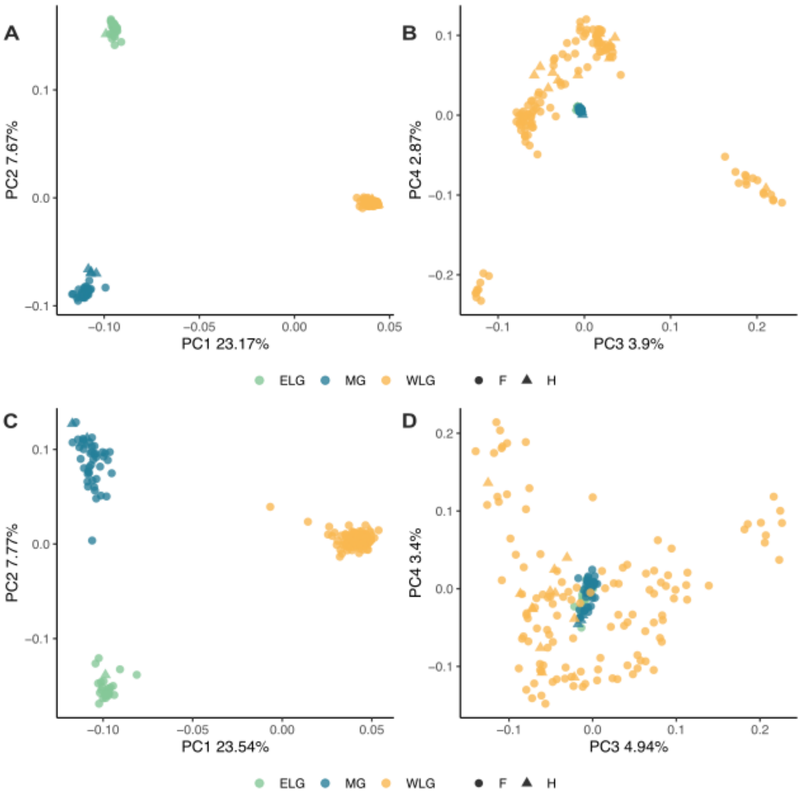


Figure 44 Supplementary. Comparison of the partial exome and chromosome X for the newly sequenced non-invasive gorilla samples. PCA of the partial exome (A and B) and the exome of chromosome X (C and D) of retained post-QC samples (< 1% human contamination, >=0.5X average coverage, no self or first-degree relatives) n=204.

PCA within each subspecies indicates subpopulation clusters within Cross River gorillas and mountain gorillas (Figure 45A and B-46C and D Supplementary), in agreement with previous literature (Xue et al. 2015; Alvarez-Estape et al. 2023) and observations in the georeferenced chromosome 21 data for the same gorillas. No clear genetic structure is observed within eastern lowland gorillas (Figure 46A and B Supplementary).

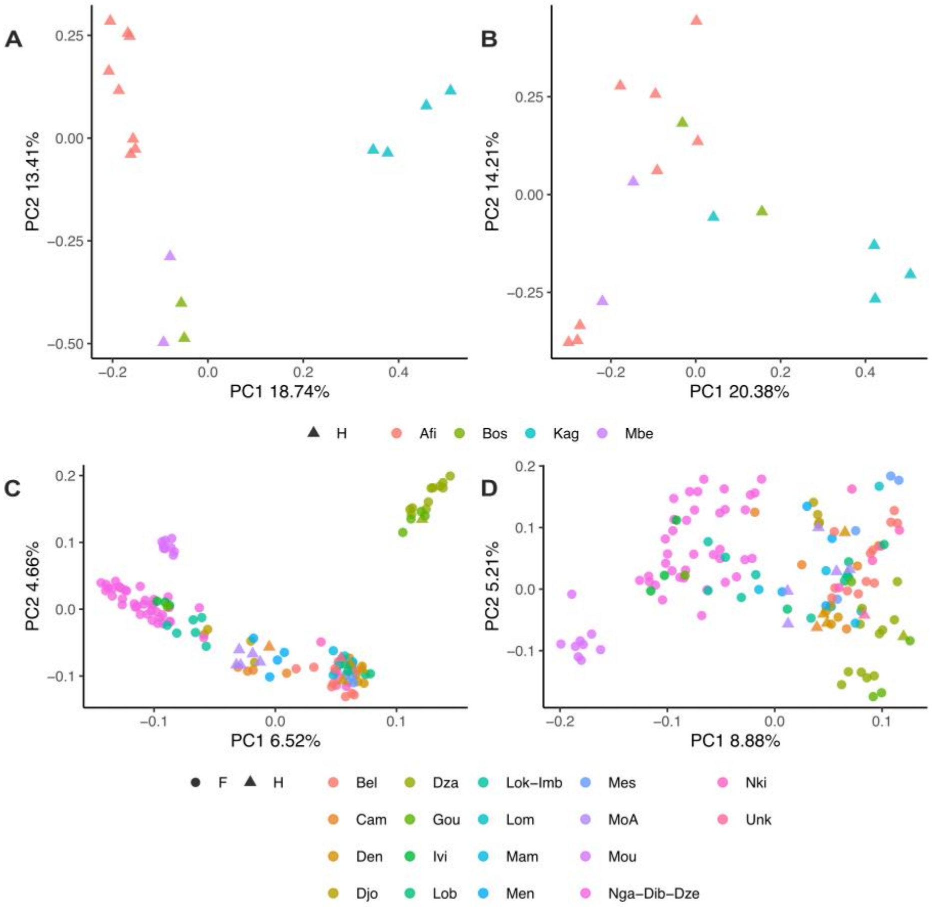


Figure 45 Supplementary. Comparison of the partial exome and chromosome X for the western gorilla species. A) Partial exome of CRG. B) Chromosome X exome of CRG. C) Partial exome of WLG. D) Chromosome X exome of WLG.

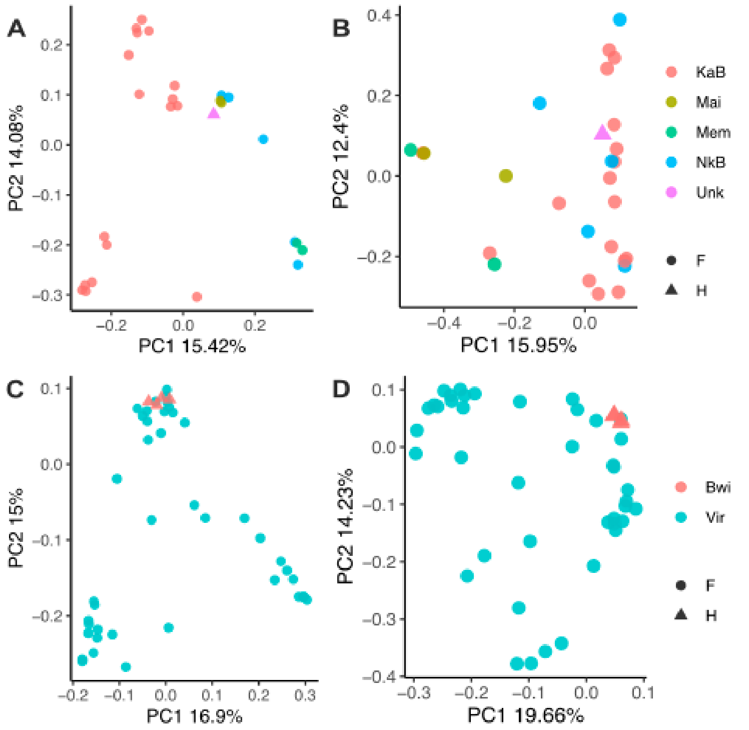

Figure 46 Supplementary. Comparison of the partial exome and chromosome X for the eastern gorilla species. A) Partial exome of ELG. B) Chromosome X exome of ELG. C) Partial exome of MG. D) Chromosome X exome of MG.

#### 3.2. Admixture

For both chromosome 21 and exome data, clear genetic differentiation between the two gorilla species is observed in PCA (Figures 33 and 43 Supplementary) and structure analysis (Figures 47 and 55 Supplementary), where K=2 is the best inferred K. This agrees with previous findings of a deep divergence between the eastern and western gorilla species (van der Valk et al. 2024; Thalmann et al. 2007).

Chromosome 21:

Using chromosome 21 data, at K=4 the Cross River gorillas component appeared. Some western lowland populations showed part of this component, distinguishing three different scenarios: not having Cross River component, having some part of Cross River component and having ~40% of Cross River component. At K=5, we observe a degree of allele sharing between eastern lowland gorillas and mountain gorillas from Bwindi. In addition, at K=5, another western lowland gorilla component appeared and was spread to almost all western lowland populations, except for the samples from south of Gabon. At K=6, Virunga and Bwindi were differentiated, showing different ancestry components. At K=7, a unique ancestry component separated samples from Dza and Gou from the rest of the western lowland gorillas.

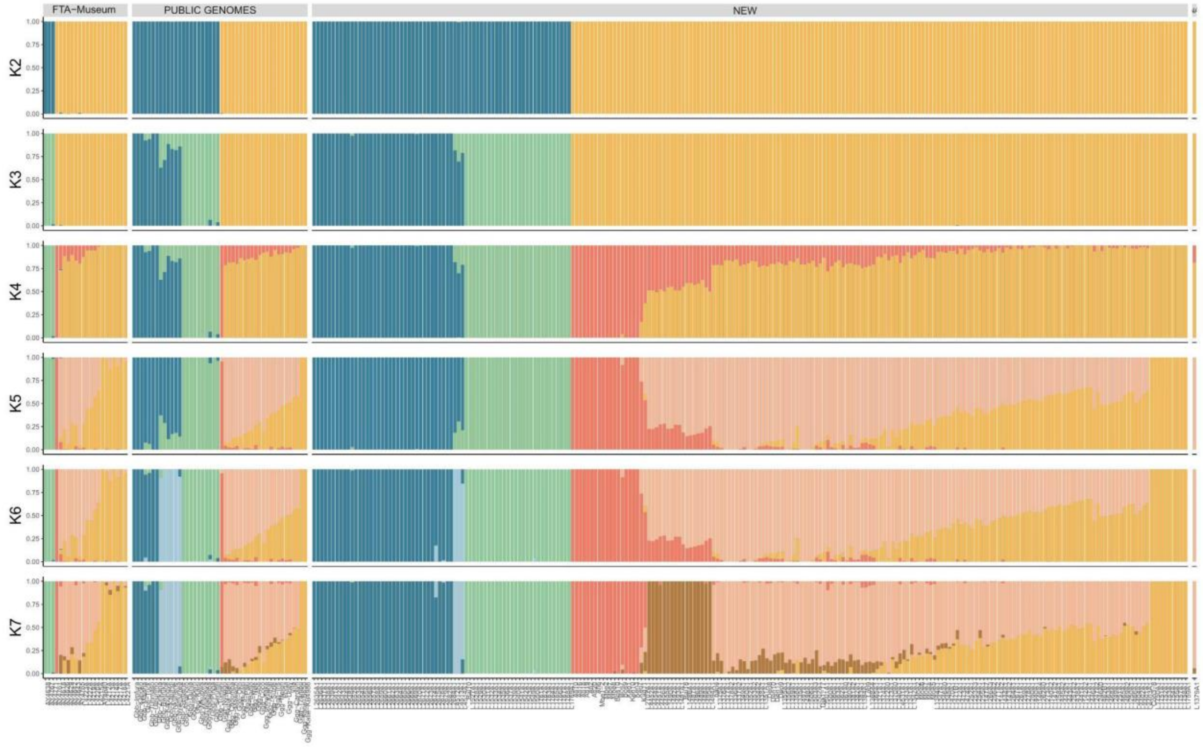


Figure 47 Supplementary. ADMIXTURE PLOT ALL DATASET

Plotting K=7 for each sample in the geographic coordinates provided showed the clear geographic structure also seen in the different principal component analyses (Figure 48 Supplementary). The unique sample from Ebo forest showed admixed components from Cross River gorillas and western lowland gorillas, in concordance with western gorilla principal component analyses (Figure 34-35 Supplementary). Samples from the central and eastern part of Cameroon showed more allele sharing with Dza and Gou than the rest of the western lowland populations (Figure 48 Supplementary).

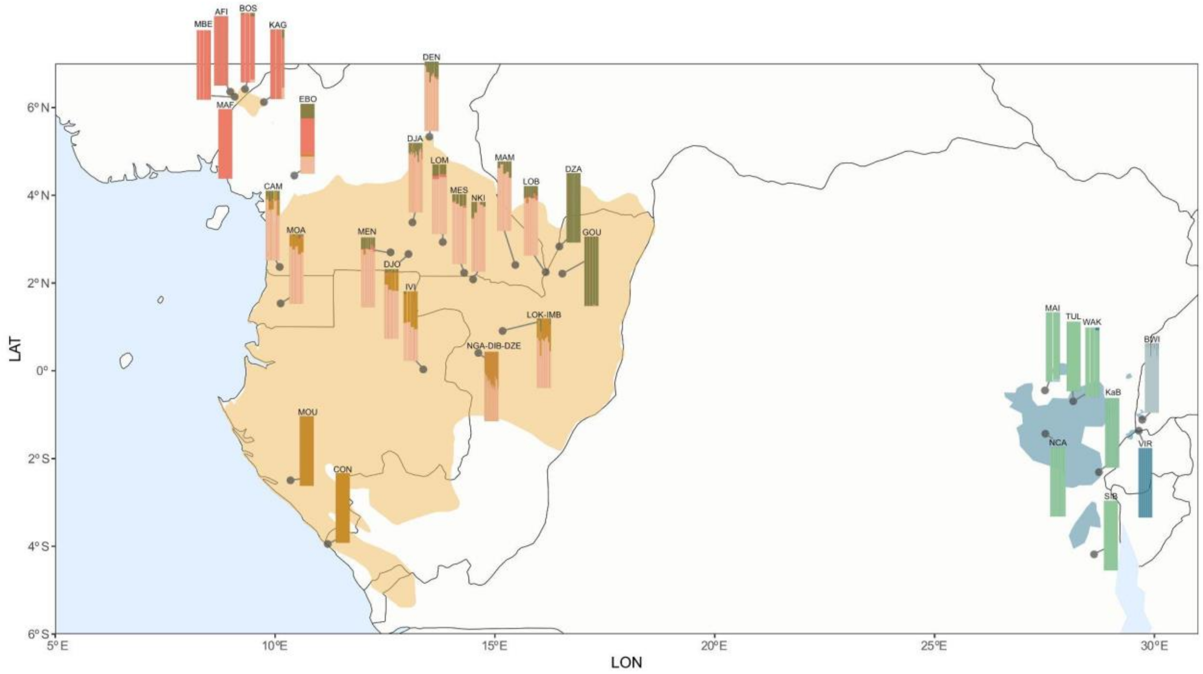


Figure 48 Supplementary. ADMIXTURE PLOT ALL DATASET showing k = 7. Each bar is plotted in the geographic coordinates provided.

Eastern gorillas admixture analysis showed a clear separation between eastern lowland and mountain gorillas at K=2 (Figure 49 Supplementary). It was also possible to see some allele sharing between the Bwindi population and eastern lowland gorillas, when compared to Virunga population. At K=3 Virunga and Bwindi were distinguished, seeing the same patterns of admixture components for the three types of data (faecal with geographic coordinates, public genomes, and museum samples, both with origin information). At K=4 and K=5 different patterns in the faecal samples were observed.

A distinct cluster inside the eastern lowland gorillas admxiture analysis at K=2 appeared showing differences between samples fromMaiko and Nkuba Conservation Area and samples from Kahuzi-Biega National Park (Figure 50 Supplementary), as seen in the first component of the eastern lowland principal component analysis (Figure 41 Supplementary). We include the hair sample Bai10 (Alvarez-Estape et al. 2023) in this analysis, and confirm their findings that this sample is an eastern lowland gorilla and we find that it has similar ancestry to Maiko and Nkuba Conservation Area samples. In the following Ks (K=3-4) for the eastern lowland gorillas admixture analysis, we can not distinguish a clear pattern of admixture separation (Figure 50 Supplementary). More samples will be needed to better assess possible population structure inside the eastern lowland subspecies.

Mountain gorillas showed no clear structure after Bwindi and Virunga separation at K=2 (Figure 51 Supplementary). Inconsistent clustering appeared in K=3 and K=4 inside Virunga and Bwindi populations, also seen in the eastern admixture at K=5 (Figure 49 Supplementary).

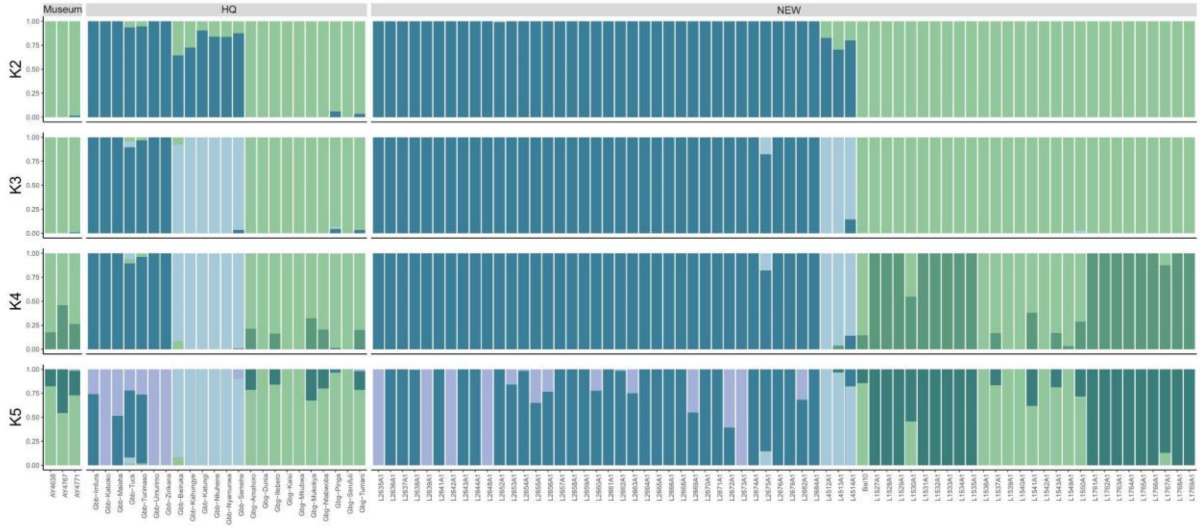


Figure 49 Supplementary. ADMIXTURE PLOT EASTERN GORILLAS

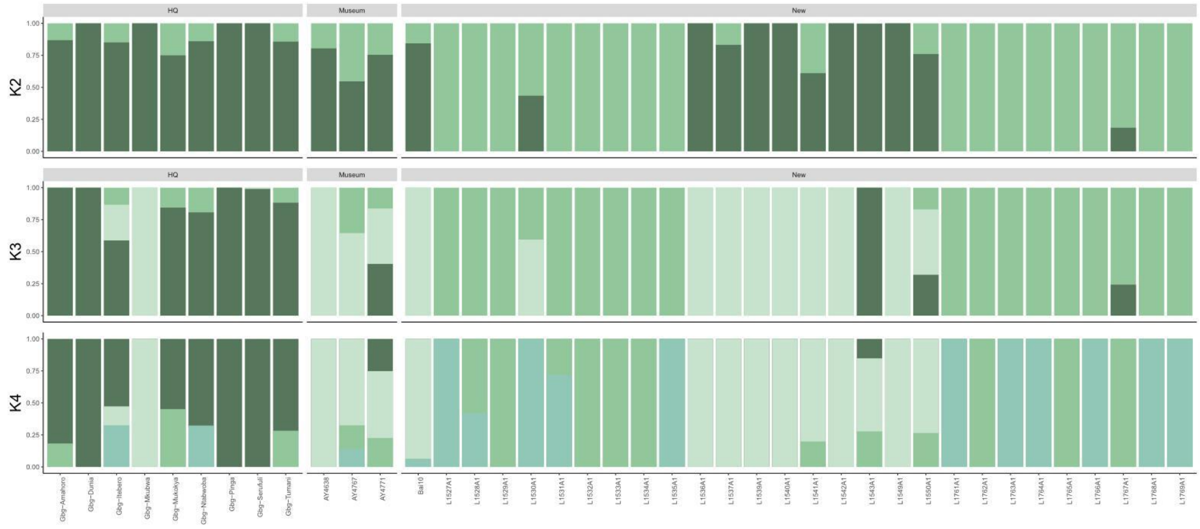


Figure 50 Supplementary. ADMIXTURE PLOT EASTERN LOWLAND GORILLAS

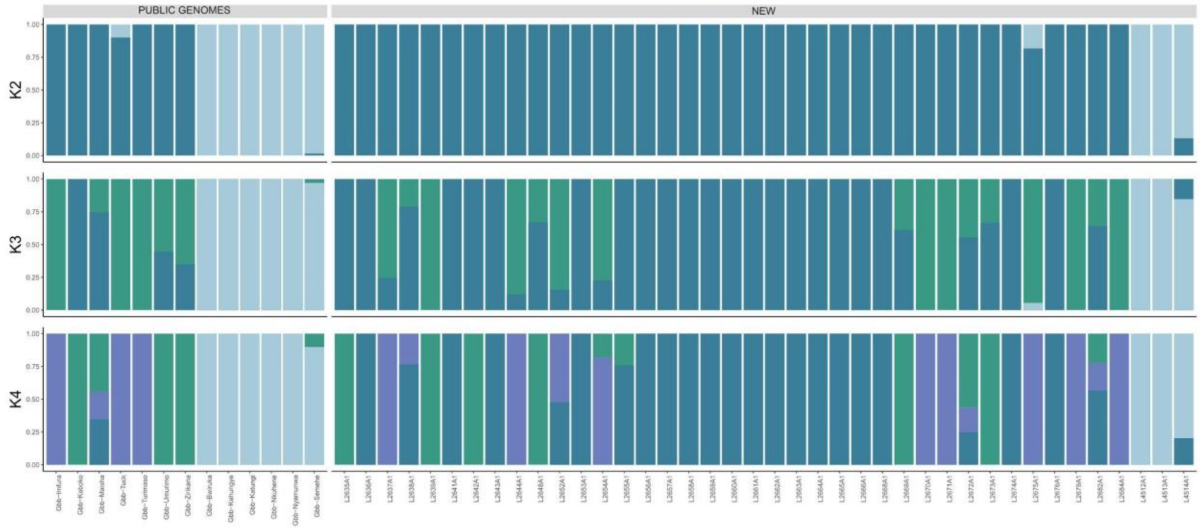


Figure 51 Supplementary. ADMIXTURE PLOT MOUNTAIN GORILLAS

Western admixture analysis showed similar patterns seen in the general admixture using all genus individuals (eastern and western gorillas) (Figure 52 Supplementary). Allele sharing with Cross River gorillas and Dza and Gou populations appeared in K=3. When just analysing western lowland gorillas the similar patterns are observed between the subspecies populations (Figure 53 Supplementary). Cross River gorilla admixture analysis using chromosome 21 data (Figure 54 Supplementary) is consistent with previous results using whole genomes of the same individuals (Alvarez-Estape et al. 2023) and clusters seen in the principal component analysis (Figure 39 Supplementary).

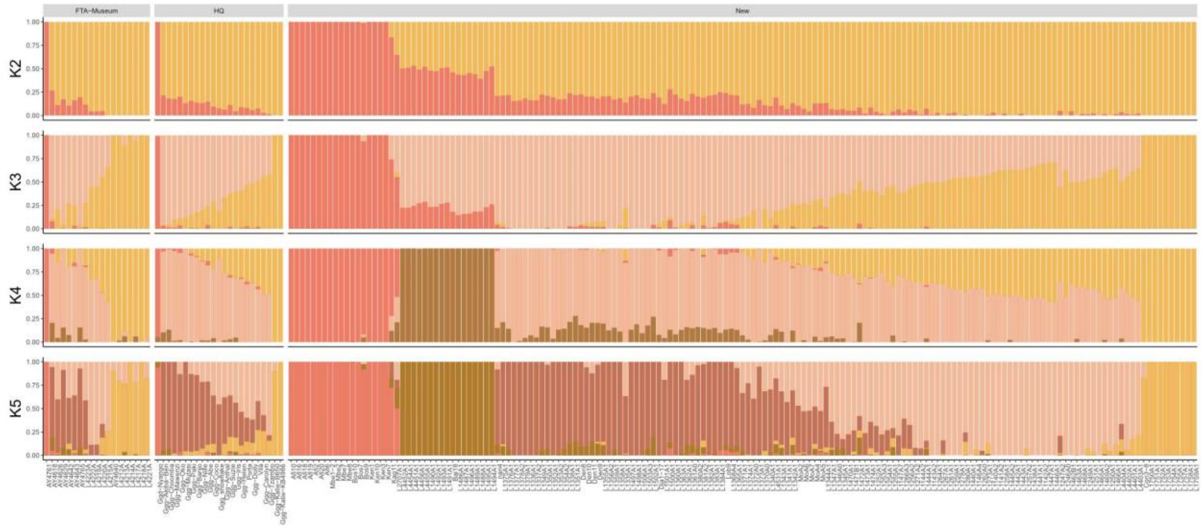


Figure 52 Supplementary. ADMIXTURE PLOT WESTERN GORILLAS

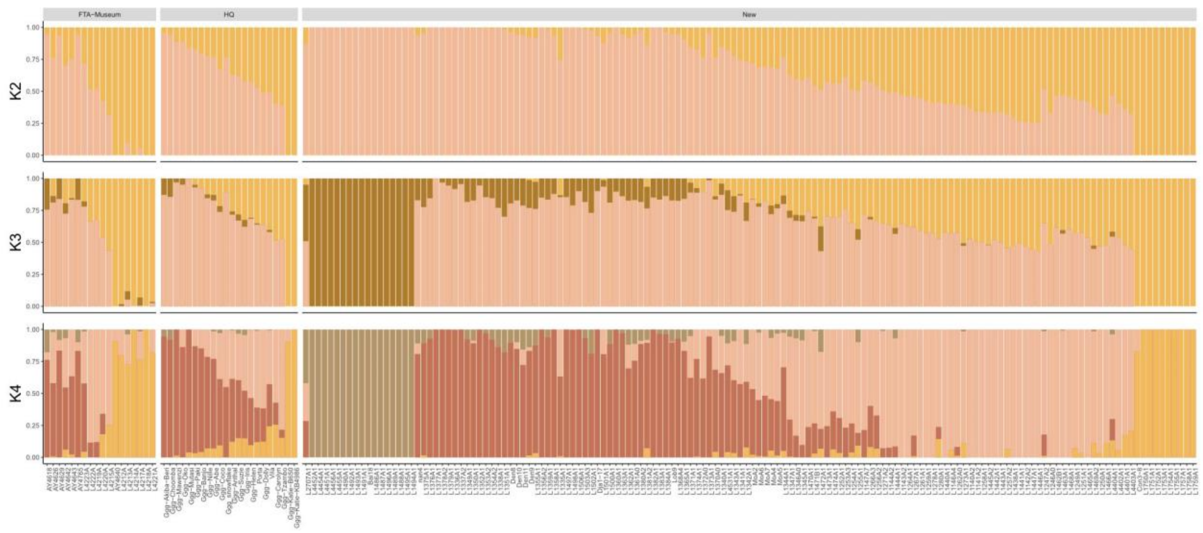


Figure 53 Supplementary. ADMIXTURE PLOT WESTERN LOWLAND GORILLAS

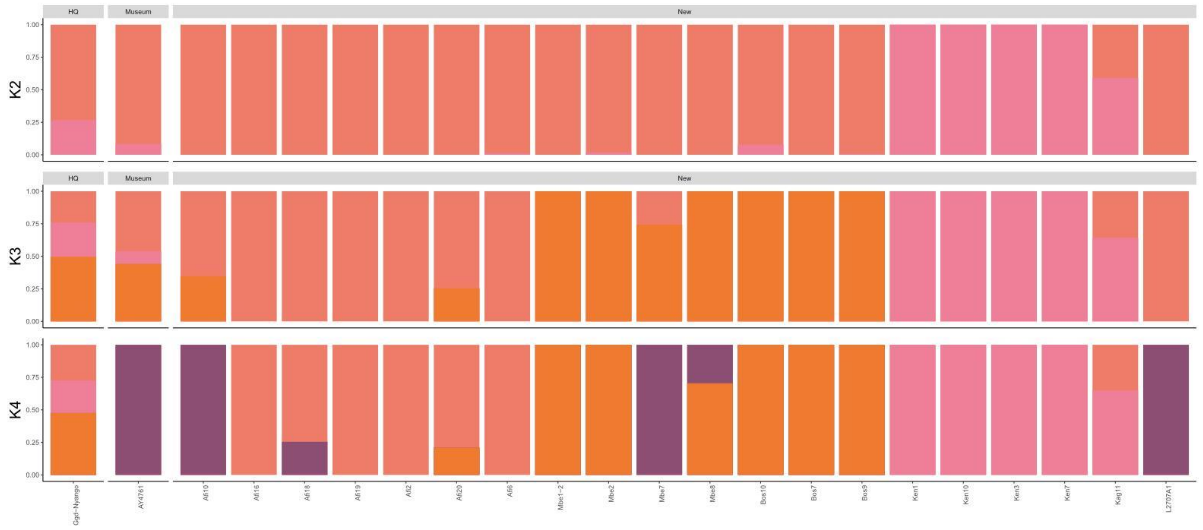


Figure 54 Supplementary. ADMIXTURE PLOT CROSS RIVER GORILLAS

#### 3.3. Heterozygosity

Chromosome 21:

Genetic diversity was assessed by calculating heterozygosity values. In order to include the maximum number of gorilla populations, genotype likelihoods were used for all available samples. All samples with more than 1.5 X of coverage were downsampled to avoid differences due to coverage.

Capture bias was observed when comparing different sample types (Figure 56 Supplementary). FTA cards and hair samples showed an increased heterozygosity value when compared to faecal samples inside the western lowland subspecies. For the rest of the subspecies, it was not possible to compare as different sample types were not available. FTA cards and hair samples were removed from further heterozygosity analysis to avoid capture bias and compare between subspecies and populations. As a consequence, no Cross River gorillas samples remained in genetic diversity analysis.

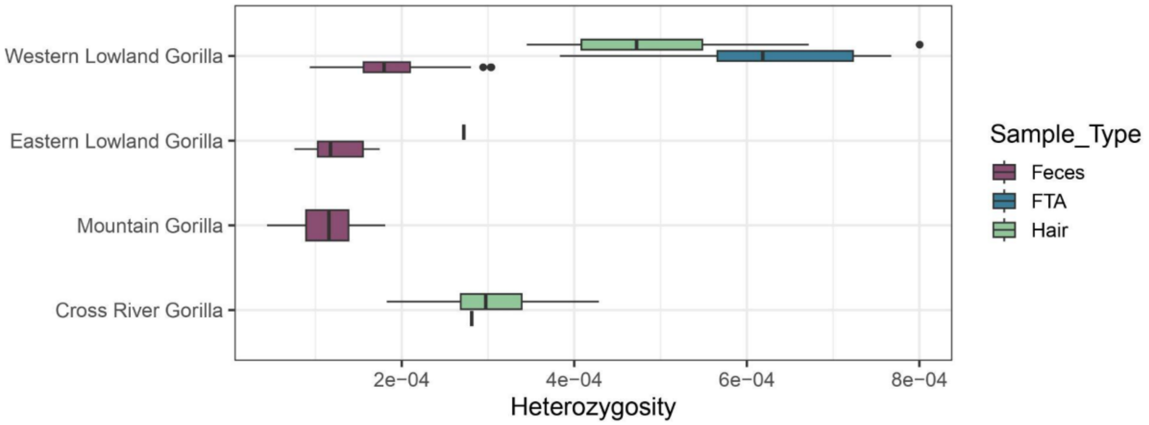


Figure 55 Supplementary. Heterozygosity samples at 1.5X (downsampled) by subspecies and colored by sample type.

After removing different sample types, captured faecal samples chromosome 21 data showed similar patterns of genetic diversity seen in previous studies between subspecies (Alvarez-Estape et al. 2023; Prado-Martinez et al. 2013; Xue et al. 2015). Western lowland gorillas showed an increased value of heterozygosity when compared to the two eastern gorilla subspecies (Figure 56 Supplementary). The difference between species was lower than observed previously due to allelic dropout caused by capture methodology.

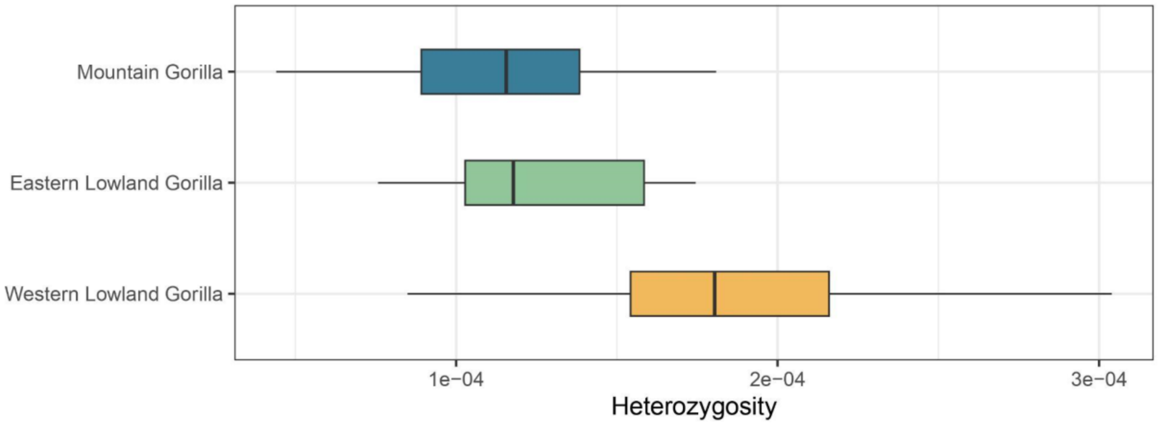


Figure 56 Supplementary. Heterozygosity samples at 1.5X (downsampled) by subspecies. Gorilla subspecies are represented by colors (western lowland gorillas-yellow, mountain gorillas-blue and eastern lowland gorillas-green).

Heterozygosity values within western lowland gorillas by sampling site showed differences between them. These differences did not correspond to PCA or admixture clusters detected, represented in the colors of each box-plot (Figure 57 Supplementary). Ivi and Gou are the sites having higher values of heterozygosity. In contrast, Lob-Imb and Men showed the lowest values.

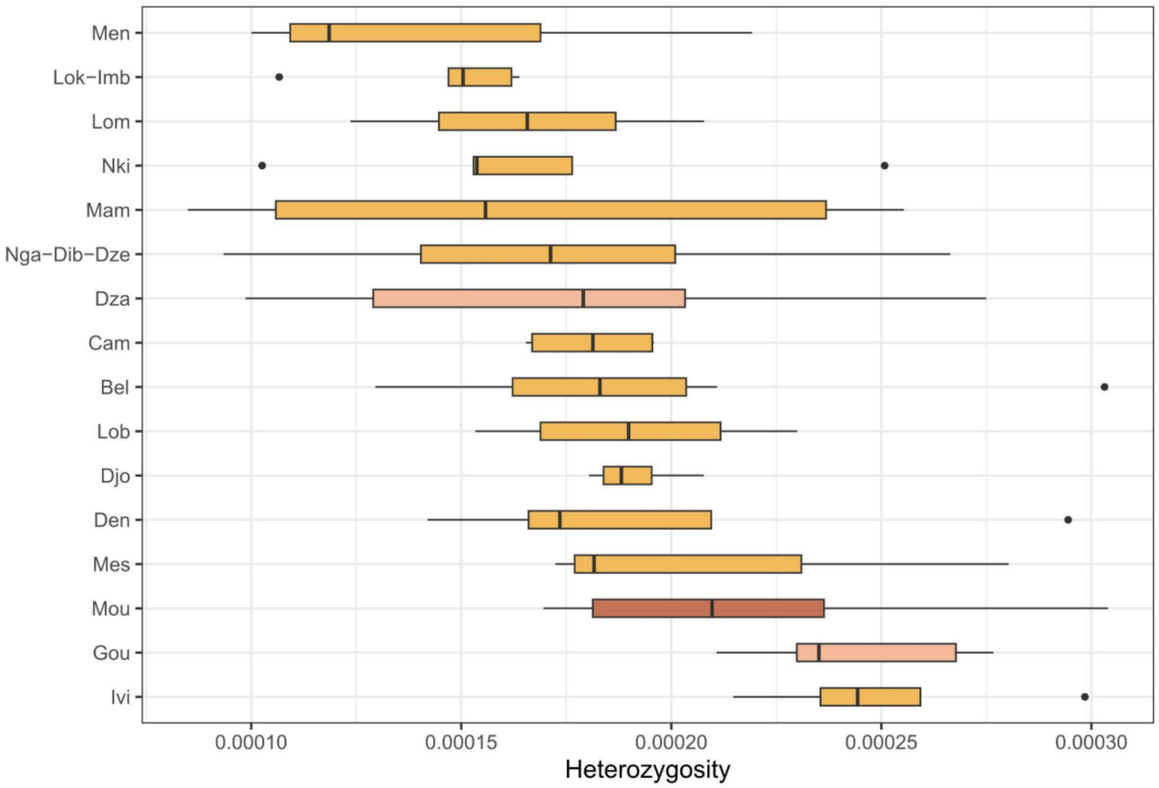


Figure 57 Supplementary. Heterozygosity samples of western lowland gorillas at 1.5X (downsampled) by sampling site. Box-plot colors represent the three main PCA clusters: Dza and Gou in pink, south of Gabon in dark red and the rest in yellow.

Heterozygosity values between eastern gorilla sites showed similar mean values between them (Figure 58 Supplementary). Eastern lowland sites (Maiko, Nkuba Conservation Area and Kahuzi-Biega) had similar low mean heterozygosity values. In contrast to other sites, Kahuzi-Biega samples showed high variability between samples, containing samples with higher values of heterozygosity compared to the rest of the eastern sites. More samples from these same sites and new ones will be needed to better capture the current genetic diversity inside the eastern lowland subspecies. Regarding mountain gorillas, the Virunga population showed lower values of heterozygosity compared to the rest of eastern lowland populations, as expected from results in van der Valk et al. (2024).

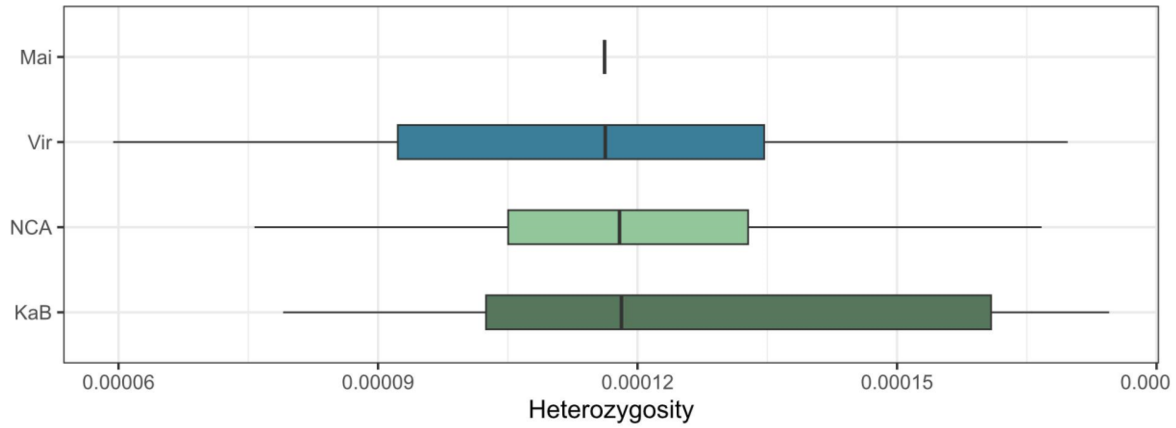


Figure 58 Supplementary. Heterozygosity of eastern gorilla samples at 1.5X (downsampled) by sampling site. Blue colors show mountain gorilla sites, and green colors show eastern lowland sites.

Information about social groups was available for eastern lowland and Virunga mountain gorillas. Comparisons between the different sampled social groups were performed in order to assess the genetic diversity variability inside each population (Figures 59-60 Supplementary). In the case of eastern lowland gorillas (Figure 59 Supplementary), the Kahuzi-Biega site was represented by 7 different social groups: Chimanuka, Ganywamulme, Langa, Mankoto, Mufanzala2, Mugaruka and Namadiriri, and NCA was represented by two different social groups: Membe and Bansamba. Maiko is represented by the same sampling site name. The social groups Membe, Maiko and Bansamba had values within a similar range. The different social groups inside Kahuzi-Biega National Park showed different patterns of heterozygosity between them. Ganywamulme, Langa and Mankoto showed the highest values of heterozygosity. In contrast, Mufanzala2 and Mugaruka showed the lowest values. These differences between social groups could be explained by drift experienced in small populations, inbreeding, gene flow between some social groups, among others. In addition, inbreeding could be affecting the chromosome 21 region; genome-wide heterozygosity and inbreeding levels for each social group will need to be assessed.

Mountain gorillas from Virunga are represented by 21 social groups (More info in Supplementary Table 3). As described for social groups inside Kahuzi-Biega National Park, there were clear differences in heterozygosity among social groups in Virunga (Figure 60 Supplementary).

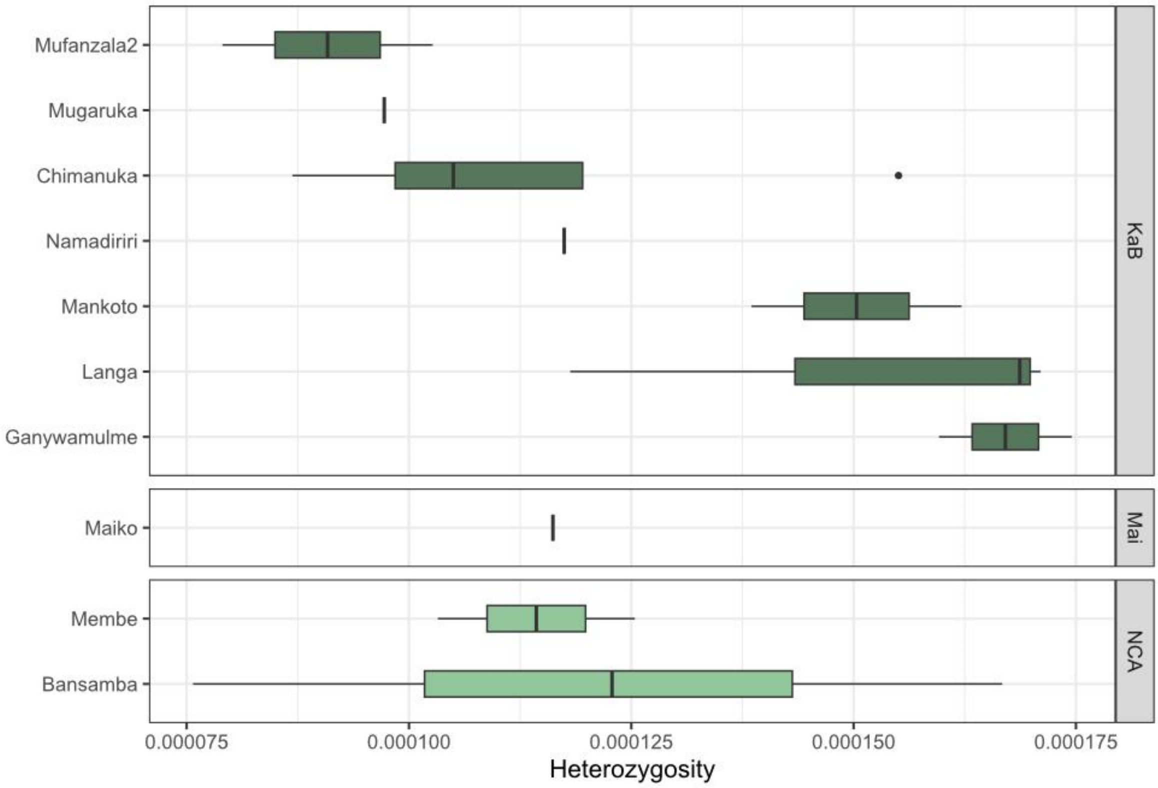


Figure 59 Supplementary. Heterozygosity samples of eastern lowland gorillas at 1.5X (downsampled) by social groups. Different social groups are colored by different green colors (social groups from KaB National Park are colored in dark green, Nkuba Conservation Area (NCA) social groups in blueish-green and Membe social groups in light green).

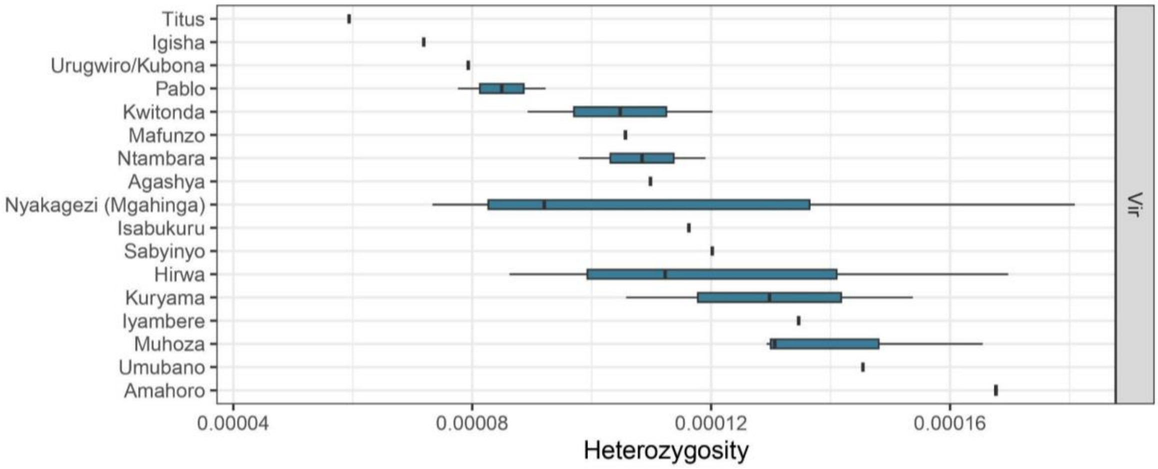


Figure 60 Supplementary. Heterozygosity samples of mountain gorillas at 1.5X (downsampled) by social groups. All Virunga social groups are colored in dark blue.

In addition, we calculated the heterozygosity in higher coverage samples, as the ability to detect heterozygote positions depends on the coverage of the sample. Samples were downsampled to 6, 10 and 20 X of coverage in order to obtain comparable heterozygosity values. For each downsampled group of samples, variants were called using GATK software as described in the methods section.

First, we checked how coverage influences the ability to detect heterozygous positions (Figure 61 Supplementary). As coverage increased, heterozygosity values between the two eastern gorilla subspecies became more distinguishable. Eastern lowland gorillas had higher values of heterozygosity than mountain gorillas, as expected and described in previous studies (Prado-Martinez et al. 2013a; Xue et al. 2015; van der Valk et al. 2024). Using genotype likelihoods or low coverage data (around 6X), these differences were not detected (Figures 56 and 61 Supplementary). Western lowland gorillas showed higher heterozygosity levels than the rest of the subspecies, as it has been previously described (Prado-Martinez et al. 2013a; Xue et al. 2015). This difference, as seen using genotype likelihoods, was not as high as expected, mainly due to possible allelic dropout when using capture data (Figure 61 Supplementary). Cross River gorillas were not added in this comparison as all genomes from this subspecies were generated from hair samples, extracting a posteriori the chromosome 21. This technical difference can produce a bias when detecting heterozygous positions in comparison with capture data, as seen when using genotype likelihoods (Figure 55 Supplementary).

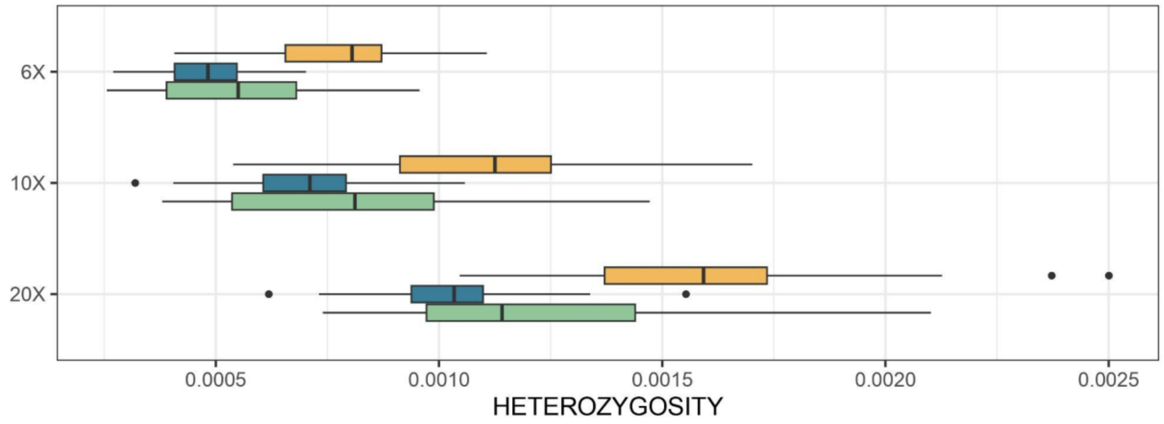


Figure 61 Supplementary. Heterozygosity values at different downsampled coverages (6-10-20X) by subspecies using faecal samples. Each subspecies is represented by different colors (eastern lowland gorillas in green, western lowland gorillas in yellow and mountain gorillas in blue).

We focused on high-coverage samples downsampled to 20X in order to evaluate the maximum detected heterozygous positions (Figure 62 Supplementary). The patterns of heterozygosity coincide with the results commented previously using genotype likelihoods and lower levels of coverage.

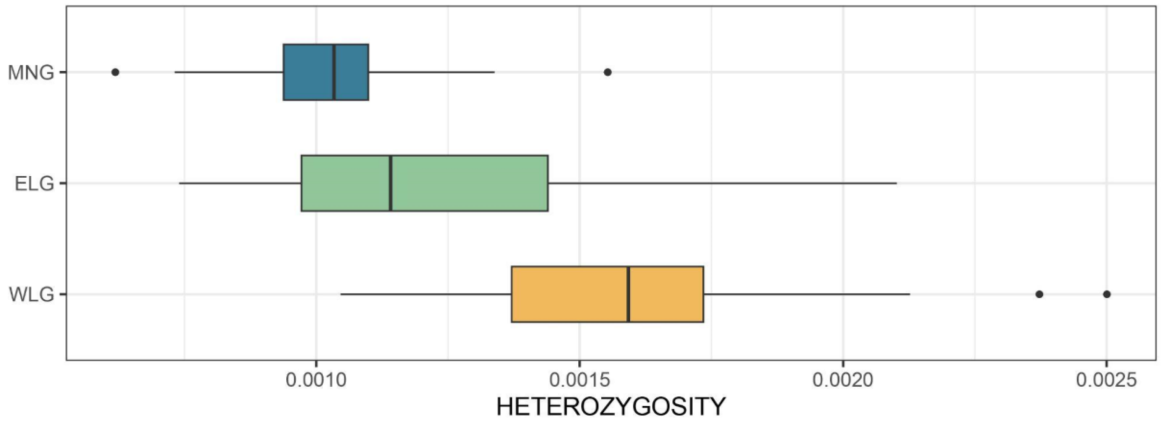


Figure 62 Supplementary. Heterozygosity samples at 20X (downsampled) by subspecies using faecal samples. Each subspecies is represented by different colors (eastern lowland gorillas in green, western lowland gorillas in yellow and mountain gorillas in blue).

Fewer sampling sites are represented when using samples at 20X coverage. The patterns observed using genotype likelihoods were maintained when calculating heterozygosity at 20X for western lowland gorillas. Differences between principal PCA clusters inside western lowland gorillas were not seen either using high coverage data (Figure 63 Supplementary). In contrast, eastern gorilla subspecies heterozygosity values were more differentiated (Figure 64 Supplementary). Eastern lowland gorillas showed higher values compared to mountain gorillas, except for the Maiko eastern lowland site, which showed values similar to Virunga mountain gorilla populations (Figure 65 Supplementary).

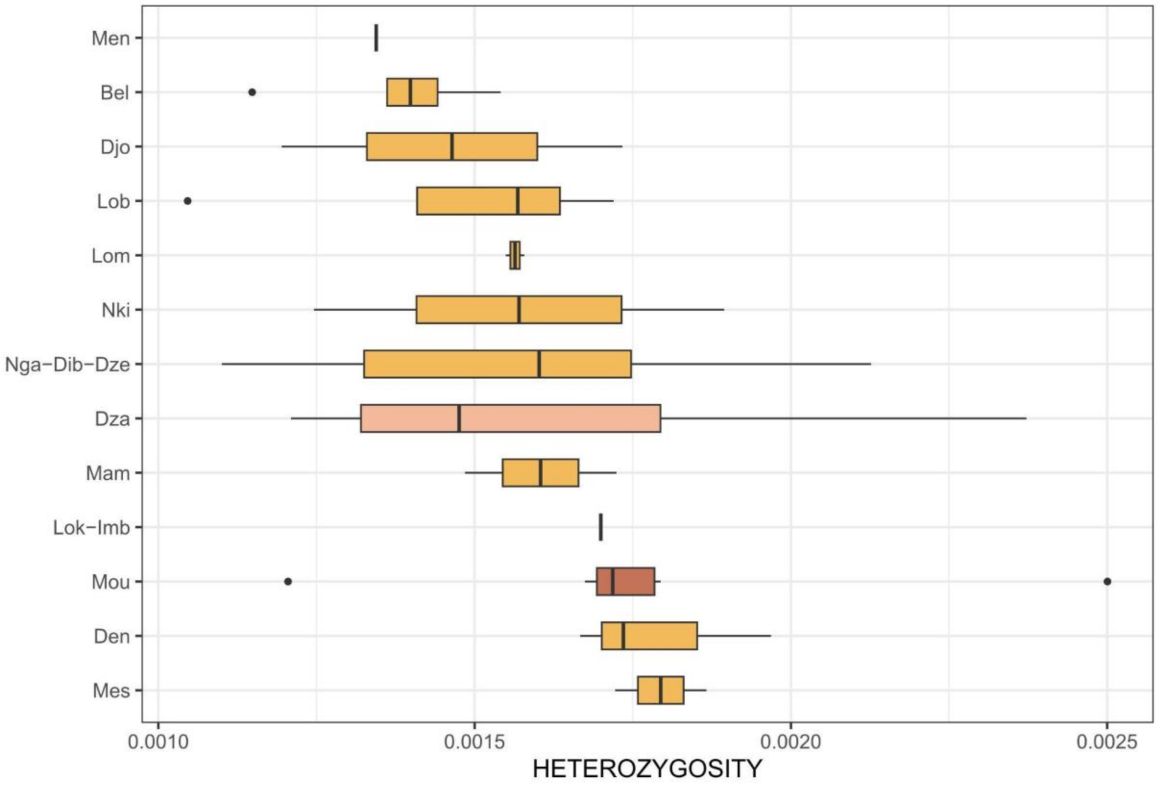


Figure 63 Supplementary. Heterozygosity samples of western lowland gorillas at 20X (downsampled) by sampling site. Box-plot colors represent the three main PCA clusters: Dzain pink, south of Gabon in dark red and the rest in yellow.

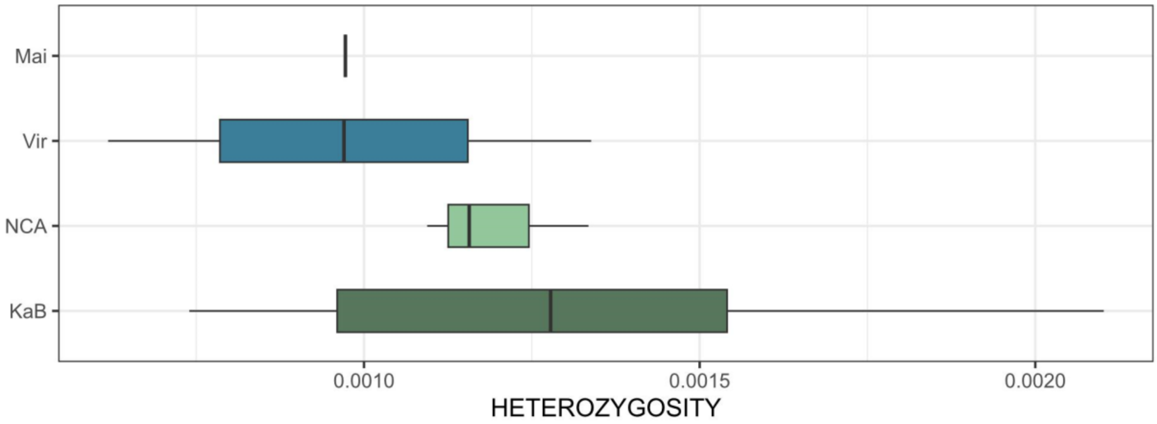


Figure 64 Supplementary. Heterozygosity samples of eastern gorillas at 20X (downsampled) by sampling site. Blue colors showed mountain gorilla sites, and green colors showed eastern lowland sites.

Fewer individuals per social group were retained after filtering by high coverage (minimum 20X). The dispersion of mean values inside the KaB site was again observed, with Ganywamulme and Mankoto showing the highest heterozygosity values, and Mugarukathe the lowest (Figure 65 Supplementary). Mountain gorillas also showed similar patterns of what was observed using genotype likelihoods (Figure 66 Supplementary).

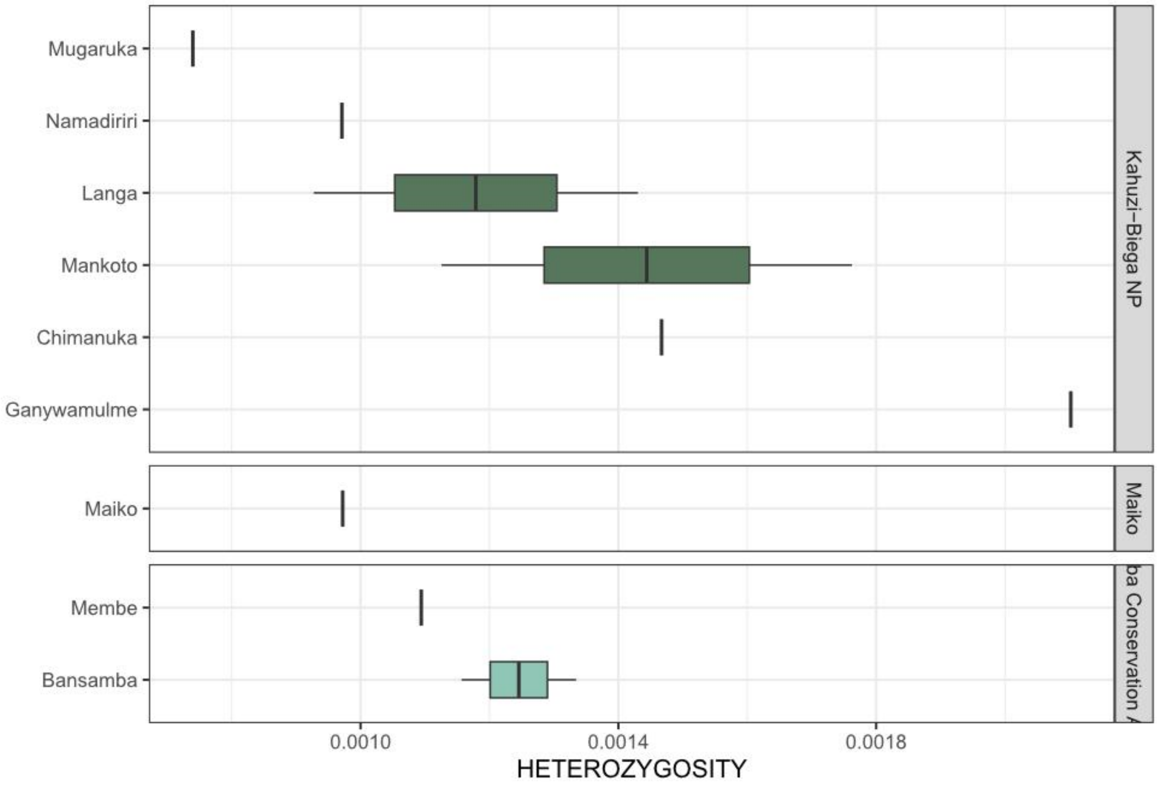


Figure 65 Supplementary. Heterozygosity samples of eastern lowland gorillas at 20X (downsampled) by social groups. Different social groups are colored by different green colors (social groups from KaB National Park are colored in dark green, Nkuba Conservation Area (NCA) social groups in blueish-green and Membe social groups in light green).

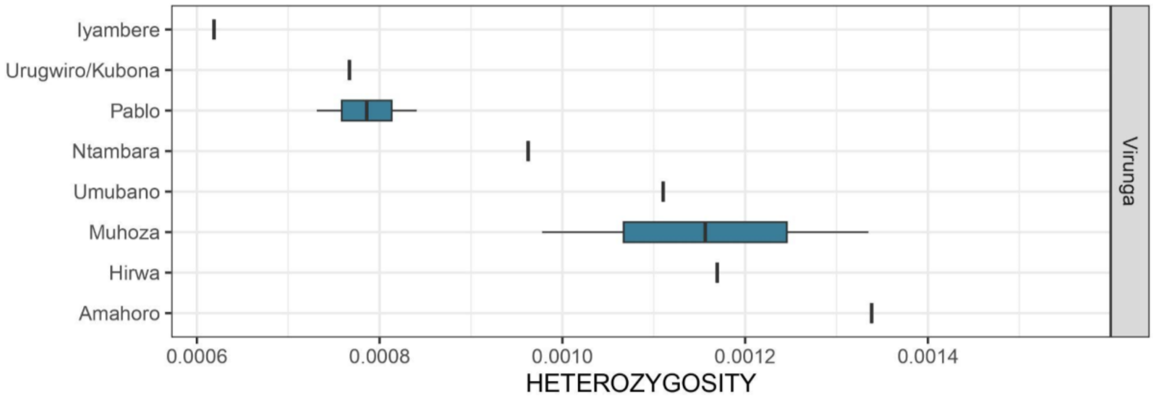


Figure 66 Supplementary. Heterozygosity samples of the Virunga population (mountain gorillas) at 20X (downsampled) by social groups.

#### 3.4 FST

Genetic population differentiation was tested through FST values using chromosome 21 and exome data separately. For both cases, 1D-SFS was calculated for the subspecies (CRG, ELG, MG, WLG) using ANGSD v.0.935 and realSFS. We estimated the 2D-SFS for each subspecies comparison using realSFS with the site allele frequency files as input. To calculate pairwise Fst, the 2D-SFS were used as priors, and the site allele frequencies files were input to realSFS with the command ‘fst index’ to generate fst binary files. We specified the flag ‘-whichFst 1’ to calculate Bhatia’s estimator of Fst (Bhatia et al. 2013).

Chromosome 21:

FST values using chromosome 21 data were calculated between subspecies, using PCA clusters as groups for western lowland gorillas and between sites. In order to avoid sample bias, all groups were downsampled to the same number of individuals as the group having the lowest value.

FST values between subspecies showed lower genetic distance between the western subspecies than the eastern subspecies (Figure 67A Supplementary). When comparing between PCA clusters, the Virunga population showed lower genetic distance to eastern lowland gorillas than the Bwindi population. Cross River gorillas showed the highest genetic distance to the eastern species in comparison to western lowland gorillas of different groups (Figure 67B Supplementary). PCA cluster separated western lowland individuals in 4 different groups (Figure 36 Supplementary): samples from the south of Gabon (GabonS), samples from Dza and Gou (CAR-RC), samples from Cameroon (Cameroon) and samples from the rest of Gabon (Gabon N). The samples from the south of Gabon were the ones more genetically distant to the Cross River gorillas. The Dza and Gou (CAR-RC) group was the group more genetically distant to the rest of the western lowland groups (Figure 68 Supplementary).

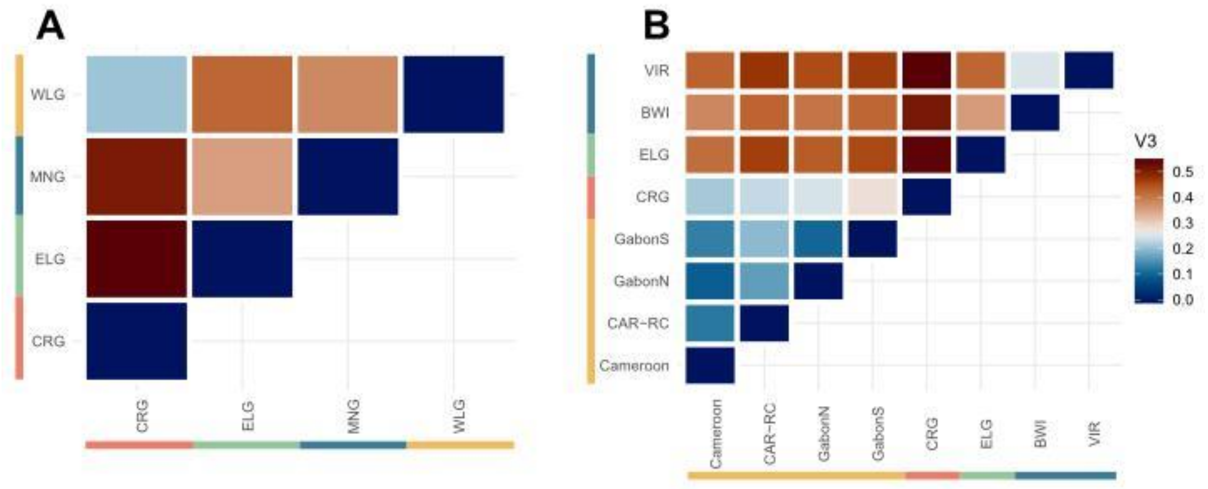


Figure 67 Supplementary. FST per subspecies (A) and per PCA cluster (B) downsampled to 18 individuals.

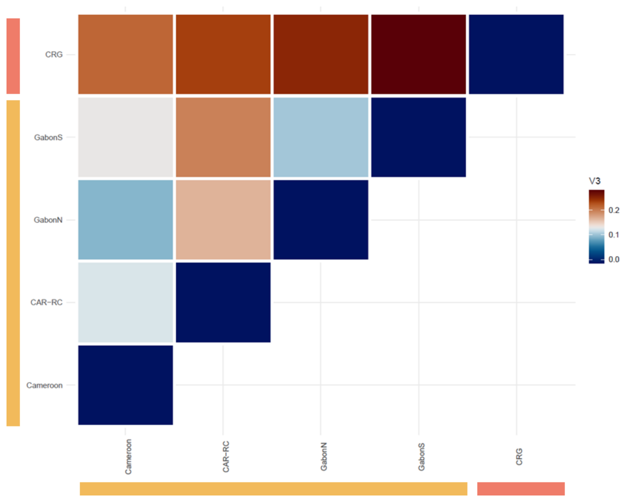


Figure 68 Supplementary. FST per PCA cluster for western gorillas downsampled to 18 individuals.

FST was also performed by sampling site. The minimum number of individuals used per site was 8 to better represent allele frequencies of each population. Sampling sites closer than 25 kilometers and no clear geographic barrier present between them were merged and studied as a unified population.

FST per sampling site showed similar results obtained for subspecies and PCA clusters at the genus level (Figure 69 Supplementary). Increasing the resolution at the population level, it was possible to better detect genetic distances inside and between specific regions of gorilla subspecies. Nga-Dib-Dze, Lok-Imb, Mou, and Nki-Mes (ordered from highest to lowest values of FST) were the sites with higher genetic distance to the Cross River gorillas. All the rest of the western lowland gorilla populations with Cross River subspecies had lower values of FST, lower than Bwindi and Virunga populations from the same subspecies (Figure 70 Supplementary).

Inside western lowland populations, lower values of genetic distance were found between Lob-Mam, Nki-Mes, Bel and Den (Figure 71 Supplementary).

For eastern lowland gorillas, only one site had at least 8 individuals to be included in the analyses (KaB). Between the eastern gorilla populations, KaB had lower values of FST with Bwindi than Virunga (Figure 72 Supplementary).

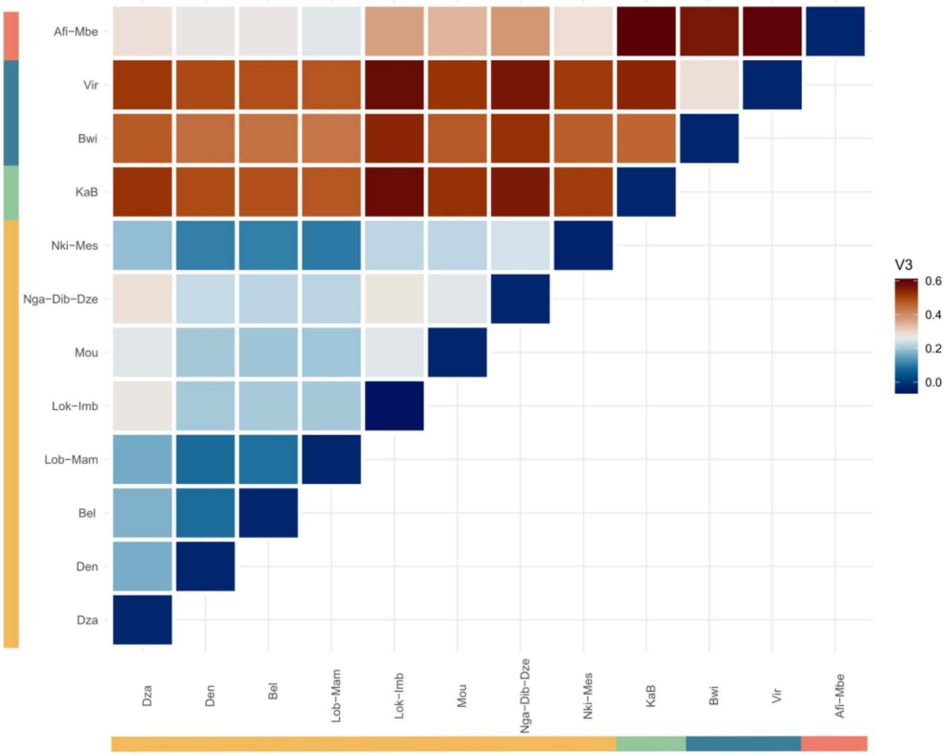


Figure 69 Supplementary. FST using all gorilla sites > 8 individuals (Afi-Mbe, Nki-Mes, Nga-Dib-Dze, Mou, Lok-Imb, Lob-Mam, Bel, Den, Dza, Vir, Bwi and KaB).

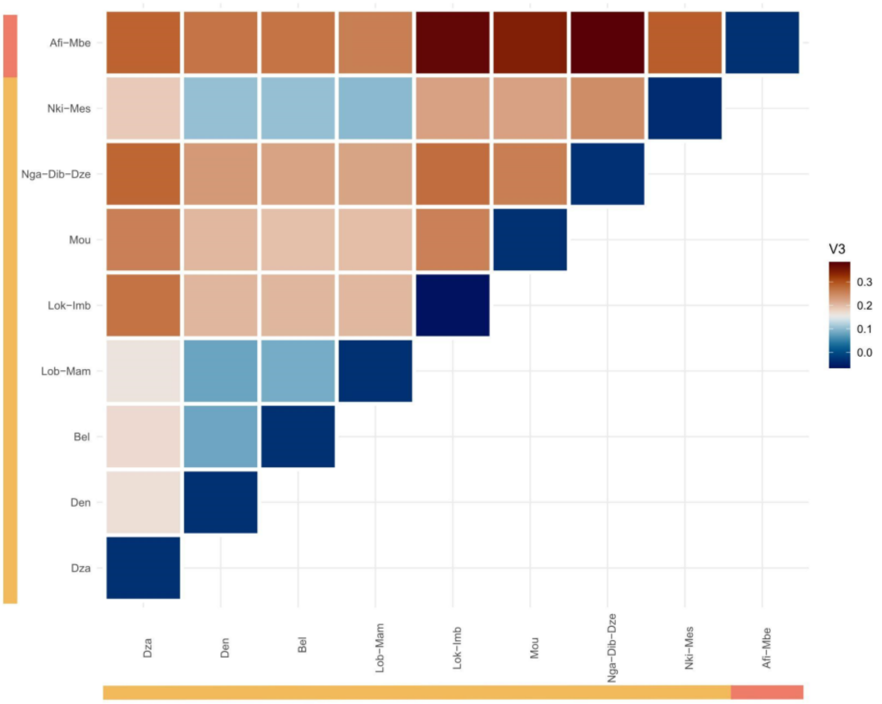


Figure 70 Supplementary. FST using western gorilla sites > 8 individuals (Afi-Mbe, Nki-Mes, Nga-Dib-Dze, Mou, Lok-Imb, Lob-Mam, Bel, Den and Dza)

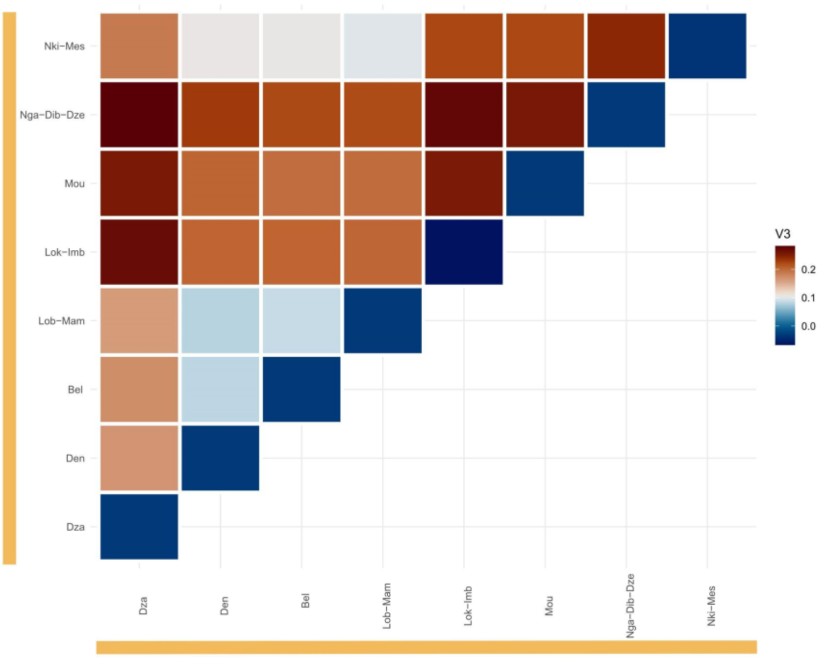


Figure 71 Supplementary. FST using western lowland gorilla sites > 8 individuals (Nki-Mes, Nga-Dib-Dze, Mou, Lok-Imb, Lob-Mam, Bel, Den and Dza).

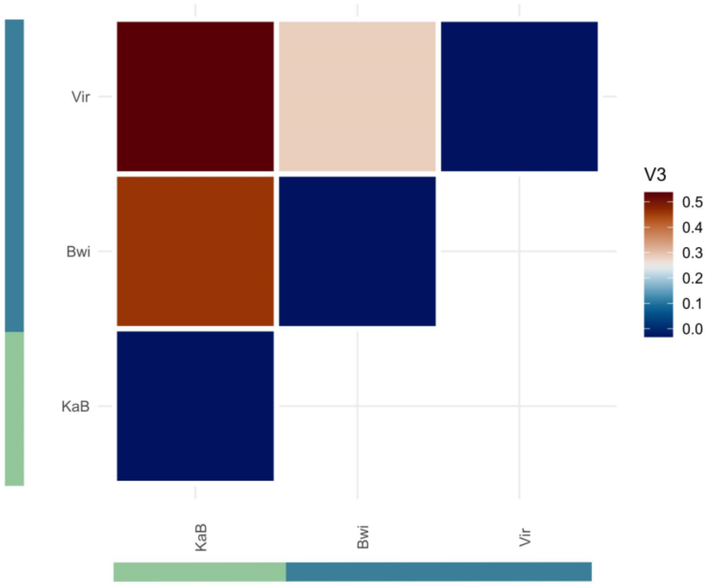


Figure 72 Supplementary. FST using eastern gorilla sites > 8 individuals (Virugna, Bwindi and Kahuzi-Biega)

### 4. Population connectivity and gene flow

Population connectivity and dynamics, and possible gene flow were assessed using D-statistic analysis and admixfrog, based on genotype likelihoods, and EEMS, using called variants.

#### 4.1 D statistics

A minimum of 8 individuals per population were used in this analysis; a few sites were merged based on proximity in order to have enough individuals to better represent the population (same as used for FST analysis).

The highest value of allele sharing between Cross River gorillas and other western lowland populations was found for Dza, followed by a cline of positive values for Lob-Mam, Nki-Mes, Bel, and Den (Figure 73 Supplementary). The sites that did not show a positive signal of an access of allele sharing with Cross River gorillas were Nga-Dib-Dze, Mou, and Lok-Imb.

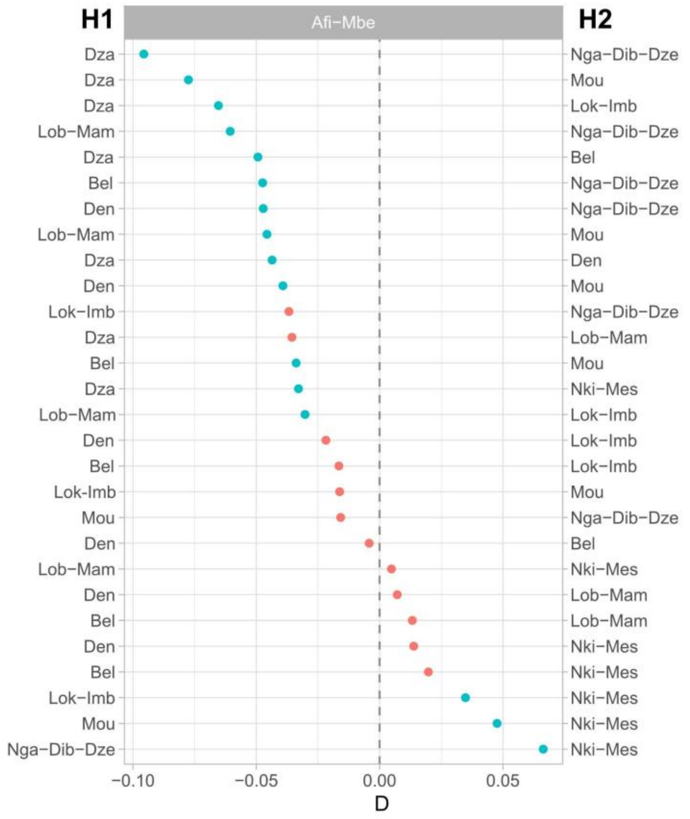


Figure 73 Supplementary. D-STATS (ABBA-BABA) using ANGSD between Cross River populations and WLG populations. Sites having more than 8 individuals were used. Geographically close sites were merged and used as a unique site for this analysis.

#### 4.2 Fragments of shared ancestry

Fragments of shared ancestry were inferred using admixfrog software on chromosome 21 data. This program uses an HMM to infer ancestry fragments from low-coverage and contaminated data. The ancestral states used were: CRG, WLG, ELG, MNG, and HUM. Each genome was divided into 35bp size bins, and then these bins were assigned to one of the ancestral states previously defined. The number of SNPs and the posterior probabilities (Figure 74 Supplementary) of each studied bin were used to filter them out (> 0.9 posterior probability).

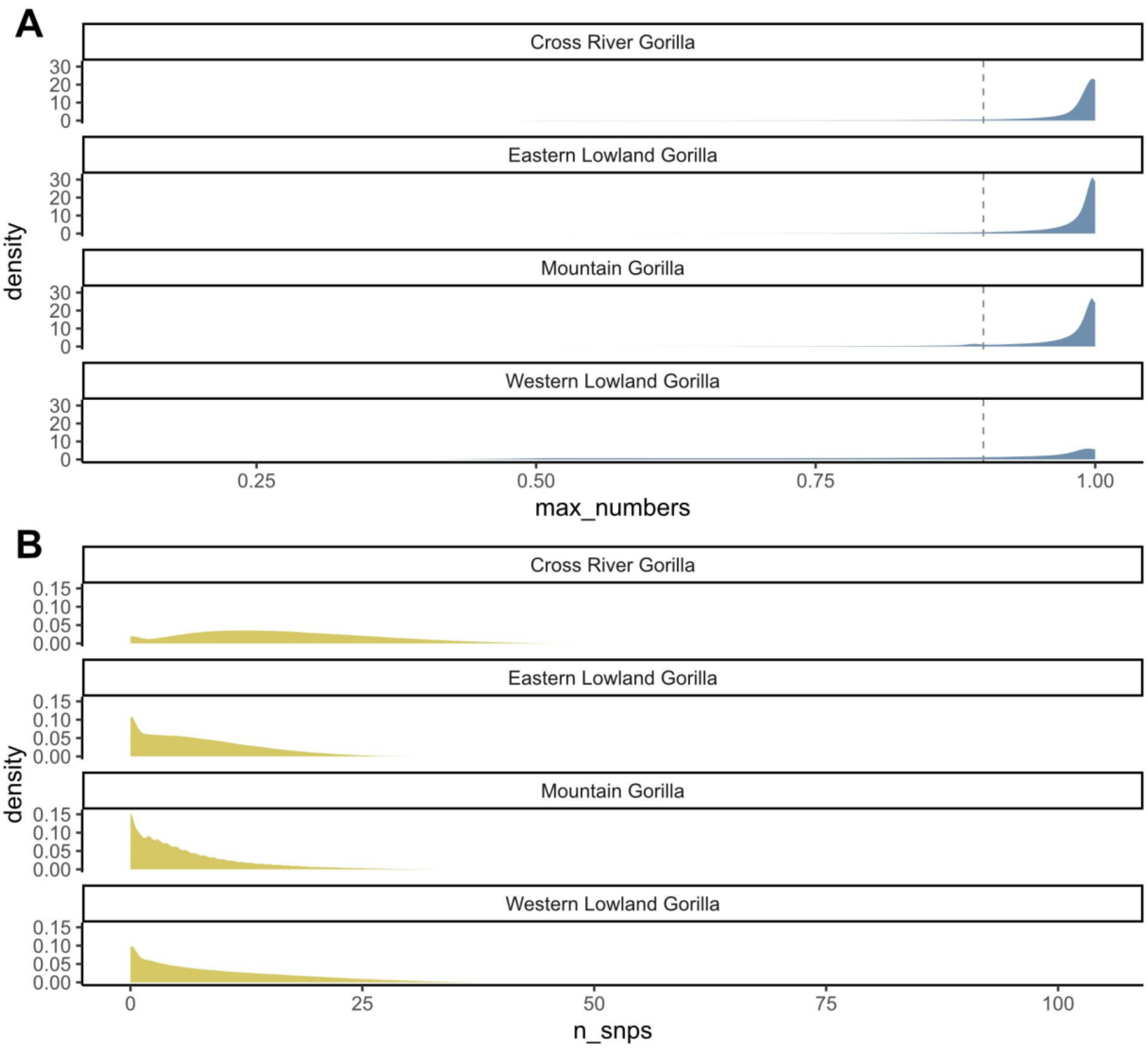


Figure 74 Supplementary. Distribution plots of the number of SNPs used per bin(A) and the number of priors used per bin(B) represented by subspecies. These are the number of SNPs used for each bin when assigning the ancestral states, and the posterior probabilities obtained.

The total number of bins for each ancestral state and each sample were counted after filtering in order to compare between subspecies and populations (Figure 75 Supplementary). The total amount of bins for each sample varies due to the filtering applied. For each subspecies the majority of bins were assigned to the same subspecies ancestral state. Some variation was seen inside western lowland populations.

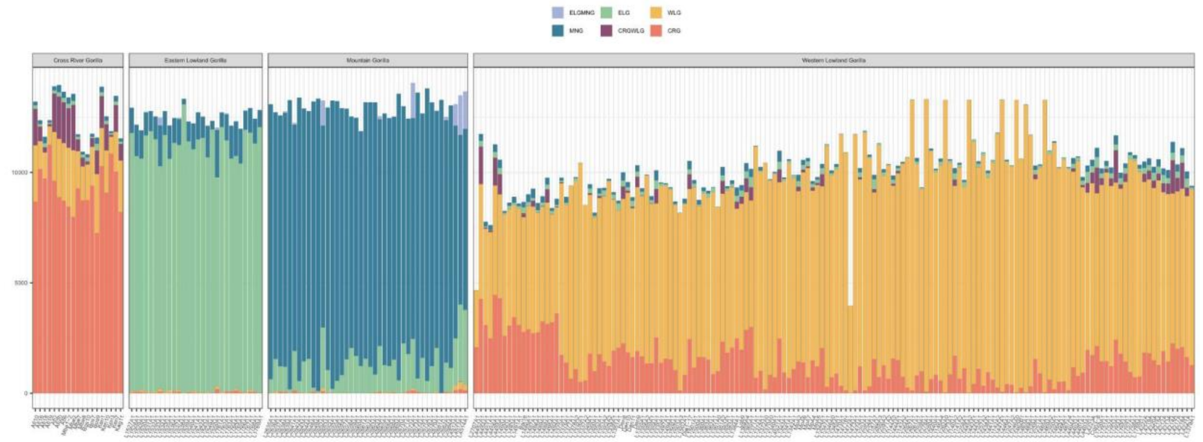


Figure 75 Supplementary. Number of bins assigned to each ancestral state. Each bar represents a unique sample. Samples are divided by subspecies: Cross River gorillas, eastern lowland gorillas, mountain gorillas and western lowland gorillas (from left to right). Each ancestral state is distinguished by different colors as indicated in the legend (western lowland gorilla in yellow, Cross River gorilla in red, mountain gorilla in blue, and eastern lowland gorilla in green).

Ancestry state-specific tracts were inferred for each sample. The sum of all ancestry tracts for each ancestral state was calculated and plotted by subspecies for each possible ancestral state (Figure 76 Supplementary). As expected, for each subspecies, the highest number of ancestry tracts detected belonged to their subspecies. The total length of tracts detected for the WLGCRG ancestral state was higher for CRG samples than for the WLG.

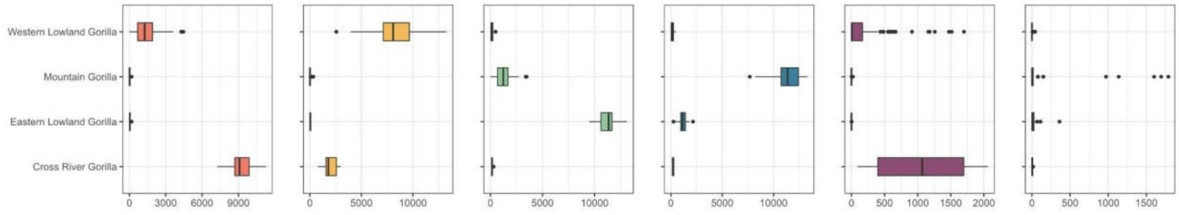


Figure 76 Supplementary. Ancestry tracts by subspecies. Each ancestry tract is represented by a unique color (add colors). The ancestry tracts are represented as the sum of all ancestry tracts found in each sample.

When checking the amount (sum of all CRG ancestry tracts) of ancestry tracts detected, which belong to the CRG ancestry for the different western lowland gorilla groups, differences were detected between them (Figures 77-78 Supplementary). Samples from Dza and Gou were the ones with the most  CRG ancestry tracts. On the contrary, inside the "Central" group, samples from Odzala National Park (Nga-Dib-Dze and Lok-Imb) were the ones having lower values of CRG ancestry tracts (Figure 78 Supplementary). In addition, the WLGCRG ancestry was more detected in samples also having a higher value of CRG ancestry tracts (Figure 77 Supplementary).

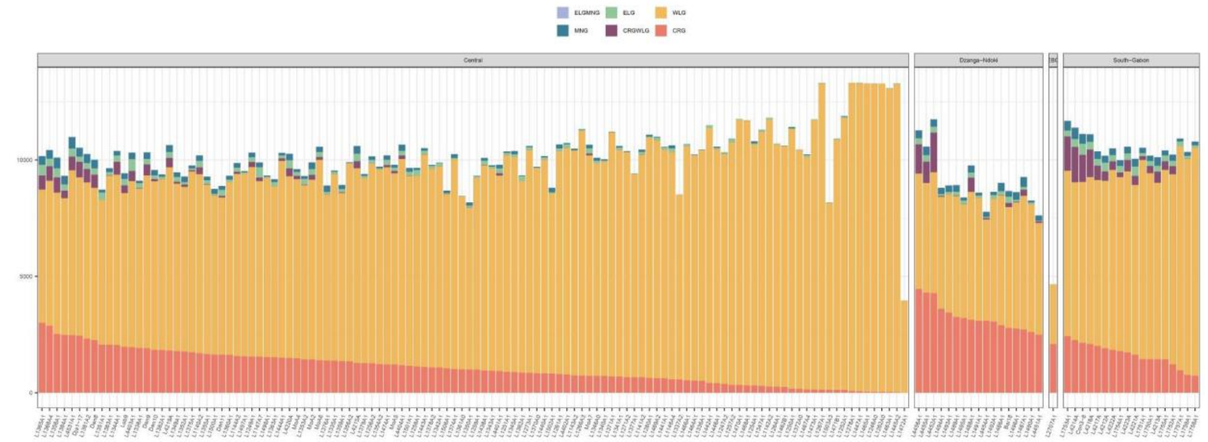


Figure 77 Supplementary. Ancestry tracts for western lowland gorillas by PCA clustering. Each ancestry tract is represented by a unique color (western lowland gorilla in yellow, Cross River gorilla in red, mountain gorilla in blue, and eastern lowland gorilla in green). The ancestry tracts are represented as the sum of all ancestry tracts found in each sample.

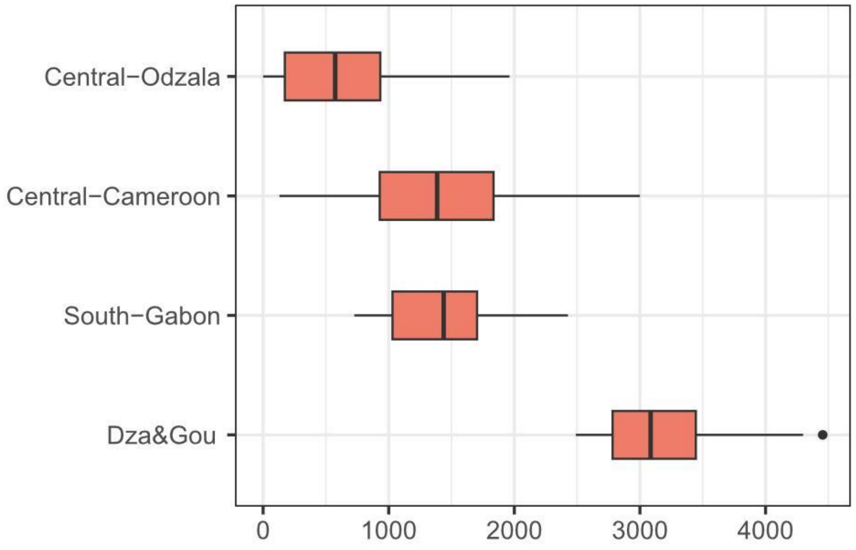


Figure 78 Supplementary. CRG ancestry tracts for western lowland gorillas by groups, including Ebo Forest sample.

We also checked the distribution across chromosome 21 of the detected Cross River tracts in western lowland individuals (Figures 79-80 Supplementary).

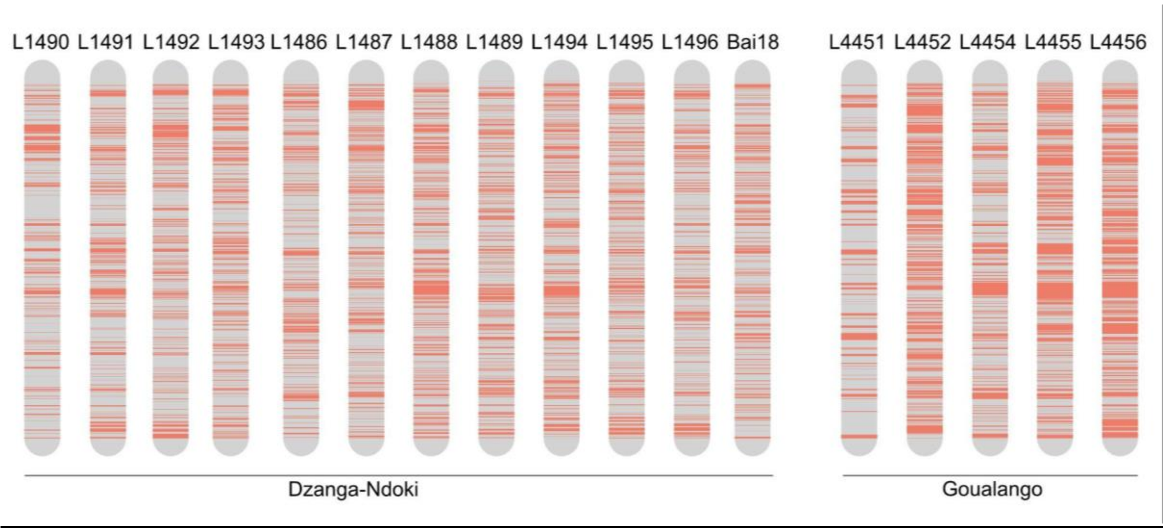


Figure 79 Supplementary. Cross River ancestry tracts across chromosome 21 for Dzanga-Ndoki and Goualougo sites.

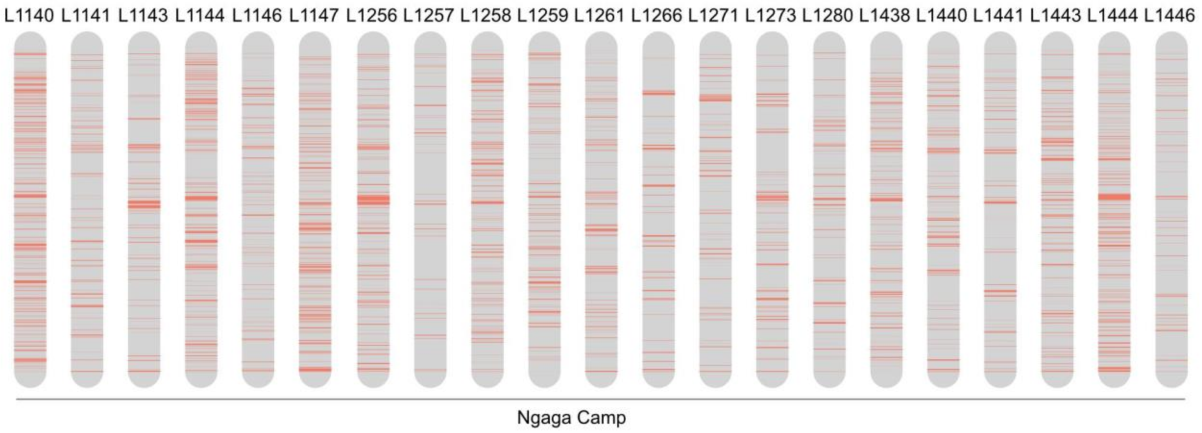


Figure 80 Supplementary. Cross River ancestry tracts across chromosome 21 for the Nga-Dib-Dze site.

Inside Cross River gorilla samples, no specific differences were detected between sampling sites (Figure 81 Supplementary).

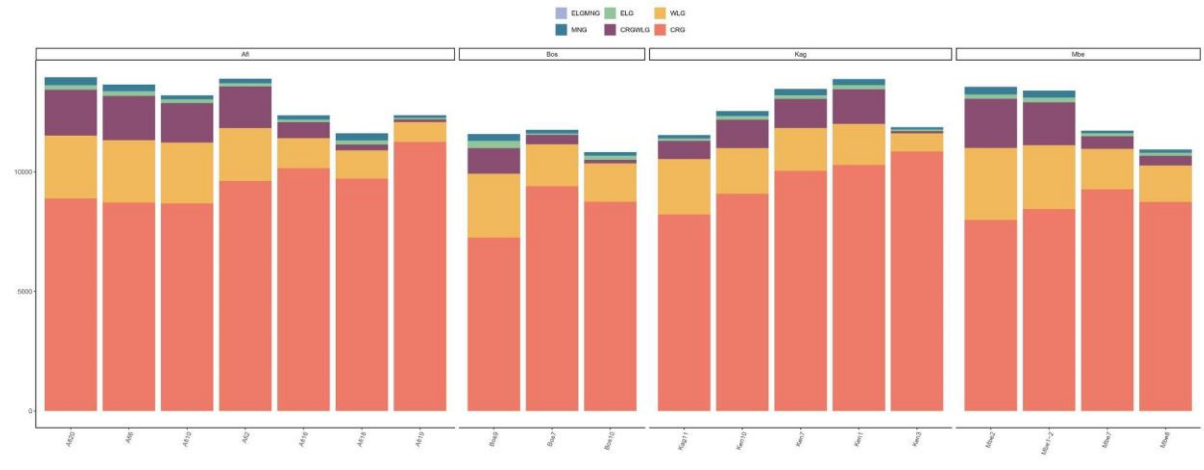

Figure 81 Supplementary. Ancestry tracts for Cross River gorillas by site. Each ancestry tract is represented by a unique color (western lowland gorilla in yellow, Cross River gorilla in red, mountain gorilla in blue, and eastern lowland gorilla in green). The ancestry tracts are represented as the sum of all ancestry tracts found in each sample.

Virunga and Bwindi showed differences when comparing the number of ELG ancestry tracts detected. The Bwindi population had higher values of ELG ancestry tracts (Figure 82-83 Supplementary). As seen for the western species, samples having higher amounts of ELG ancestral tracts also showed higher values of ELGMNG ancestral tracts.

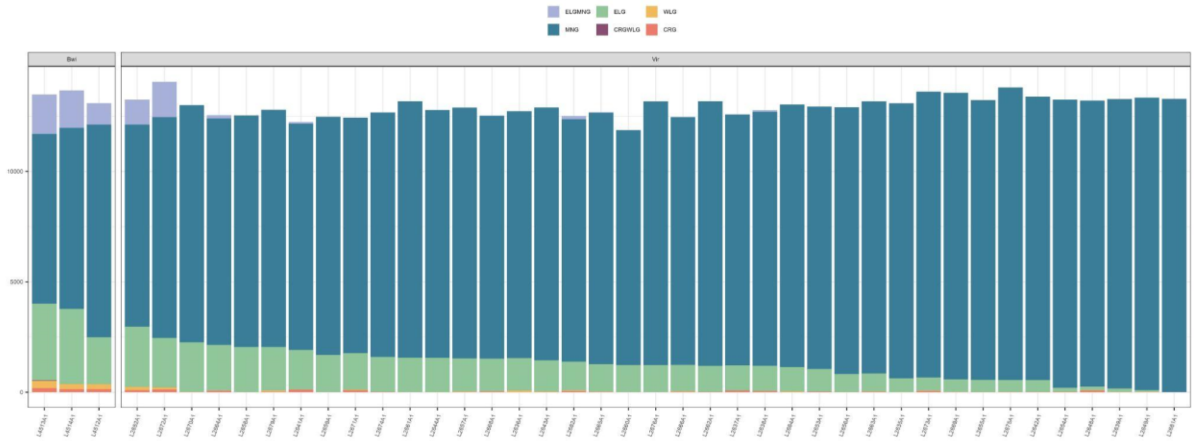


Figure 82 Supplementary. Ancestry tracts for mountain gorillas by site (Virunga and Bwindi). Each ancestry tract is represented by a unique color (western lowland gorilla in yellow, Cross River gorilla in red, mountain gorilla in blue, and eastern lowland gorilla in green). The ancestry tracts are represented as the sum of all ancestry tracts found in each sample.

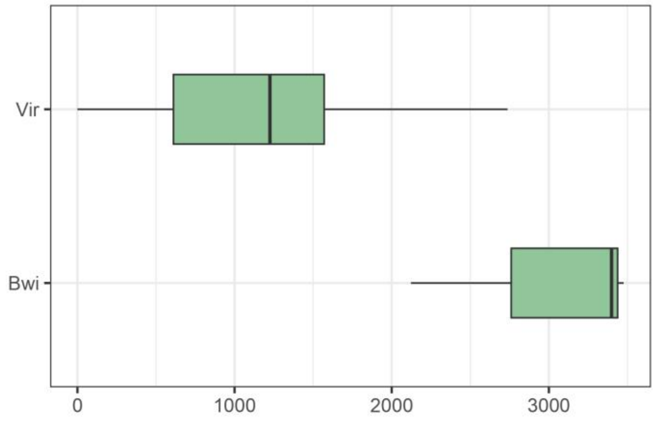


Figure 83 Supplementary. ELG ancestry tracts for mountain gorillas by site (Virunga and Bwindi).

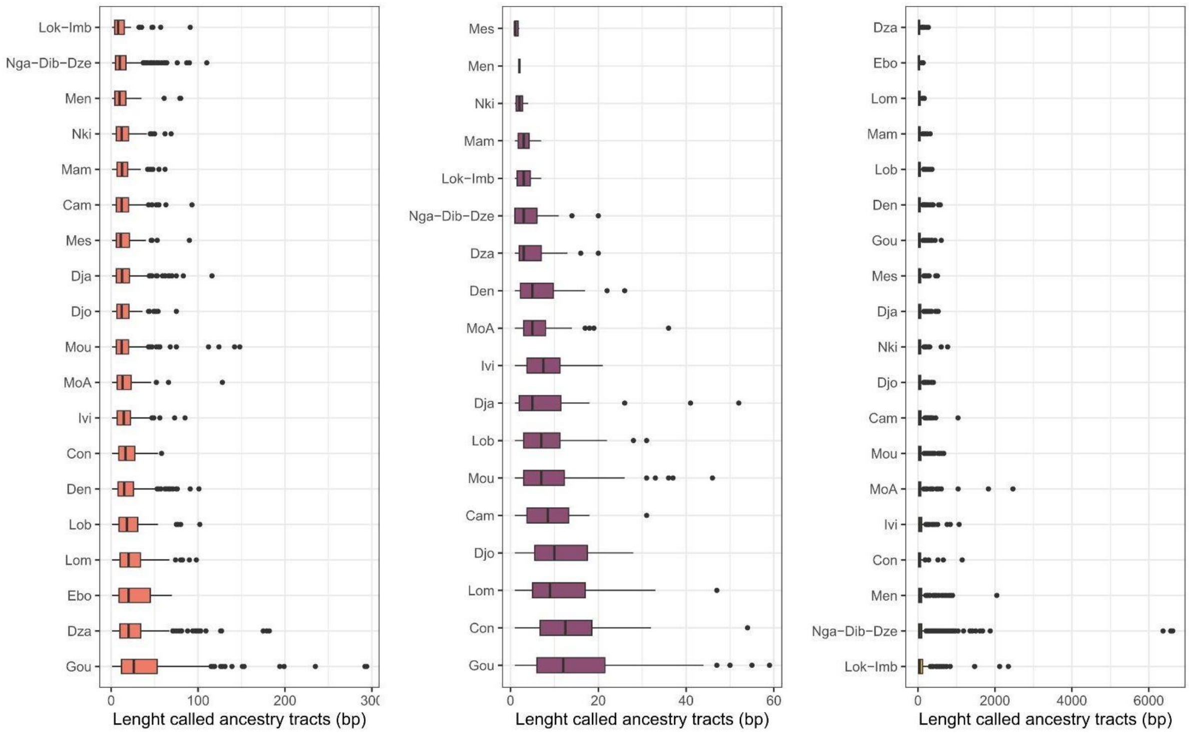


Figure 84 Supplementary. Tract length per sampling site and ancestry STATE (A-CRG; B-WLGCRG; C-WLG).

### 4.3 EEMS

Population dynamics and connectivity was studied through EEMS (Estimated Effective Migration Surfaces). Deviations from isolation-by-distance were assessed using EEMS, based on samples with specific geographic coordinates, more than 5X of coverage, and less than 40% of missing data. The analysis was performed at genus and species level.

At genus level it was possible to observe a reduction of the migration rates between species and to a lesser extent between subspecies (Figure 85 Supplementary). When applying EEMS at subspecies level it was possible to better detect the reduction in the migration rates between species and inside the western lowland gorillas between some populations.

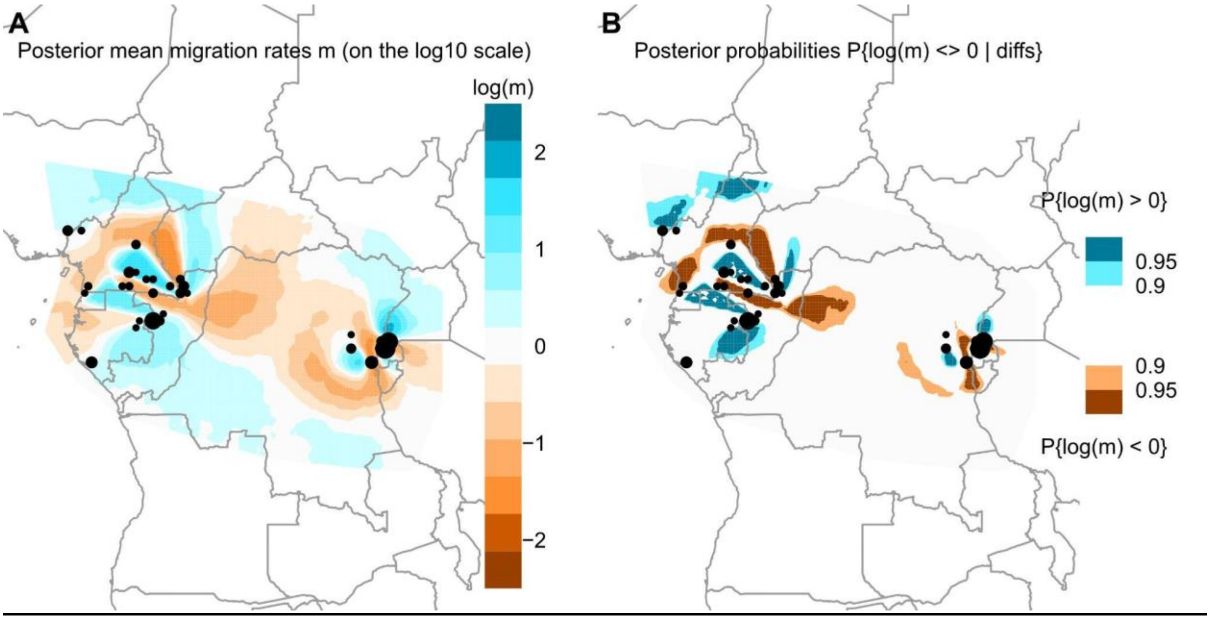


Figure 85 Supplementary. Effective Migration plots for chromosome 21 data of all gorilla samples above 5X of coverage and less than 40% of missing data. A. Posterior mean migration rates. B. Posterior probabilities.

For mountain gorillas, it was possible to see the reduced migration rates between the two isolated populations, Virunga and Bwindi (Figure 86 Supplementary). Inside the eastern lowland gorilla subspecies, no pattern was observed as few sampling sites were retained after filtering the dataset for this analysis. More samples with more quality and at higher coverage will be needed in order to assess deviation from the isolation-by-distance inside this subspecies.

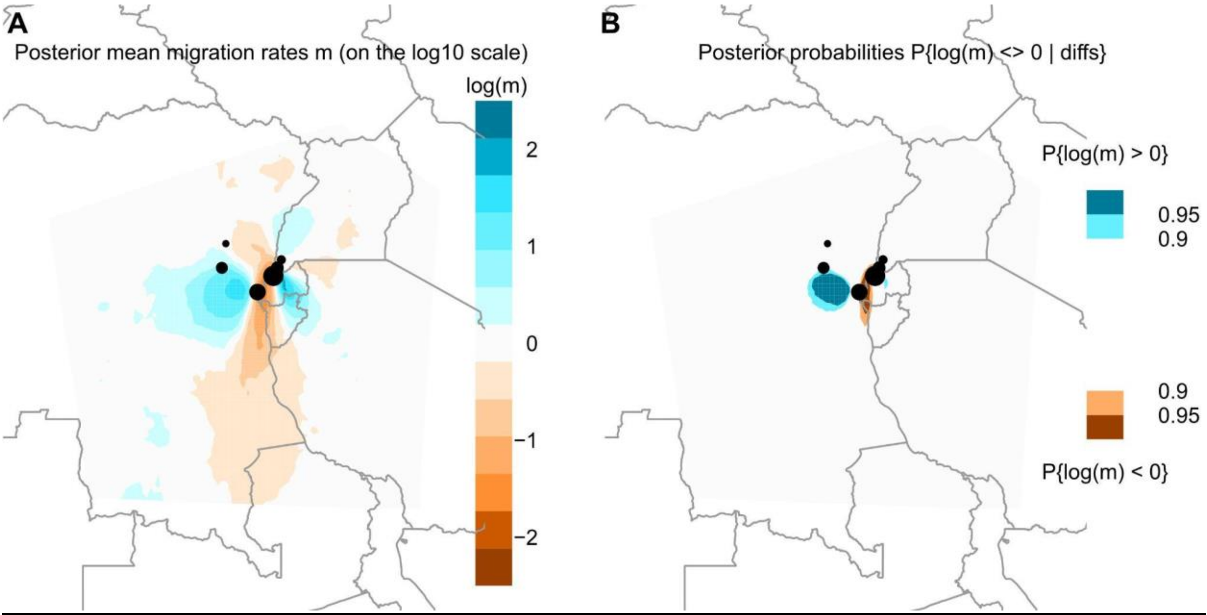


Figure 86 Supplementary. Effective Migration plots for eastern gorilla species (eastern lowland gorillas and mountain gorillas) above 5X of coverage and less than 40% of missing data. A. Posterior mean migration rates. B. Posterior probabilities.

The Sanaga river which acts as a geographic barrier was detected in EEMS analysis as a reduction in the migration rate between the two western gorilla subspecies (Figure 87 Supplementary). Within the western lowland subspecies different deviations from the isolation-by-distance model were also observed. The Sangha river and its affluents (Kadei and Ngoko rivers) were detected as genetic barriers between Gou and Dza sampling sites and the rest of the western lowland gorilla populations. In addition, Dja river also was detected as a barrier decreasing the migration rate between the sampling sites from Cameroon located above (Lob, Mam, Bel, Den, Nki, Mes, Lom) and below (Djo and Men) the Dja river. The Ogoué river was also detected acting as a barrier, but it was not significant as shown in the posterior probabilities plot (Figure 87B Supplementary). A unique sampling site (Mongambe) from the south of Gabon was retained after filtering. More samples from different populations from this area are needed in order to better detect possible deviations from the isolation-by-distance model. Finally, three different areas were detected with increased migration rates. The first one was detected in the border of the Republic of Congo and Gabon, between sampling sites from Nga-Dib-Dze in the Odzala National Park and one sampling site from Gabon (Ivi). In the east part of Cameroon distribution range was detected another possible genetic corridor between Den, Bel, Lob, Mam, and Lom. Finally, an inter-subspecies possible genetic corridor was detected between Gou and Dza sampling sites and Cross River gorilla populations.

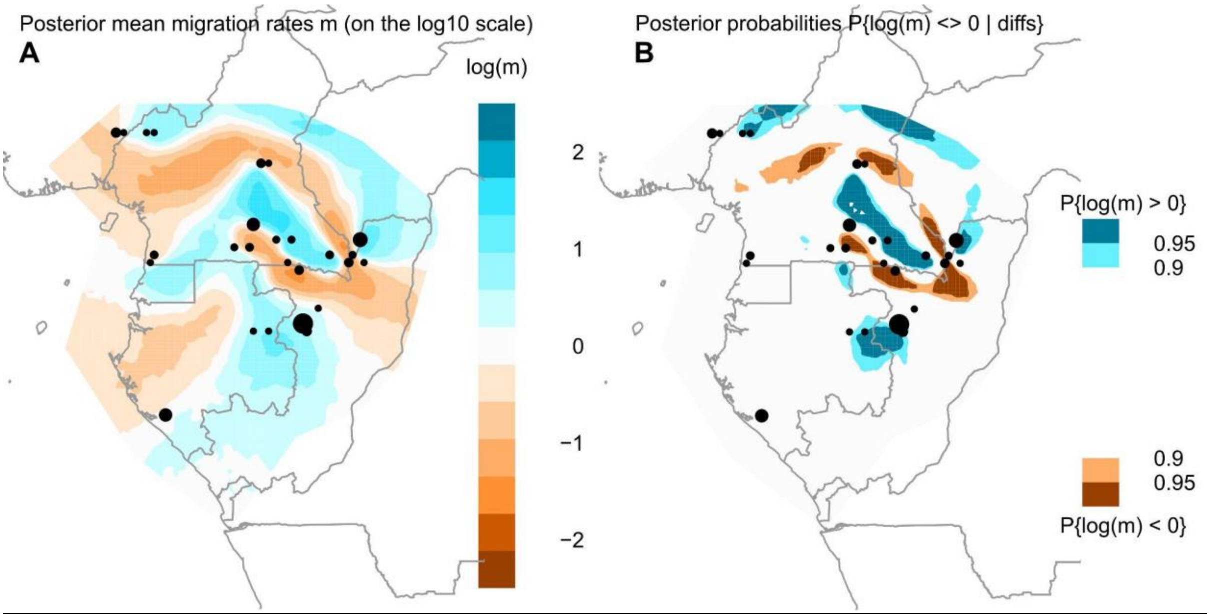


Figure 87 Supplementary . Effective Migration plots for western gorilla species (western lowland gorillas and Cross River gorillas) samples above 5X of coverage and less than 40% of missing data. A. Posterior mean migration rates. B. Posterior probabilities.

### 5. Geolocalisation

We tested two different methodologies to predict the geographic coordinates using our chromosome 21 data. Once the comparison was done, we also predicted geographic coordinates of public wild-born gorilla genomes for which sample origin information was not available. In addition, we also predicted geographic coordinates for the newly sequenced FTA cards and museum samples.

Locator

First, we applied the method Locator (Battey et al. 2020), a machine learning method developed to predict geographic coordinates, which has been applied to a range of species, including humans and mosquitoes. We used it to the georeferenced dataset of gorilla non-invasive samples (taking samples with coverage >5X). We applied Locator to predict geographic coordinates for the 169 georeferenced non-invasive samples (157 faecal samples (including 3 faecal samples from Fontsere et al. 2022), 12 hair samples (including hair samples from Alvarez-Estape et al. 2023), consisting of 70 eastern gorillas and 99 western gorillas). Note that we applied the method separately for eastern and western gorillas. In the input file, we assigned the coordinates of the test sample to NA (missing) in a leave-one-out process. This approach was applied to all the samples in the dataset one by one. We calculated the difference between the predicted and geographic location for each sample. We visualised this using distribution plots to evaluate the overall prediction of all samples for each sampling site. We assessed uncertainty in the predictions using bootstrapping [--bootstrap --nboots 50] which generates 50 replicates. We calculated the correlation between the difference in predicted and geographic location (in kilometres) and the number of samples per sampling site. We also calculated the correlation between the difference in predicted and geographic location (in kilometres) and levels of missingness per sample.

We assessed the impact of the number of samples at a sampling site on the geographic predictions made by Locator, as well as the impact of missing genotypes (Figure 88 Supplementary). The number of samples per site, as expected, influenced the kilometers of error when predicting the geographic coordinates (not significant correlation). On the other hand, the amount of missing data in the test samples had a highly significant correlation in the kilometers of errors of the predicted geographic coordinates. This significant correlation highlighted the importance of good-quality DNA in the tested genome. The Nga-Dib-Dze sampling site showed high differences between samples in the kilometers of error due to the high amount of missing data found in some of the samples. For Nga-Dib-Dze samples having missing data, their mean difference between true and predicted location was 427.65 km, versus Nga-Dib-Dze samples without missing data, for which the mean difference between true and predicted location was 22.48 km (Figure 88B Supplementary).

Differences between subspecies in the predictions were detected (Figure 88 Supplementary). Cross River gorilla sampling sites showed low numbers of individuals per site and high missing data after filtering, negatively affecting the final geographic predictions in comparison to the rest of subspecies.

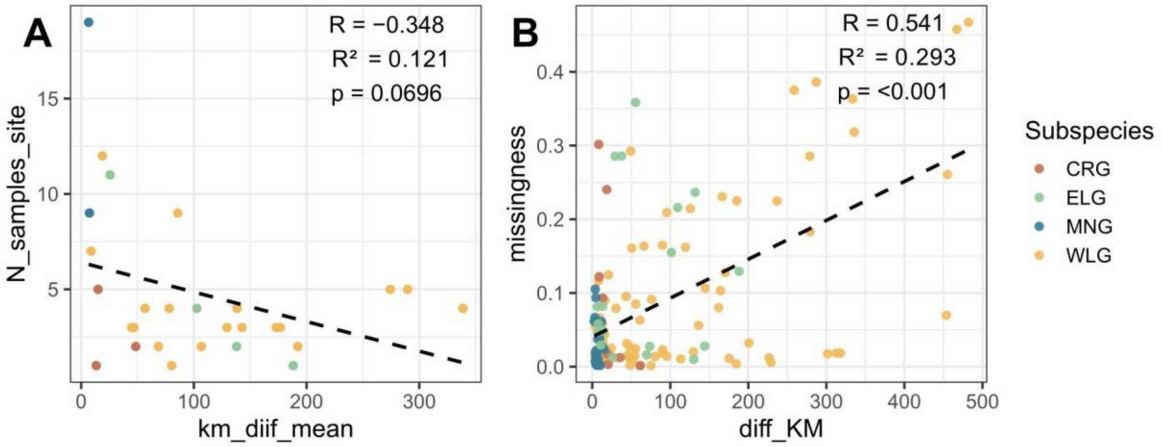


Figure 88 Supplementary. Assessment of the impact of missingness and number of samples per site on the performance of the Locator method (Battey et al. 2020) for gorilla samples. A) Correlation plot indicating the mean kilometres of error and number of samples per site. The number of samples per site counted was the one used in this analysis. B) Correlation plot indicating the kilometers of error and missing data for each sample used in this analysis.

As mentioned, geographic predictions were also influenced by genetic differentiation and population structure. An example was found for the Den western lowland gorilla population. Despite the high geographic distance to other western lowland populations (~240-430 km), this sampling site had low values of FST with those western lowland populations: Lob, Mam, Bel and Cam.

<u>PCA points distances:</u>

Second, subspecies-specific PCA were used in order to predict geographic coordinates from the nearest points for which GPS coordinates were available. Specifically, we used the point distances of PC1 and PC2 of each PCA (Figures 36, 39, 41 and 42 Supplementary). The four nearest neighbour points of the tested sample were used to predict the geographic coordinates, applying a mean weighted by the Euclidean distances of each point to the tested sample point and the amount of difference explained in each of the two first components used.

<u>Comparison between methods:</u>

We compared the performance per site of both methods to determine which one of the two methods performed better using our chromosome 21 dataset. We used the prediction error (kilometers of difference between the real and the predicted location) to compare both methodologies (Figure 89 Supplementary).

Within the western lowland gorilla sites, similar trends between both methods were found for the majority of sites. Sampling sites for which the prediction errors were low (Dza, Nga-Dib-Dze, Mou, Gou), the accuracy of prediction were higher when applying Locator for Dza and Nga-Dib-Dze camp (Figure 89 Supplementary), in contrast to Mou and Gou, for which the predictions were better when using PCA point proximity or equal between the two methods respectively. On the other hand, regarding the sampling sites which had the highest values of prediction errors, the performance was improved when using PCA point distances (Cam, Den, Ivi, Bel and Djo). Lob, Mam, Nki and Mes were the only four sites for which the performance was better when using Locator. Remarkably, when using Locator only samples above 5X of coverage and low missing data were used, losing all samples from Mont Allen (the unique site from Equatorial Guinea). For other sites, the number of samples used was also reduced due to the filtering when using Locator.

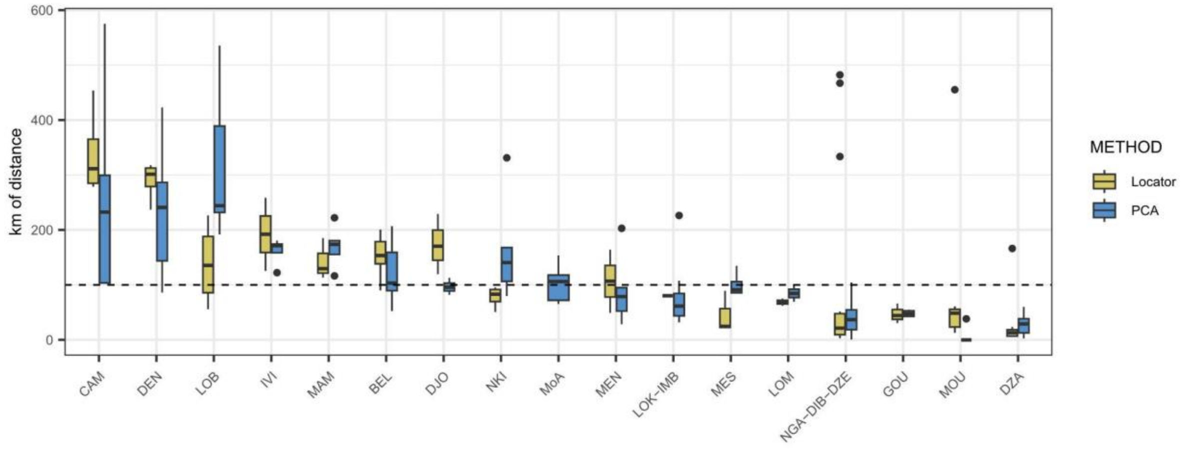


Figure 89 Supplementary. Comparison between two geolocalization methods (Locator and PCA point proximity) using western lowland gorilla chromosome 21 data. Each box-plot represents a unique site of the western lowland distribution. The green box-plot showed Locator prediction errors, and the blue box-plots showed prediction errors when applying PCA point distances. Prediction errors are calculated using the kilometers of distance between the true and the predicted location of each sample for each sampling site.

When comparing methods between the Cross River and the eastern lowland gorilla subspecies, higher differences were found (Figures 90A and C Supplementary). The accuracy when using Locator for these two subspecies was low due to the fact of missing sampling sites, a low number of samples per site and high missing data. For mountain gorillas, the numbers were similar between both methods (Figure 90B Supplementary).

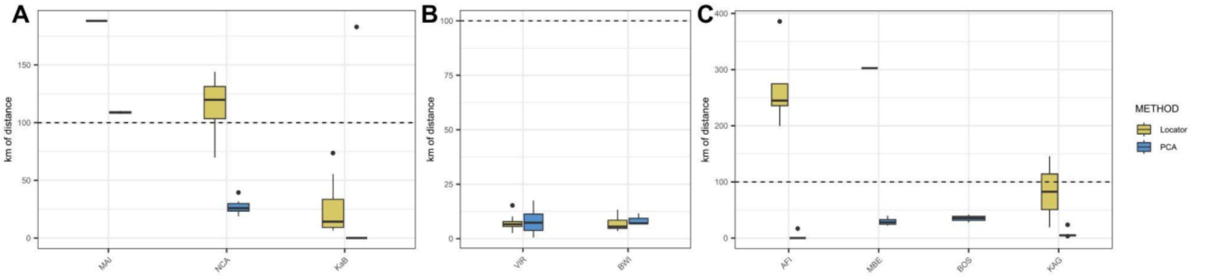


Figure 90 Supplementary. Comparison between two geolocalization methods (Locator and PCA point proximity) using chromosome 21 data. A) Comparison of eastern lowland gorilla sites. B) Comparison of mountain gorilla sites. C) Comparison of Cross River gorilla sites. The green box-plot showed Locator prediction errors, and the blue box-plots showed prediction errors when applying PCA point distances. Prediction errors are calculated using the kilometers of distance between the true and the predicted location of each sample for each sampling site.

As more sampling sites and the number of samples per site were higher, we decided to apply the PCA point proximity method on previously published wild-born gorilla genomes for which the origin information was not available (Prado-Martinez et al. 2013a, Prado-Martinez et al. 2013b, Xue et al. 2015, Pawar et al. 2023).

Regarding the eastern lowland gorillas, geographic coordinates were predicted for individuals for which the origin information was unknown (Figure 91A Supplementary). Gbg-Itebero geographic coordinates were predicted near the Nkuba Conservation Area. The other three individual geographic coordinates were predicted near the Walikale site, with Gbg-Serufuli being the closest one. The rest of the eastern lowland public genomes (Supplementary table 3) with known origin were also used as a reference dataset in the PCA to predict geographic coordinates.

Public mountain gorilla samples (Xue et al. 2015, Pawar et al. 2023) had population origin information available. Using this information, we check the predicted geographic coordinates for these samples using PCA point distances in our new chromosome 21 dataset. All samples were correctly assigned to the corresponding population (Bwindi or Virunga) (Figure 91B Supplementary). Population structure and high genetic differentiation characteristic of this subspecies could help the accuracy and precision of the predictions.

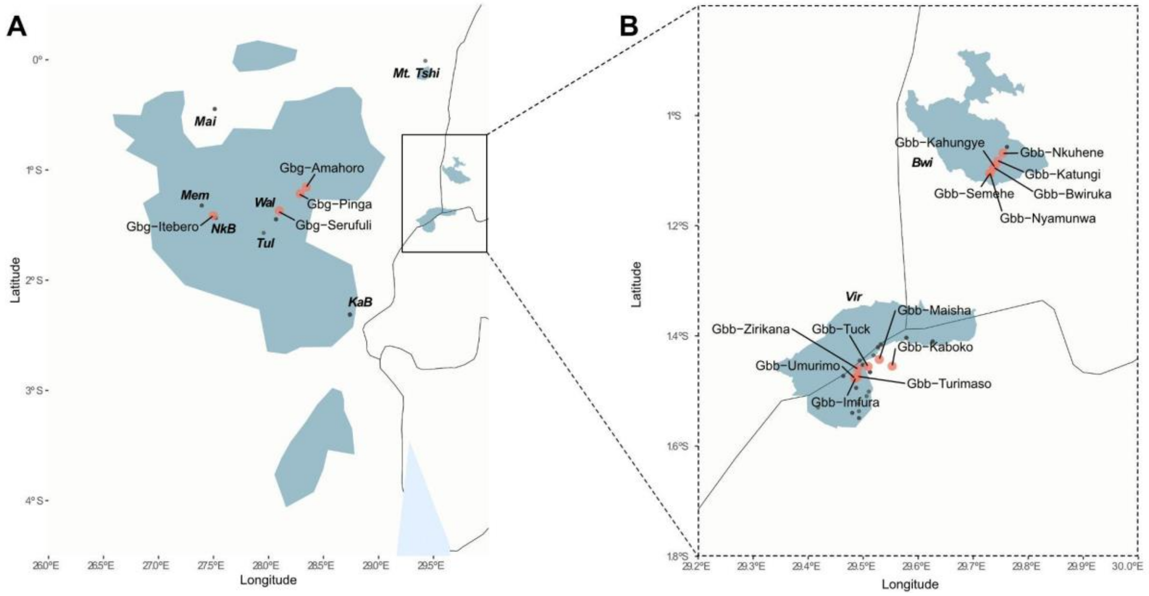


Figure 91 Supplementary. Geolocalization of eastern lowland (A) and mountain gorilla (B) public genomes (Xue et al. 2015; Pawar et al. 2023; Prado-Martinez et al. 2013). Black points represent the locations used for predicting locations, and in red are the locations predicted.

Western lowland gorilla origin information was not available for the majority of public genomes (Prado-Martinez et al. 2013; Prado-Martinez et al. 2013b). In order to evaluate the geographic coordinates predicted from these samples, the available country information in the zoo records was used. More detailed information in the results and discussion section of the main manuscript and Figures 92 and 93 of Supplementary.

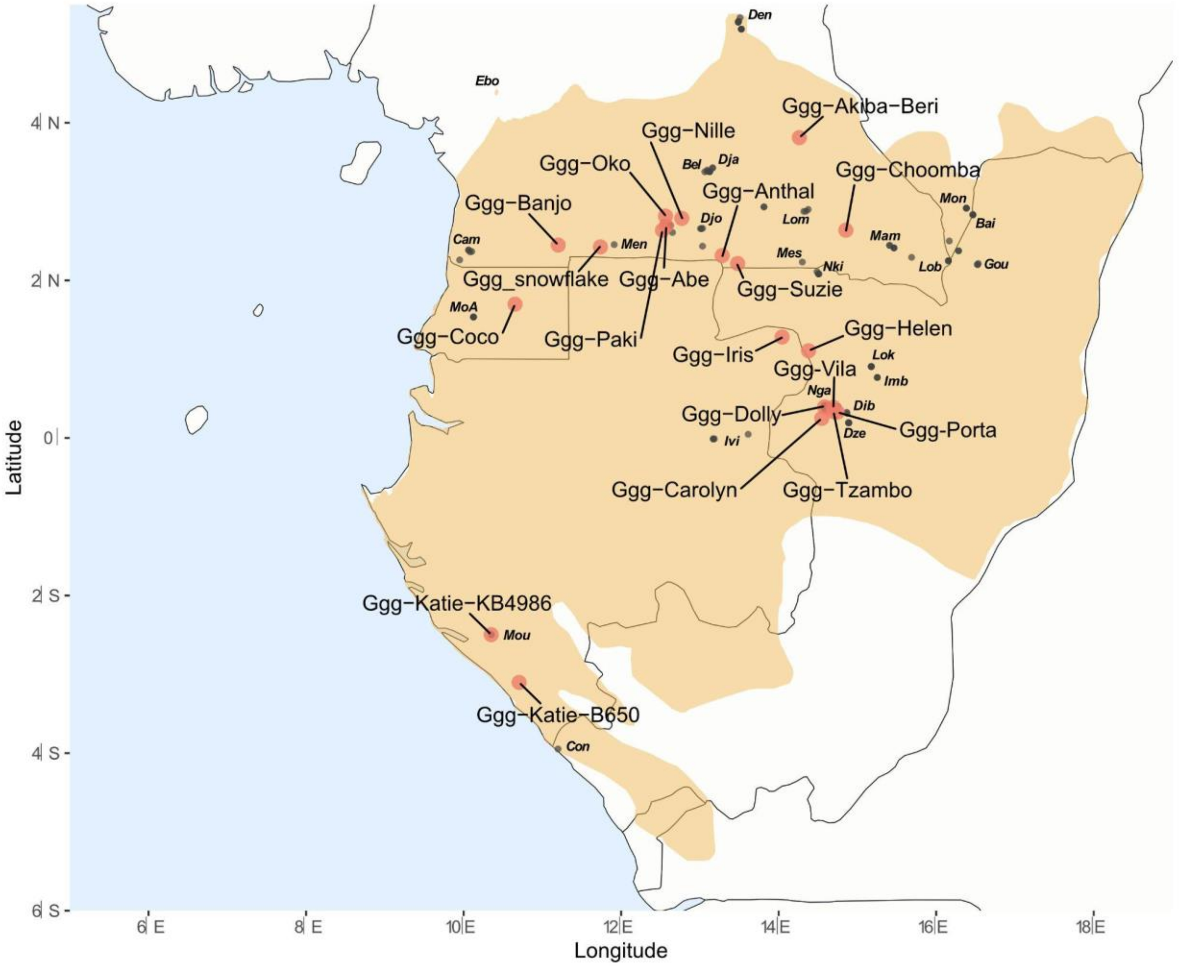


Figure 92 Supplementary. Geolocalization of western lowland gorilla public genomes (Prado-Martinez et al. 2013; Prado-Martinez et al. 2013b). Black points represent the locations used for predicting locations, and in red are the locations predicted.

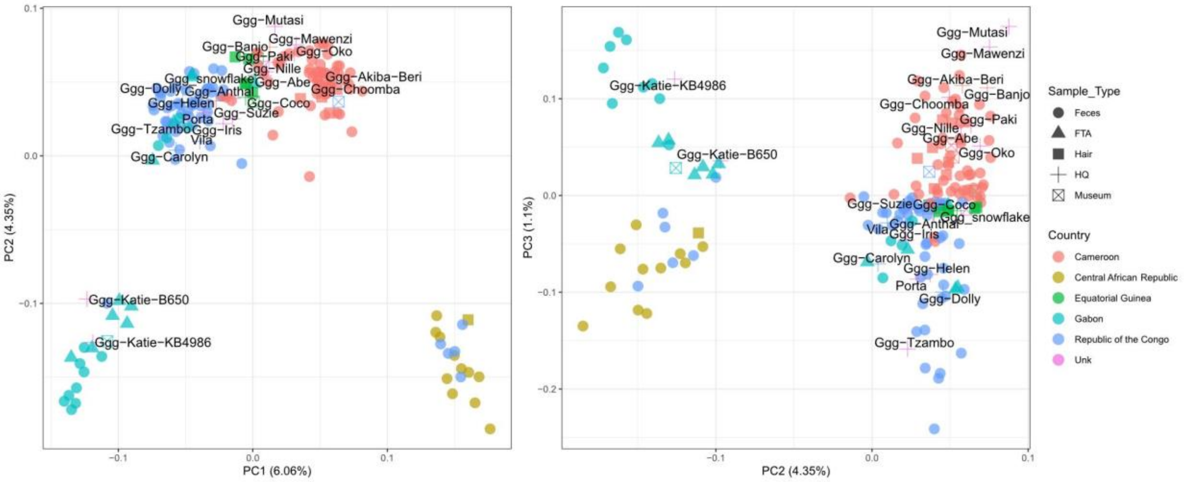


Figure 93 Supplementary. Principal Component Analysis western lowland gorillas, including Ebo forest sample and public genomes of the subspecies indicated with sample id labels (Prado-Martinez et al. 2013; Prado-Martinez et al. 2013b).

Finally, geographic coordinates for the Cross River gorilla Nyango were predicted between Boshi and Mbe areas, being closer to Boshi (Figure 94 Supplementary). This fact agrees with the previously published PCA using the same Cross River gorilla dataset (Alvarez-Estape et al. 2023).

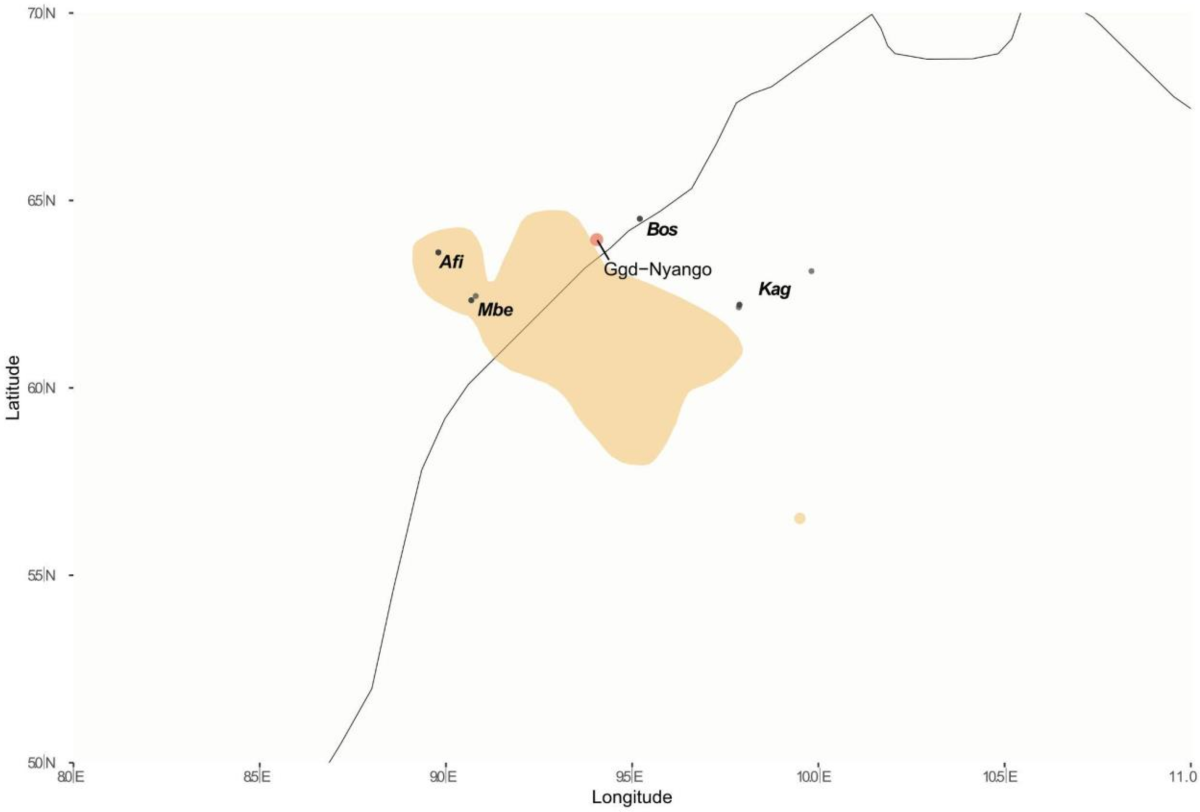


Figure 94 Supplementary. Geolocalization of Nyango (Cross River gorilla public genome) (Prado-Martinez et al. 2013). Black points represent the locations used for predicting locations, and in red are the locations predicted.

FTA cards and Museum sample geographic coordinates for the four subspecies were also predicted following the same approach. FTA card samples were obtained from rescued gorillas from illegal trafficking. The geographic coordinates were compared to the reported origin from where each individual was found. For some museum samples, the country of origin information was available for the majority of them, and was used to compare with our predicted geographic coordinates.

<u>WLG FTA and Museum samples:</u>

The country of origin for all FTA card samples was Gabon, in contrast to principal component analysis (Figure 37 Supplementary) and geographic coordinate prediction results (Figure 95B Supplementary). Three individuals (L4221A, L4222A and L4223A), which were reported to be rescued in the Port-Gentil area in Gabon, clustered in different groups in the PCA, resulting in different coordinate predictions. L4223A geographic coordinates were predicted in Equatorial

Guinea. In contrast, for the L4221A sample, the geographic coordinates were predicted in the south of Gabon and for the L4222A sample in the Odzala National Park. Two other FTA cards were also predicted to be from the Odzala National Park (L4220A and L4219A), but their rescued locations were Okondja and Mekambo, respectively. The rest of the rescued individuals were predicted to be from the South of Gabon. These individuals were rescued in Bakoumba (L4212A and L4213A), Gamba (L4214A), Haut-Ogooue (L4215A), Lambarene (L4217A) and Libreville (L4218A). Of these, two (L4218A and L4214A) were predicted to be close to Moukalaba-Doudau National Park and the others (L4212A, L4213A, L4217A and L4221A) from a southern area of the Republic of Congo, closer to Conkouati.

For 5 of the 6 western lowland gorilla museum samples, country origin information was available and was used to compare with our predicted geographic coordinates (Figure 95A Supplementary). The sample with unknown origin was labelled as a captive individual from 1965 (Frankfurt museum - AY4626), possibly captured from wild populations. For AY4643 and AY4642 country of origin was Equatorial Guinea. The predicted location of AY4642 was inside the same country, but for AY4643, the coordinates were in the same Cameroon area close to Equatorial Guinea, where Floquet's geographic coordinates were also predicted. AY4640 sample information origin (Gabon) also matched with the predicted location, south of Gabon area near Moukalaba-Doudau National Park. For the other four museum samples, the geographic coordinates were predicted within Cameroon. Of these, two were labelled to be from Cameroon (AY4629 and AY4765). The unique museum sample from the Republic of Congo (AY4618) was also grouped with all samples from Cameroon, indicating a mismatch with the origin information. For internal and original IDs of the museum samples, see Supplementary Table 3.

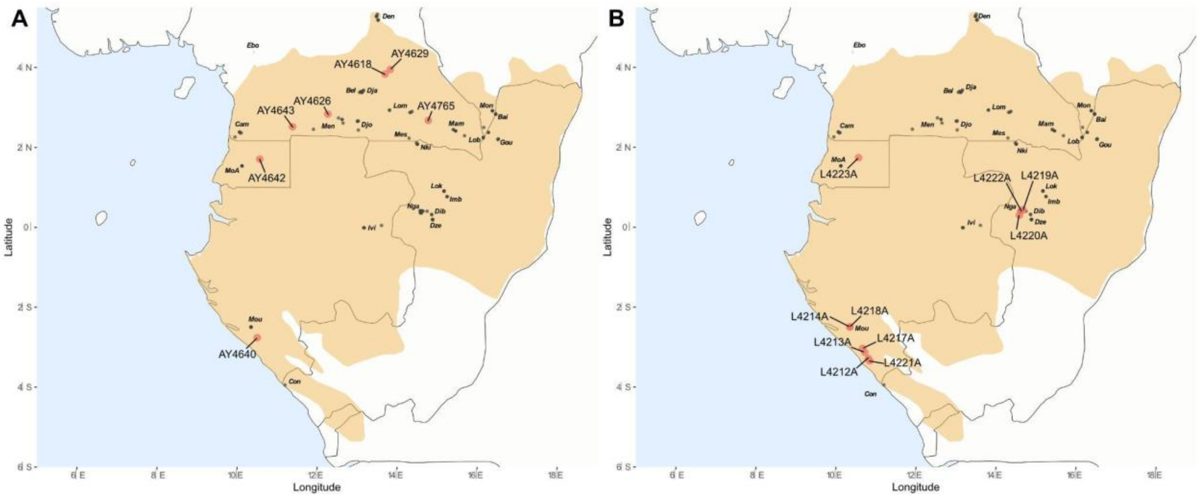


Figure 95 Supplementary. Geolocalization of newly sequenced western lowland gorilla museum (A) and FTA card samples (B). Black points represent the locations used for predicting locations, and in red are the locations predicted.

ELG Museum samples:

Three eastern lowland museum samples were collected and used in this geolocalization analysis (Figure 99A Supplementary). The geographic coordinates for the AY4638 sample were predicted within the Nkuba Conservation Area. The geographic coordinates of AY4771 were closer to the Membe area, while AY4767's geographic coordinates were predicted at a mid-point between Membele and Wallikale. This could be explained by missing samples representing sites from the true origin area.

CRG Museum sample:

The unique Cross River museum sample was predicted to be between Afi, Boshi and Bos areas (Figure 96B Supplementary). However, the museum metadata indicated Mamfe as the origin area, which is an isolated area within the Cross River distribution range. More samples from different missing Cross River populations will be needed to better represent the genetic differences and population dynamics, which will allow us to better predict geographic coordinates from populations for which genetic information is not available.

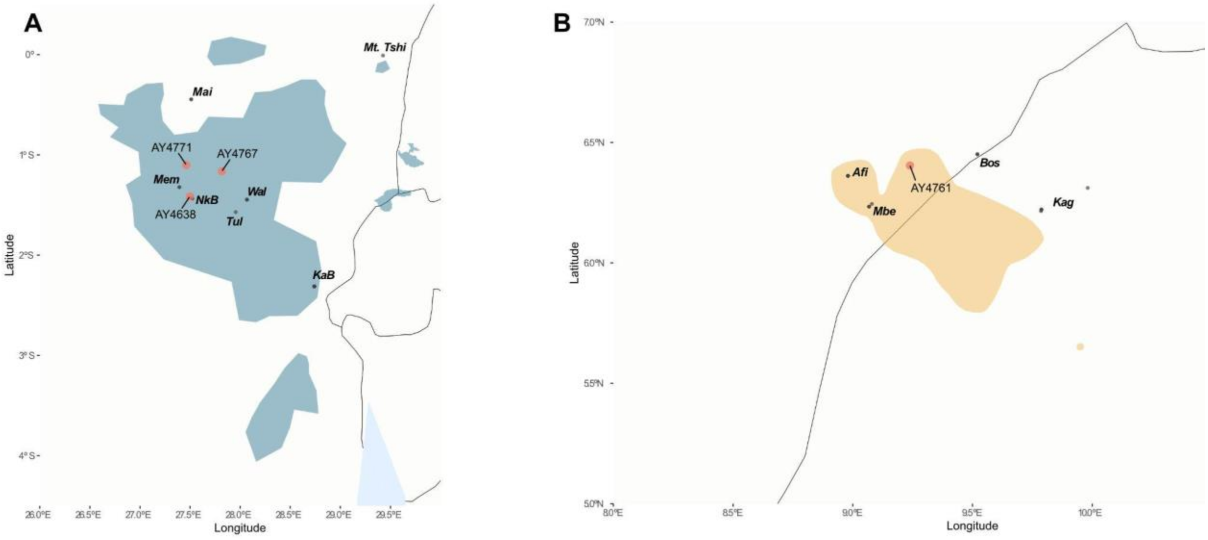


Figure 96 Supplementary. Geolocalization of newly sequenced eastern lowland (A) and Cross River gorillas (B) museum samples (B). Black points represent the locations used for predicting locations, and in red are the locations predicted.

### References


Alvarez-Estape, Marina, Harvinder Pawar, Claudia Fontsere, Amber E. Trujillo, Jessica L. Gunson, Richard A. Bergl, Magdalena Bermejo, et al. 'Past Connectivity but Recent Inbreeding in Cross River Gorillas Determined Using Whole Genomes from Single Hairs'. *Genes* 14, no. 3 (18 March 2023): 743. https://doi.org/10.3390/genes14030743.

Fontsere, Claudia, Marina Alvarez-Estape, Jack Lester, Mimi Arandjelovic, Martin Kuhlwilm, Paula Dieguez, Anthony Agbor, et al. 'Maximizing the Acquisition of Unique Reads in Noninvasive Capture Sequencing Experiments'. *Molecular Ecology Resources* 21, no. 3 (April 2021): 745–61. https://doi.org/10.1111/1755-0998.13300.

Hernández-Rodríguez, Jessica, Javier Fernandez-Gonzalez, Cristina F. Evans, Ines Lorente-Galdos, Cristina Wilson, Antonio F. Garcia-Perez, Tomas Marques-Bonet, and Arcadi Navarro. 2018. "The Impact of Endogenous Content, Replicates and Pooling on Genome Capture from Faecal Samples." Molecular Ecology Resources 18 (2): 319–33. https://doi.org/10.1111/1755-0998.12728.

Maeda, H. et al. "Unlocking the Potential of Animal Hair Shafts for Genomic Studies: A Comprehensive Evaluation of DNA Quality." Biology (MDPI), vol. 14, no. 4, 2023, article 353.

Plumptre, Andrew J., Sarah Nixon, David Caillaud, Jennifer S. Hall, Joseph A. Hart, Radar Nishuli, and Elizabeth A. Williamson. 2016. Gorilla beringei ssp. graueri. The IUCN Red List of Threatened Species 2016: e.T39995A102328430. https://doi.org/10.2305/IUCN.UK.2016-2.RLTS.T39995A17989838.en researchgate.net+1en.wikipedia.org+1

Prado-Martinez, Javier, Peter H. Sudmant, Jeffrey M. Kidd, Heng Li, Joanna L. Kelley, Belen Lorente-Galdos, Krishna R. Veeramah, et al. 'Great Ape Genetic Diversity and Population History'. Nature 499, no. 7459 (July 2013): 471–75. https://doi.org/10.1038/nature12228.

Smith LM, Burgoyne LA. Collecting, archiving and processing DNA from wildlife samples using FTA® databasing paper. BMC Ecol. 2004;4(1):4. doi:10.1186/1472-6785-4-4

Thalmann, Olaf, et al. 2007. "The role of Pleistocene refugia and rivers in shaping gorilla genetic diversity in central Africa." Proceedings of the National Academy of Sciences 104(51): 20432–37.

UNEP-WCMC. 2025. Protected Area Profile for Kahuzi-Biega National Park from the World Database on Protected Areas. Available at ProtectedPlanet.net. Accessed June 2025.

van der Valk, Tom, Axel Jensen, Damien Caillaud, and Katerina Guschanski. 2024. “Comparative Genomic Analyses Provide New Insights into Evolutionary History and Conservation Genomics of Gorillas.” BMC Ecology and Evolution 24 (14).

Xue, Yali, Javier Prado-Martinez, Peter H. Sudmant, Vagheesh Narasimhan, Qasim Ayub, Michal Szpak, Peter Frandsen, et al. ‘Mountain Gorilla Genomes Reveal the Impact of Long-Term Population Decline and Inbreeding’. Science 348, no. 6231 (10 April 2015): 242–45. https://doi.org/10.1126/science.aaa3952.
